# Collagen fibril formation at the plasma membrane occurs independently from collagen secretion

**DOI:** 10.1101/2024.05.09.593302

**Authors:** Adam Pickard, Richa Garva, Antony Adamson, Ben C. Calverley, Anna Hoyle, Christina E. Hayward, David Spiller, Yinhui Lu, Nigel Hodson, Oriana Mandolfo, Kevin K. Kim, George Bou-Gharios, Joe Swift, Brian Bigger, Karl E. Kadler

## Abstract

Collagen fibrils are the primary supporting scaffold of vertebrate tissues but how they are assembled is unclear. Here, using CRISPR-tagging of type I collagen and SILAC labelling, we elucidate the cellular mechanism for the spatiotemporal assembly of collagen fibrils, in cultured fibroblasts. Our findings reveal multifaceted trafficking of collagen, including constitutive secretion, intracellular pooling, and plasma membrane-directed fibrillogenesis. Notably, we differentiate the processes of collagen secretion and fibril assembly and identify the crucial involvement of endocytosis in regulating fibril formation. By employing Col1a1 knockout fibroblasts we demonstrate the incorporation of exogenous collagen into nucleation sites at the plasma membrane through these recycling mechanisms. Our study sheds light on the assembly process and its regulation in health and disease. Mass spectrometry data are available via ProteomeXchange with identifier PXD036794.

## Introduction

Collagen is the most abundant structural protein in mammals where it accounts for approximately 25% of total body protein mass (Smejkal and Fitzgerald, 2017). It occurs mostly as *D*-periodic (where *D* ∼ 67 nm) fibrils that can be centimeters in length (Craig et al., 1989), range in diameter from ∼12 nm to ∼300 nm depending on tissue and stage of development (Parry et al., 1978), and provide sites of attachment for a wide range of macromolecules including signaling complexes and cell receptors (Wickstrom and Fassler, 2011). However, elucidating the complex process of collagen fibril assembly has remained challenging due to technical constraints. For example, individual fibrils are too narrow to be visualized by conventional light microscopy and electron microscopy provides information only on pre-formed fibrils. Studies *in vitro* using purified collagen in the absence of cells show that the fibrils form by self-assembly (Gross and Kirk, 1958; Kadler et al., 1987; Kadler et al., 1990) but the *in vivo* mechanisms that determine the site of fibril assembly (nucleation) and how fibrils elongate remain elusive. *In vivo* observations using electron microscopy of embryonic tendon and cornea reveal fibrils attached to plasma membranes (Birk and Trelstad, 1984; Birk and Trelstad, 1986; Canty et al., 2004; Kalson et al., 2013; Starborg et al., 2013; Trelstad and Hayashi, 1979; Young et al., 2014). Our studies described here were motivated by a need to record, in real time, fibril assembly in the presence of cells. We aimed at capturing time-lapse images of collagen molecules as they traffic through the cell and subsequently appear as fibrils in the extracellular space.

In this study, we leverage CRISPR/Cas9 gene editing to tag collagen precursor proteins with photoswitchable and bioluminescent markers, and used advanced imaging techniques to visualize fibril assembly at the plasma membrane. We go further to reveal a role for endocytic collagen recycling in the early events of fibril nucleation. We discuss our findings in the context of tissue engineering and therapeutic interventions targeting collagen-related disorders.

## Results

### Engineering of Tagged Procollagen-I in NIH3T3 Cells

In this study, we employed CRISPR-Cas9 technology to engineer photoswitchable Dendra2 or nanoluciferase (Nluc) downstream of the signal peptide sequence of proa2(I) in NIH3T3 cells (shown schematically in **Fig. 1A**), resulting in the expression of tagged PCI (procollagen I) protein. Our quantification analysis revealed that fibroblasts secrete approximately 100,000 procollagen molecules per hour, equivalent to their entire procollagen content in approximately 2.5 hours (**Fig. S1**). Detection of Nluc tagged PCI in the medium within 5 minutes of changing the culture medium confirmed rapid secretion rates (**Fig. S1B)**, consistent with previous findings (Canty et al., 2004; Mirigian et al., 2014), highlighting the significant synthetic capacity of fibroblasts.

**Figure 1:**
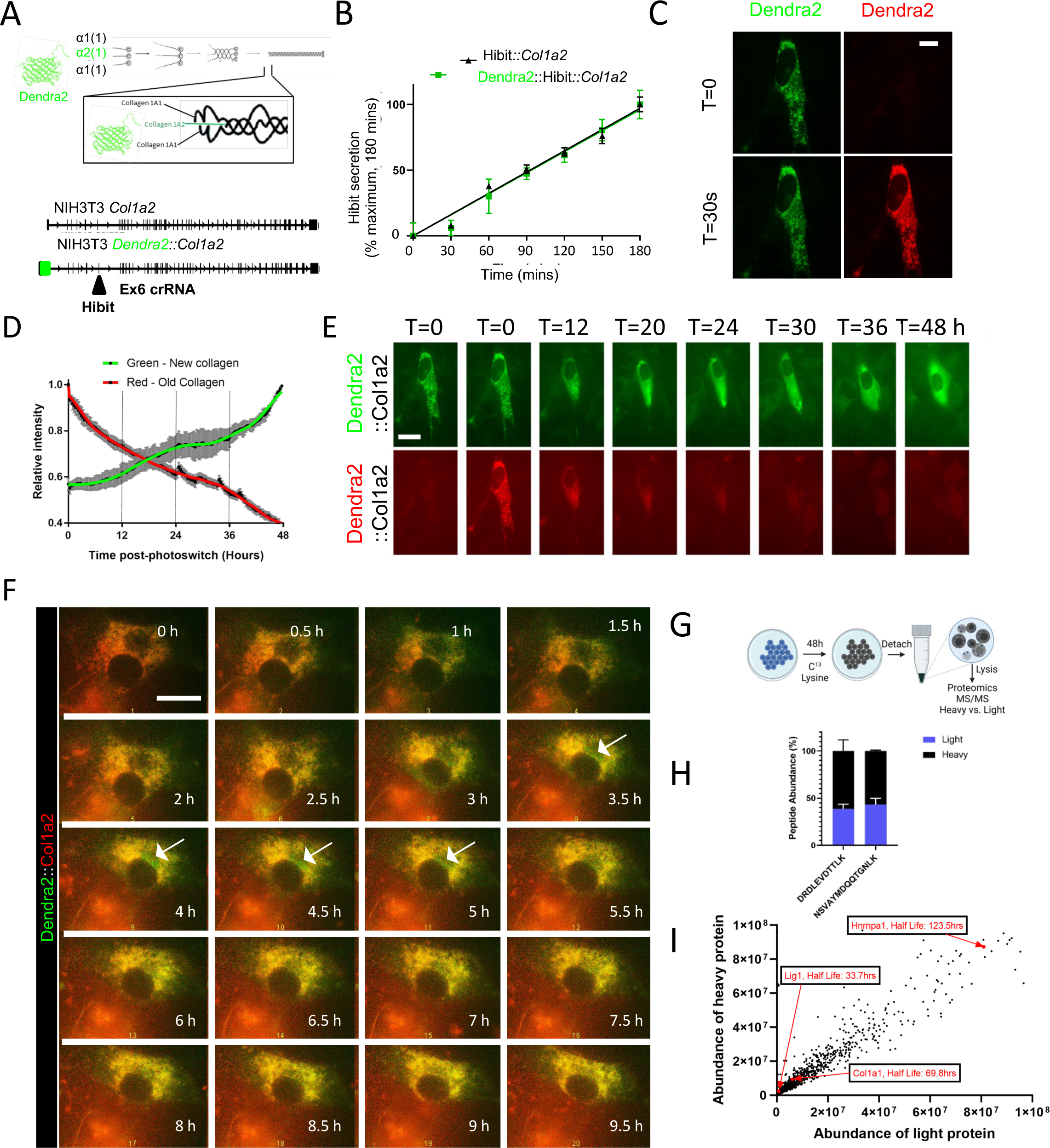
PCI secretion and collagen fibril assembly are separate processes. A) Schematic showing the site of integration of photoswitchable Dendra2 at the N-terminus of proa2(I) and location of Hibit tagging sites in Col1a2. B) Introduction of the split nanoluciferase sequence encoding HiBit was confirmed by detection of HiBit in the medium of edited cells. The rate of secretion for HiBit::Col1a2 and Dendra2::Hibit::Col1a2 were comparable. C) Photoswitching (green to red) of Dendra2-PCI after 30 second exposure to 405 nm light. D) Time-lapse microscopy showing continued synthesis of Dendra2-PCI (green) and secretion of Dendra2-PCI (red). E) Measurement of fluorescence intensity in the green and red channels shows synthesis (green channel) and secretion (red channel) of Dendra2-PCI over a period of 48 h. Note the 24 h rhythmic synthesis of Dendra2-PCI. F) Close up showing newly synthesized Dendra2-PCI (green) transiting through a pool of Dendra1-PCI, which remained detectable beyond 9 h post-photoswitching. G) Schematic showing SILAC labeling of unedited NIH3T3 cells. Cells were incubated with C^13^ lysine for 48 h before detachment, lysis and proteomic analysis. H) Quantification of both heavy and light labeled Col1a1-derived peptides demonstrate that not all PCI is secreted within 48 h. Peptide sequences shown are from the NC1 domain of proa1(I) mouse. N = 3 independent experiments. I) Global protein abundance in SILAC labeled NIH3T3 cells, the indicated protein half-lives were estimated by measuring the abundance of both heavy and light peptides for each protein as described in Methods.

### Tracking the Fate of Procollagen-I Molecules

To track the fate of procollagen molecules, we utilized Dendra2 to tag PCI at the N-terminus (**Fig. S1C-F**). Insertion of Dendra2 maintained allele responsiveness to pro-fibrotic stimuli (**Fig. S1G**) and did not alter secretion rates, as assessed by insertion of a HiBit tag into exon 6 using CRISPR-Cas9 (**Fig. 1B**). Leveraging the photoswitchable properties of Dendra2, we ‘switched’ the PCI (**Fig. 1C**) then followed the fate of existing PCI (which had been ‘switched’ to red) and the fate of newly-synthesized PCI (which was ‘green’). During 48 hours, pre-existing Dendra2-PCI (red) was secreted, while some of the newly-synthesized Dendra2-PCI (green channel) remained in the cell (**Fig. 1D**). The photoswitched Dendra2-PCI exhibited a half-life of approximately 24 hours (**Fig. 1E**). These results showed that not all PCI is rapidly secreted. Time-lapse analysis revealed that newly synthesized PCI initially located in puncta before becoming diffuse throughout the cell between 24 and 48 hours (**Video 1**). At higher magnifications, newly-synthesized Dendra2-PCI is observed transitioning through the Golgi apparatus whilst a large cellular pool of pre-existing Dendra2-PCI (yellow) was resident within the cell for over 9 hours (**Fig. 1F**).

### An intracellular pool of PCI confirmed by SILAC

To assess the residence time of PCI in cells, we utilized Stable Isotope Labeling by Amino Acids in Cell Culture (SILAC) labeling. This approach enabled measurement of protein turnover rates and secretion dynamics. Unedited NIH3T3 fibroblasts were cultured for 48 hours with C13-lysine followed by analysis of intracellular proteins using liquid chromatography coupled tandem mass spectrometry (LC-MS/MS) (**Fig. 1G**). The analysis revealed the presence of ‘heavy’ and ‘light’ peptides for 1380 proteins. Notably, both ‘heavy’ and ‘light’ Col1a1-encoding peptides were detected after 48 hours of labeling, with 39-43% of the peptides being ‘light’ (**Fig. 1H**), which supported the existence a longer-lived pool of intracellular PCI. The mass spectrometry proteomics data have been deposited to the ProteomeXchange Consortium via the PRIDE (Perez-Riverol et al., 2022) partner repository with the dataset identifier PXD036794 and 10.6019/PXD036794.

The ratio of ‘light’ to ‘heavy’ peptides facilitated the determination of protein half-lives (**Fig. 1I**). The presence of both ‘heavy’ and ‘light’ peptides derived from proa1(I) suggested the existence of a stable PCI pool (resulting in ‘light’ peptides) and a PCI pool synthesized during the 48-hour C13-lysine labeling (’heavy’ peptides). The ratio of light/heavy peptides allowed for the calculation of the half-life of ‘light’ proa1(I), which was found to be 69.8 hours.

Comparisons with other proteins revealed a wide range of half-lives, with Hnrnpa1 involved in pre-mRNA packaging exhibiting one of the longest half-lives (∼123 hours), while Lig1, a DNA ligase, had a half-life of ∼34 hours. This identification of a long-lived pool of PCI in unedited cells corroborates microscopy data obtained using Dendra2-edited cells. However, the discrepancy in half-life estimates between imaging and SILAC labeling suggested that fibroblasts may employ additional mechanisms to re-uptake secreted proteins (which we explored in experiments, described below), thereby lengthening the cellular residency of ‘light’ peptides.

### Collagen fibrils assemble at the plasma membrane

Dendra2-labelled collagen fibrils were not observed within the first 24 hours of imaging; however, with extended culture, Dendra2-positive fibrils became visible approximately 36 hours after cell plating (**Fig. 2A**). Multiple fibrils appeared simultaneously and grew in length over the next 12 hours (**Video 2**). This time lag in fibril formation aligns with previous findings using fixed samples (Chang et al., 2020; Kubow et al., 2015). Correlative Airyscan microscopy and atomic force microscopy (AFM) confirmed that the Dendra2-positive fibrils exhibited a *D*-period of 65 ± 7.4 nm, which was within the expected range for collagen fibrils (Ushiki, 2002) (**Fig. 2B, C**). These fibrils were confirmed as type I collagen using immunofluorescent detection (**Fig. S2**), and much like the effect of Dendra2 tagging on type I secretion, the location of the fluorescent protein did not impede fibril assembly. A key observation was that Dendra2-positive collagen fibrils were always associated with cells (**Fig. 3A, Video 3**); fibrils did not form independently of cells as might have been predicted from *in vitro* fibril assembly studies. Optical sectioning of the cultures and visualization of the cell body using an ER-targeted blue fluorescent protein (KDEL-BFP) revealed that Dendra2-positive fibrils deposited on the basal cell surface, near to but distinct from the BFP signal (**Fig. 3B, Video 4**).

**Figure 2:**
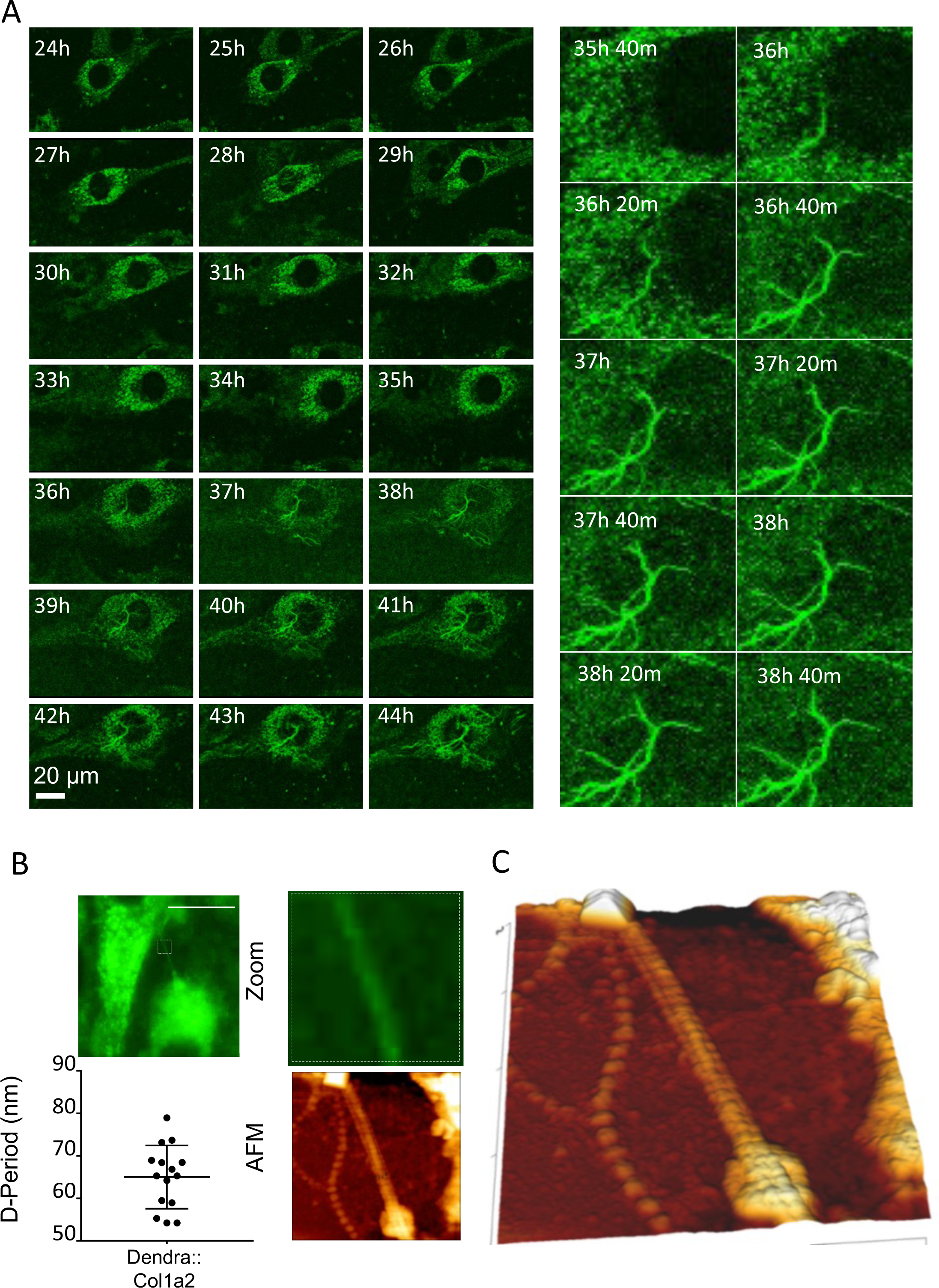
Collagen fibrillogenesis occurs on the plasma membrane. A) Time-lapse Airyscan microscopy of Dendra2::Col1a2 NIH3T3 cells. Dendra2 positive fibrillar structures were detected 36 h into culture. B) Dendra2::Col1a2 NIH3T3 cells were grown on correlative grid coverslips and imaged by widefield fluorescence microscopy. Samples were then fixed to perform AFM of the same region. Assessment of the periodicity of the Dendra2 positive fibril demonstrated a D-period of 65 nm. C) Zoom of the fibril shown in B. Scale bar 10 µm.

**Figure 3:**
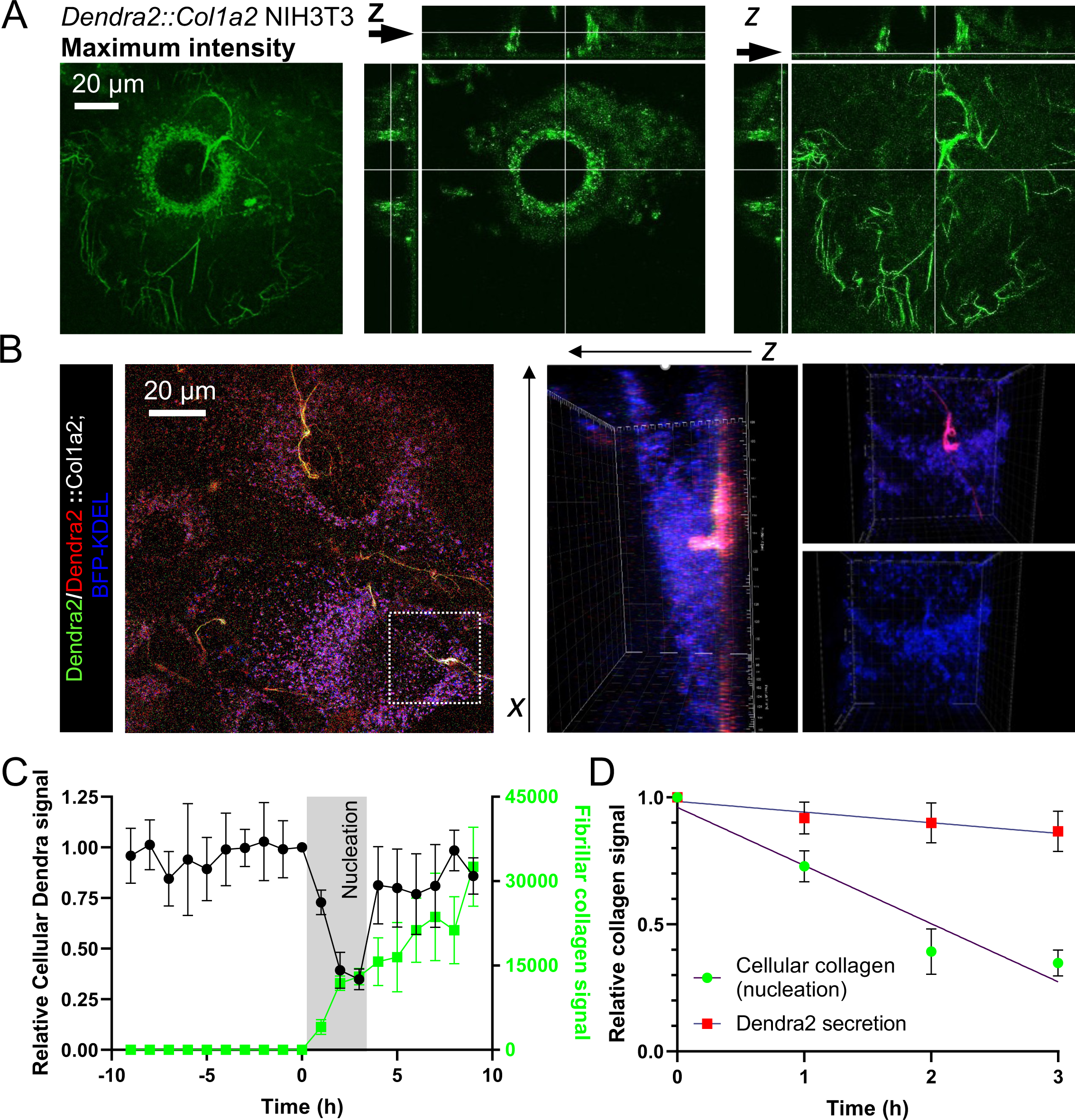
Collagen fibrils are formed between the basal surface of fibroblasts and the culture dish. A) Airyscan microscopy of Dendra2::Col1a2 NIH3T3 cells demonstrating that Dendra2 positive fibrils are formed within the boundary of the cell body. Images were captured after 48 h in culture. The maximum intensity projection of 51 different Z positions are shown on the left, middle panel shows the localization of Dendra2::Col1a2 in a single Z plane within the cell body, right panel shows that Dendra2 fibrils are abundant on the cells basal surface, a single Z plane is shown. Arrows indicate Z position of each image. **Video 2** shows all 51 Z positions. B) Expression of BFP-KDEL in Dendra2::Col1a2 NIH3T3 cells demonstrates that fibrils are formed on the basal surface of cells. A maximum intensity projection of 42 z positions is shown, left. The middle panel shows that photoswitched Dendra2 positive fibrils are beneath the cell body, however loops in collagen fibrils were observed extending into the cell body, a reconstruction is shown in **Video 3**. The, importantly at positions of where loops are engulfed by the cell the endoplasmic reticulum (BFP-KDEL) align with the fibrils, right. C) Quantification of cellular Dendra2::Col1a2 signal intensity in the cell body prior to the onset of fibril formation. The cellular Dendra2::Col1a2 signal drops dramatically during nucleation but then later recovers as fibrils elongate. Nucleation was set at the frame prior to the first detection of Dendra2 positive fibrils, this was designated as T=0, and the cellular Dendra2 signal was set as 1, background pixel intensity was set as 0. Data from five cells producing collagen fibrils are shown. Average data points are shown, n=5 individual cells, error bars represent SD. D) The loss of cellular Dendra2 signal observed in C demonstrates rapid release of cellular collagen at the onset of fibril formation, the rate of loss is compared to the rate of loss of photoswitched Dendra2 signal in the absence of fibril formation. Average data points are shown, n = 5 individual cells, error bars represent SD.

Insights into how the cellular pool of collagen contribute to the assembly process are also captured in these experiments; cellular Dendra2-PCI levels decreased rapidly during fibril nucleation, at a rate five times greater than we had observed when assessing secretion (**Fig. 3B, C**). Further examination of KDEL-BFP transfected cells demonstrates close proximity of the ER with the extracellular collagen fibril, with the ER arranged aligned to the forming collagen fibrils suggesting these sites may contribute to the rapid secretion. We also observed looped fibrillar structures engulfed by the cell body, and were often observed to be pulled by migrating cells. These images reinforce the conclusion that collagen fibril assembly is under strict cellular control and involves the plasma membrane in close association with the secretion machinery.

### Fibril nucleation is not dependent on classical collagen secretion

To generate further understanding, we employed a bioreactor to apply a constant flow of culture medium over the apical surfaces of CRISPR edited NIH3T3 cells and thereby maintain a low extracellular concentration of collagen (**Fig. 4A**). Applying a constant flow rate (0.05 mL/minute) throughout culturing significantly reduced extracellular media collagen levels by 99.5% when compared to static conditions (**Fig. 4B-C**). Decellularisation of the cell monolayer maintained under constant flow showed reduced cellular levels of Nluc (**Fig. 4D**) and reduced incorporation of Nluc into the matrix (**Fig. 4E**). Fibril assembly, assessed by microscopy, of Dendra2::Col1a2 NIH3T3 cells showed that the number of collagen fibrils per cell was not significantly affected by flow (**Fig. 4F, G**). The reduced cellular levels of Nluc in the face of unchanged collagen fibril numbers was explained when measurements of Dendra2 positive fibrils showed significantly shorter fibrils when cultured under flow. Together these results demonstrate that secreted collagen contributes to fibril growth but that nucleation is controlled and driven by cells.

**Figure 4:**
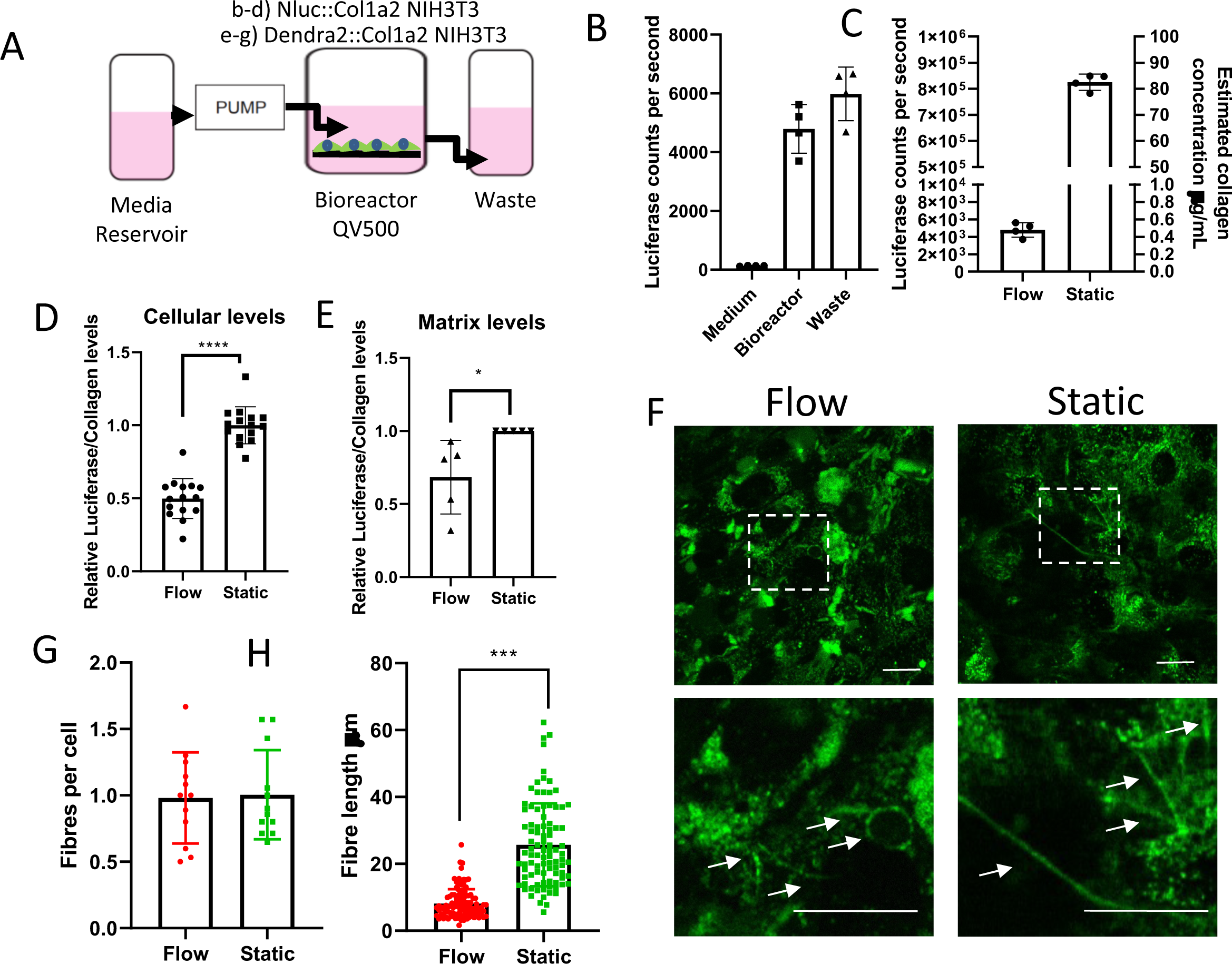
Rapidly secreted collagen contributes to fibril growth but not fibril nucleation. A) Schematic of the bioreactor used to study the effects of flow on fibril formation. B) To validate the flow system Nluc::Col1a2 NIH3T3 cells were grown under flow. Nluc activity was assessed in the medium within the Bioreactor housing cells seeded on glass coverslips. The activity was also measured in the medium within the waste outflow (n=4 independent experiments, error bars show SD). C) Comparison of Nluc activity within the medium of Nluc::Col1a2 NIH3T3 cells grown under flow conditions or under static ‘normal’ culture conditions for 72 h. Comparison of Nluc activity in the medium was compared to recombinant Nluc protein to estimate the concentration of collagen within the medium as described previously (Calverley et al., 2020). D) After 72 h culture cells were removed from the bioreactor, coverslips were decellularized, measurements of cellular Nluc activity was measured in triplicate (n=4 independent experiments, error bars show SD). E) After decellularization, Nluc activity in the deposited matrix were also assessed. N=5 independent experiments, error bars show SD. F) In parallel experiments the deposition of collagen fibrils by Dendra2::Col1a2 NIH3T3 cells were assessed under flow or static conditions. Airyscan microscopy of live cells after 72 hours culture identified Dendra2 positive fibrils. Scale bar represents 20 µm) G) The number of deposited collagen fibrils were quantified per cell, n=3 fields per view from 4 independent samples. H) The length of individual Dendra2 collagen fibrils were assessed using Image J image analysis software across all samples identified in G.

Having established that fibril nucleation is under cellular control, we next sought to understand how secreted collagen contributes to fibril growth. This was assessed in two ways, i) elimination of collagen nucleation sites and ii) addition of exogenous collagen. First, to eliminate fibril nucleation sites we crossed the Col1a1fl/fl mouse (Li et al., 2017; Yang et al., 2013) with the Col1a2-CreERT(2) mouse (Li et al., 2017; Yang et al., 2013) and isolated tendon fibroblasts. These primary fibroblasts were then incubated with tamoxifen to stop endogenous full-length collagen production. Immunofluorescence detection showed no type I collagen fibrils after tamoxifen treatment (**Fig. 5A**). When these cells were treated with conditioned medium from Dendra2:Col1a2-edited NIH3T3 cells, tagged collagen was observed intracellularly and was assembled into Dendra2-PCI into fibrils (**Fig. 5B**). This observation suggested that Dendra2:Col1a1 collagen was endocytosed by the cells and repurposed into fibrils. The endocytosis of collagen was confirmed by addition of Nluc-PCI conditioned media to NIH3T3 cells (**Fig. 5C**), which was found to be an active process unlike the uptake of the untagged Nluc enzyme (**Fig. S3**). Uptake of Nluc-PCI was identified to be dynamin dependent, evidenced by Dyngo treatment (**Fig. 5D**). The observation that endocytosed collagen contributes to fibril growth was confirmed by addition of Cy3-labelled rat tail collagen to NIH3T3 cultures; not only did the cells uptake Cy3-labelled collagen as assessed by flow cytometry (**Fig. 5E**), they also assembled this into the extracellular matrix (**Fig. 5F**). Uptake was also confirmed using human lung fibroblasts and SAOS2 cells (**Fig. S3**). These data support our earlier findings that extracellular collagen is endocytosed, which likely contributes to the longer than anticipated half-life of cellular collagen identified by SILAC labelling. Furthermore, these data demonstrate that the uptake of collagen from the extracellular space contributes to fibril assembly. This process functions independently from the collagen synthesis pathway and provides new understanding of how the extracellular matrix is assembled.

**Figure 5:**
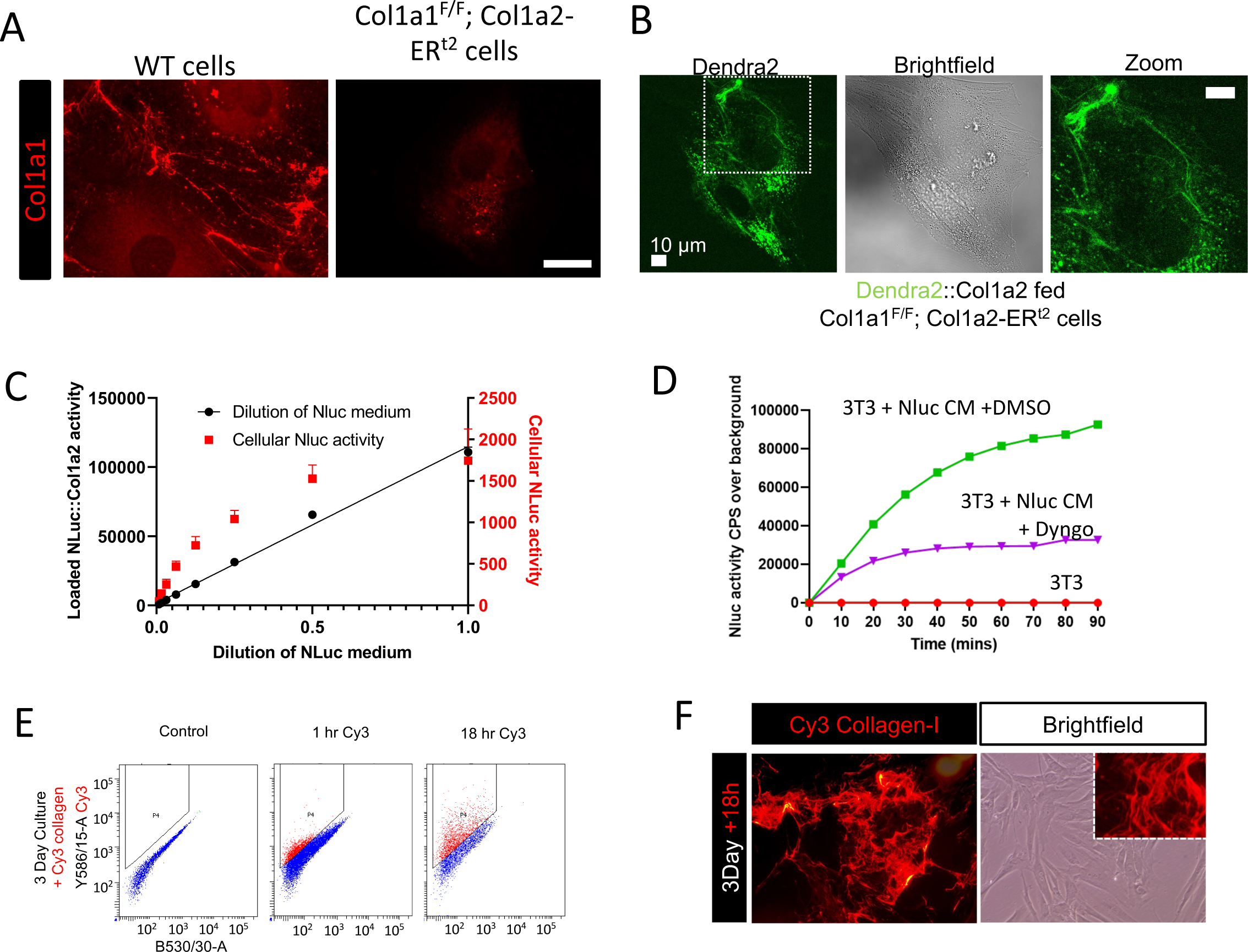
Endocytosed collagen contributes to fibril assembly. A) Deposition of type-I collagen fibrils in cultures of tamoxifen treated tendon fibroblasts from from wild-type and Col1a1^F/F^; Col1a2-ER^t2^ mice (Li et al., 2017; Yang et al., 2013). Type I collagen is detected by immunofluorescence. B) Tamoxifen treated fibroblasts from tendons of Col1a1^F/F^; Col1a2-ER^t2^ mice were incubated with conditioned medium from Dendra2::Col1a2 NIH3T3 for 72 h. The medium was replaced each day. Dendra2 positive fibrils were only detected associated with cells. Scale bar represents 10 µm. C) NIH3T3 fibroblasts were fed with conditioned medium from Nluc:Col1a2 NIH3T3 cells. The amount of Nluc activity taken up by tendon fibroblasts after 18 h was measured, demonstrating that cells were able to endocytose Nluc:Col1a2, however there was a limit to the amount of Nluc activity taken that can be endocytosed. N=3 independent experiments, error bars show SD. D) NIH3T3 cells were seeded in 35mm dishes and loaded with the live cell nanoluciferase substrate, endurazine for 2 h, cells were then treated with DMSO or 10 µM Dyngo4a. Luminescence was then measured every 10 minutes after addition of 1 mL of NLuc::Col1a2 conditioned medium cell metabolized substrate. Representative traces are shown. N=3 independent experiments. E) Cy3 labeled rat tail collagen (10 µg/mL) was added to cultures of human foreskin fibroblasts for 1 or 18 h cells were detached and analysed by flow cytometry demonstrating time dependent uptake by fibroblasts. Uptake was also observed in SAOS2 and lung fibroblasts Figure S3). F) Human foreskin fibroblasts incubated with 0.5 µg/mL Cy3 collagen for 18 h assembled labeled collagen into fibrillar structures.

### Collagen fibril formation blocked in the presence of monensin but not brefeldin A

To further examine how cells assemble collagen fibrils, we utilized Brefeldin A to inhibit ER-Golgi transport (Colanzi et al., 2013). Brefeldin A inhibited secretion of Nluc-PCI into the culture medium (**Fig. 6A**), but did not result in a reduction in collagen incorporation into the extracellular matrix (**Fig. 6B, C**). This finding was confirmed when cultures of Dendra2:Col1a2 cells were grown in the presence of brefeldin A, and indicated that pathways beyond the classical ER-Golgi secretory pathway might be involved in collagen fibril assembly. Next, we used Monensin to inhibit both ER-Golgi transport and transport to lysosomes (Stenseth and Thyberg, 1989). Monensin blocked fibril formation (**Fig. 6D-F**), which suggested that transport of collagen to the lysosome may be required for fibril formation. To explore whether the lysosome compartment has a role in fibril assembly, the inhibitors of lysosome function, bafilomycin A1 and chloroquine (reviewed by (Fedele and Proud, 2020)) were applied to unedited fibroblasts. At doses that did not alter cellular proliferation, both bafilomycin A1 and chloroquine significantly inhibited fibril formation and fibril length as assessed by immunofluorescence and by total collagen deposition assessed by hydroxyproline quantification (**Fig. S4**). When the combination of Bafilomycin and Brefeldin A were applied, fibril formation was completely blocked suggesting that ER-Golgi and lysosomal trafficking together are required for collagen fibril formation (**Fig. 6F**). This was also supported by more targeted interruption of only the ER-Golgi traffic, where knockdown of Syntaxin 5 (STX5) disrupted only procollagen secretion but had no impact on collagen incorporation into the matrix (**Fig. 6G-J, Fig. S5**).

**Figure 6:**
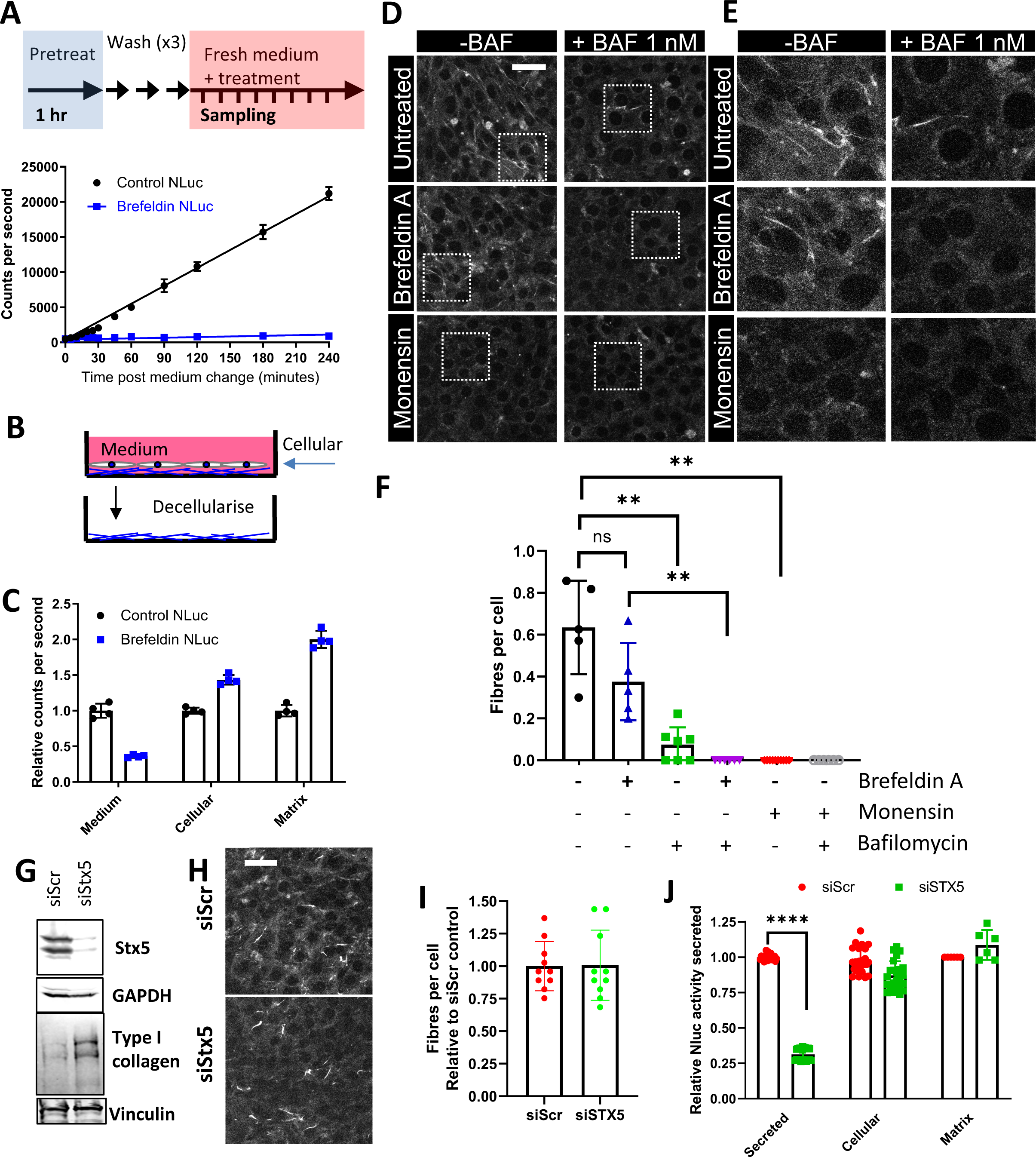
Non-classical PCI secretion feeds fibril assembly. A. Schematic for monitoring Nluc activity secretion from Nluc::Col1a2 NIH3T3 cells. Comparison of Nluc secretion rates in control cultures and cells treated with 100 nM brefeldin A identified robust reduction of Nluc activity secretion. B. Schematic to quantify the deposition of Nluc::Col1a2 into the extracellular matrix. C. Nluc::Col1a2 NIH3T3 cells were cultured with 100 nM brefeldin A for 72 h, Nluc activity levels in the conditioned medium, cellular fraction and matrix were assessed. N=3 independent experiments, N=4 technical repeats from a representative experiment are shown. Error bars represent standard deviation. D. Representative Airyscan confocal microscopy images of Dendra2::Col1a2 NIH3T3 cells cultured for 72 h in the presence of either 100 nM brefeldin A, or 1 µM monensin with or without addition of the lysosome proton pump inhibitor 1 nM bafilomycin. Dendra2 signals are shown to highlight deposited fibrils. Maximum intensity projections of 5 z planes are shown. Scale bar represents 50 µm. Boxes are enlarged in E. E. Enlargements of images within the regions highlighted in D. F. Quantification of Dendra2 fibril numbers per cell, n=5 independent experiments for brefeldin and monensin treatment, with an additional n=2 independent experiments for treatments including bafilomycin treatment. A minimum of 200 cells per condition, per experiment, were score. Error bars represent SEM. ** represents p<0.01, Student’s T-test, unpaired. G. Western blot of Syntaxin 5 (Stx5) knockdown with 100 pmol siRNA in NIH3T3 cells. N=3 independent experiments. H. Syntaxin 5 (Stx5) knockdown with 100 pmol siRNA in Dendra2::Col1a2 NIH3T3 cells were cultured for 72 h before live imaging of Dendra2 fibrils.. Maximum intensity projections of 5 z planes are shown. Scale bar represents 50 µm. I. Quantification of Dendra2 fibril numbers per cell in siRNA treated Dendra2::Col1a2 NIH3T3. n=2 independent. Error bars represent SD. J. Syntaxin 5 (Stx5) knockdown with 100 pmol siRNA in Nluc::Col1a2 NIH3T3 cells were cultured for 72 h, Nluc activity levels in the conditioned medium, cellular fraction and matrix were assessed. N=3 independent experiments, each with N=4 technical repeats are shown for conditioned medium and cellular fractions, or n=2 technical repeats for matrix measurements. Error bars represent standard deviation.

### Dendra2-PCI is transported in Lamp1-positive carriers

The possible involvement of lysosomal or lysosomal-like compartments in collagen fibril assembly motivated additional experiments. We used confocal microscopy with Airyscan detection to live image Dendra2:Col1a2 cells with the aim of visualizing pre-fibril forming events. Dendra2-PCI was conspicuous in being localized to a constellation of punctate compartments (**Fig. 7A**). These compartments were in close association with collagen fibrils, which also could be seen to form loop structures (**Fig. 7A, insert**). Such close association of collagen fibril-containing fibripositors with darkly-stained compartments (presumed to be lysosomal compartments) are frequently observed in electron microscope images of embryonic tendon (**Fig. 7B**) (Canty et al., 2004). In tendon fibroblasts cultured for 72 h, intracellular PCI localizes largely to the ER, as indicated by co-localization with the ER resident enzyme, protein disulfide isomerase (PDI) (**Fig. 7C**).

**Figure 7:**
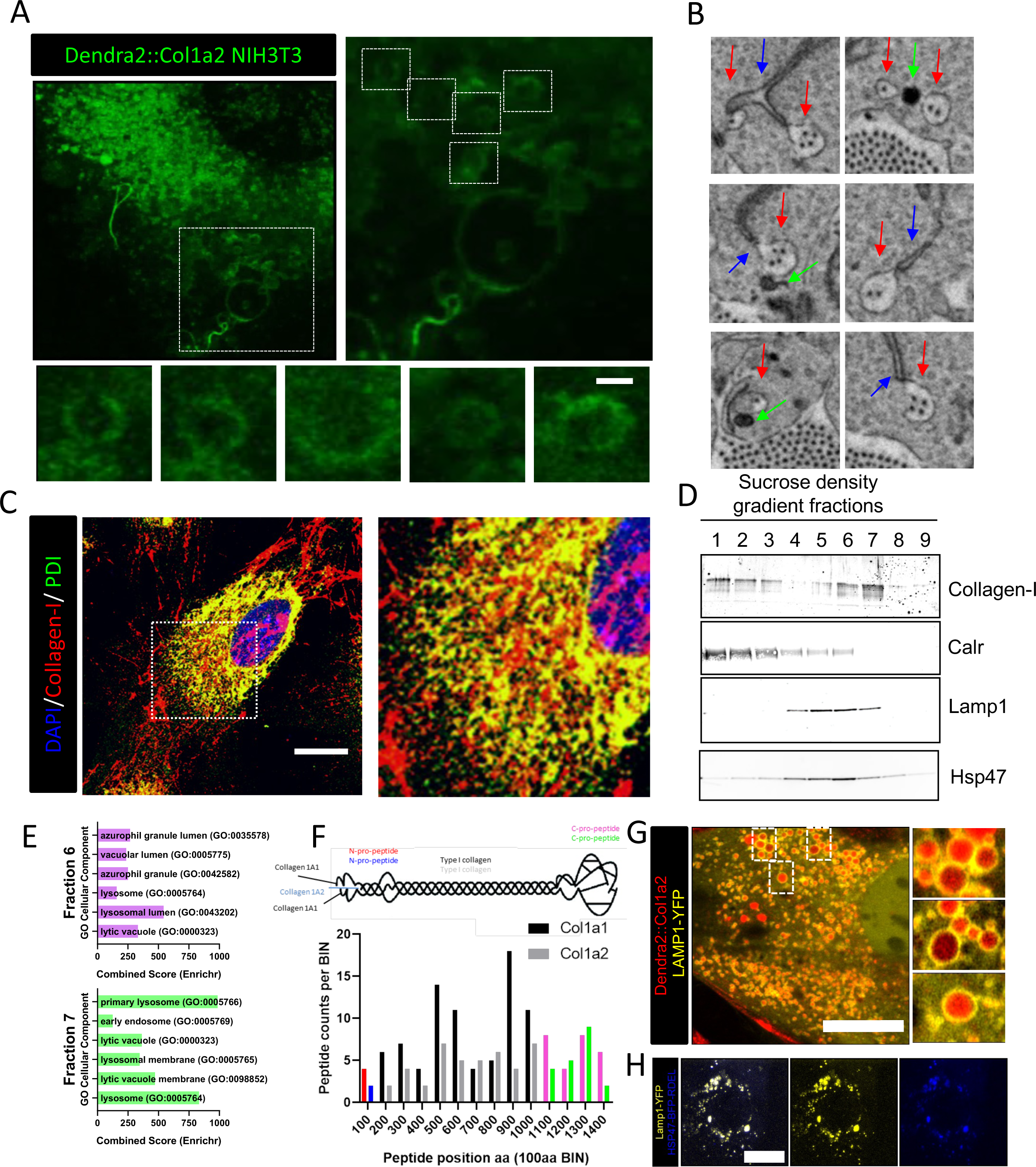
PCI localizes to LAMP1-positive compartments. A. Dendra2::Col1a2 NIH3T3 imaged using Airyscan microscopy after 18 h after culture, 1000x magnification, and 2.5x digital zoom, a maximum intensity projection is shown. Vesicles containing Dendra2 signal are observed at sites of fibril assembly. Dendra2 signal was observed arranged at the periphery of vesicles, scale bar represents 1 µm. Area outlined by the box is enlarged in the right hand panel. Small panels show looped structures containing Dendra2-positive collagen. B. Individual frames taken from Video 6 demonstrating that a single fibripositor (highlighted in the red box) is contacted numerous times by both endoplasmic reticulum and electron dense vesicles. C. Immunofluorescence imaging of type I collagen in mouse embryonic fibroblasts with the ER marker PDI. Scale bar represents 20 µm. D. Fractionation of NIH3T3 using a sucrose density gradient, type I collagen is co-resident with the ER protein Calreticulin (Calr) but also with the lysosomal protein Lamp1 and the collagen chaperone Hp47. E. Proteomic analysis of lysosomal fractions 6 and 7 identified significant enrichment of proteins identified by proteomic analysis of fractions 6 and 7 based on GO Cellular component terms. F. Col1a1 (Red and Magenta) and Col1a2 (Blue and Green) derived procollagen peptides were present within both fractions 6 and 7 suggesting that newly synthesized collagen transitions these compartments. G. Dendra2::Col1a2 NIH3T3 imaged using Airyscan microscopy 48 h after transfection with LAMP1-YFP. Images were recorded at 1000x magnification with 2.5x digital zoom. A maximum intensity projection of 31 images is shown. LAMP1-YFP vesicles (shown in green) containing photoswitched Dendra2 signals are observed. Three LAMP1-positive areas are enlarged. H. Live cell super resolution microscopy of NIH3T3 stably transduced with Hsp47-BFP-RDEL lentivirus and transfected with LAMP1-YFP. Scale bar, 20 µm.

There were also regions where the two proteins did not co-localize. In one set of experiments to determine the identity of these compartments we performed cellular fractionation followed by western blotting and LC-MS/MS protein identification. As expected PCI was identified in fractions that contained ER-resident protein (e.g. calreticulin, CALR) (**Fig. 7D**). PCI was also found in fractions that did not contain CALR (see fractions 5-7 in **Fig. 7D**). Some PCI-positive fractions contained the lysosome associated membrane protein (LAMP)-1, and these also contained Hsp47, which binds triple helical collagen (Ishikawa and Bachinger, 2013). These PCI-positive/CALR-negative fractions were analyzed by LC-MS/MS (**Table 1**). Pathway analysis of the proteins identified ‘endocytosis’ and ‘lysosome’ as the enriched pathways (**Fig. 7E**). Peptides originating from LAMP1 and LAMP2 as well as core components of the retromer complex (Vps35, Vps26, and Vps29) and the sorting nexins SNX1 and SNX6 were present in these PCI-containing fractions (**Table 1**). Retromer is responsible for the recycling of transmembrane proteins to the cell surface or the trans-Golgi network (TGN) and prevents their degradation by the lysosomal system (Bonifacino and Hurley, 2008). Within these fractions, peptides for both proa1(I) and proa2(I) were identified including peptides from the N- and C-propeptides. Further analysis of the proteins within these fractions identified 35 of the known collagen-interacting proteins (Doan et al., 2019). These included Hsp47 and PPIB, which catalyzes peptidyl proline isomerization and the rate-limited step in folding of the collagen triple helix. Similarly, Colgalt1, which transfers beta-galactose to the hydroxylysine residues on collagen, was also present. The collagen crosslinking enzymes, PLOD1-3, were also readily detected (**Table 1**). Taken together, the presence of these proteins suggests that triple helical PCI molecules and the procollagen folding machinery are present at the sites of collagen fibril formation.

**Table 1:**
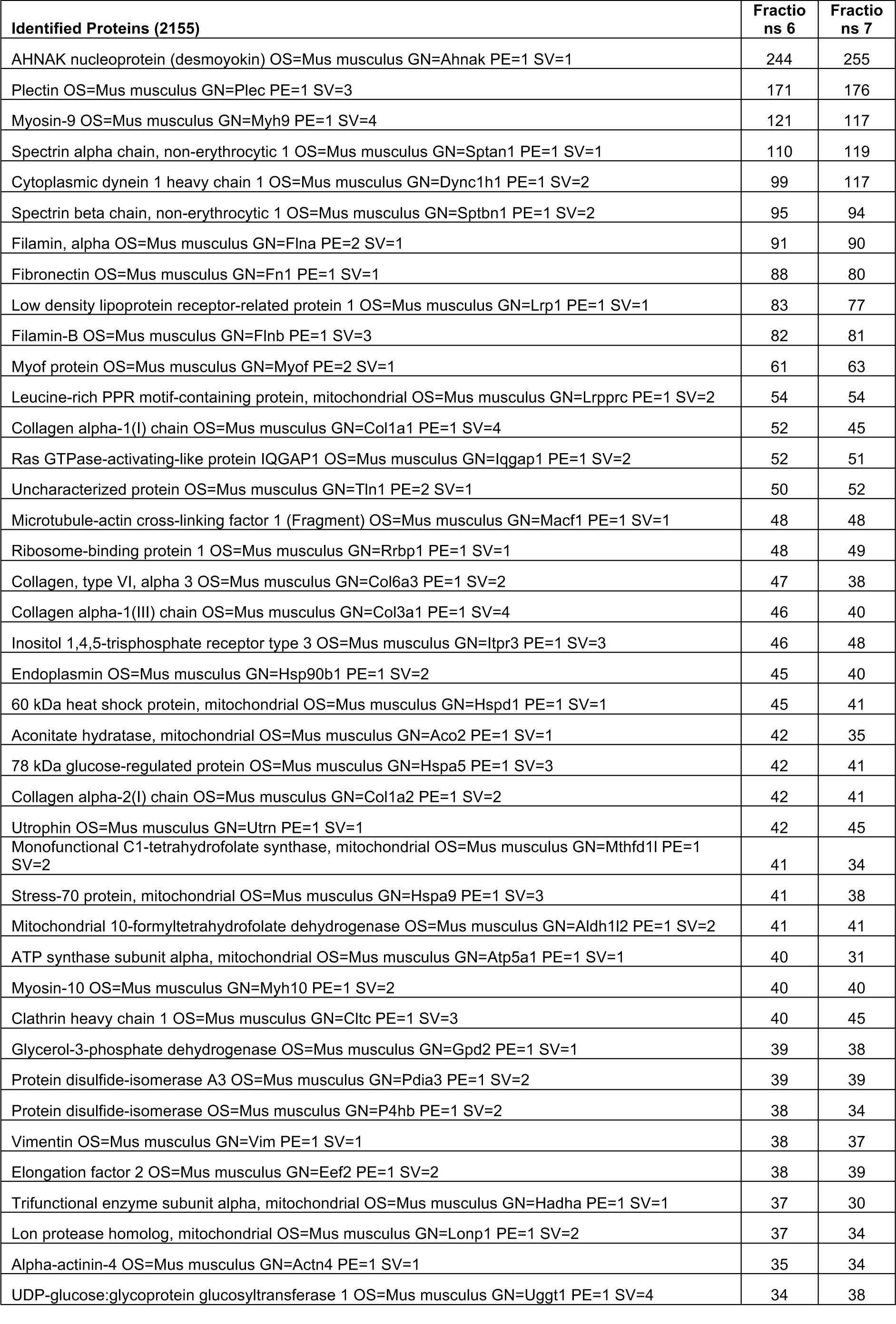
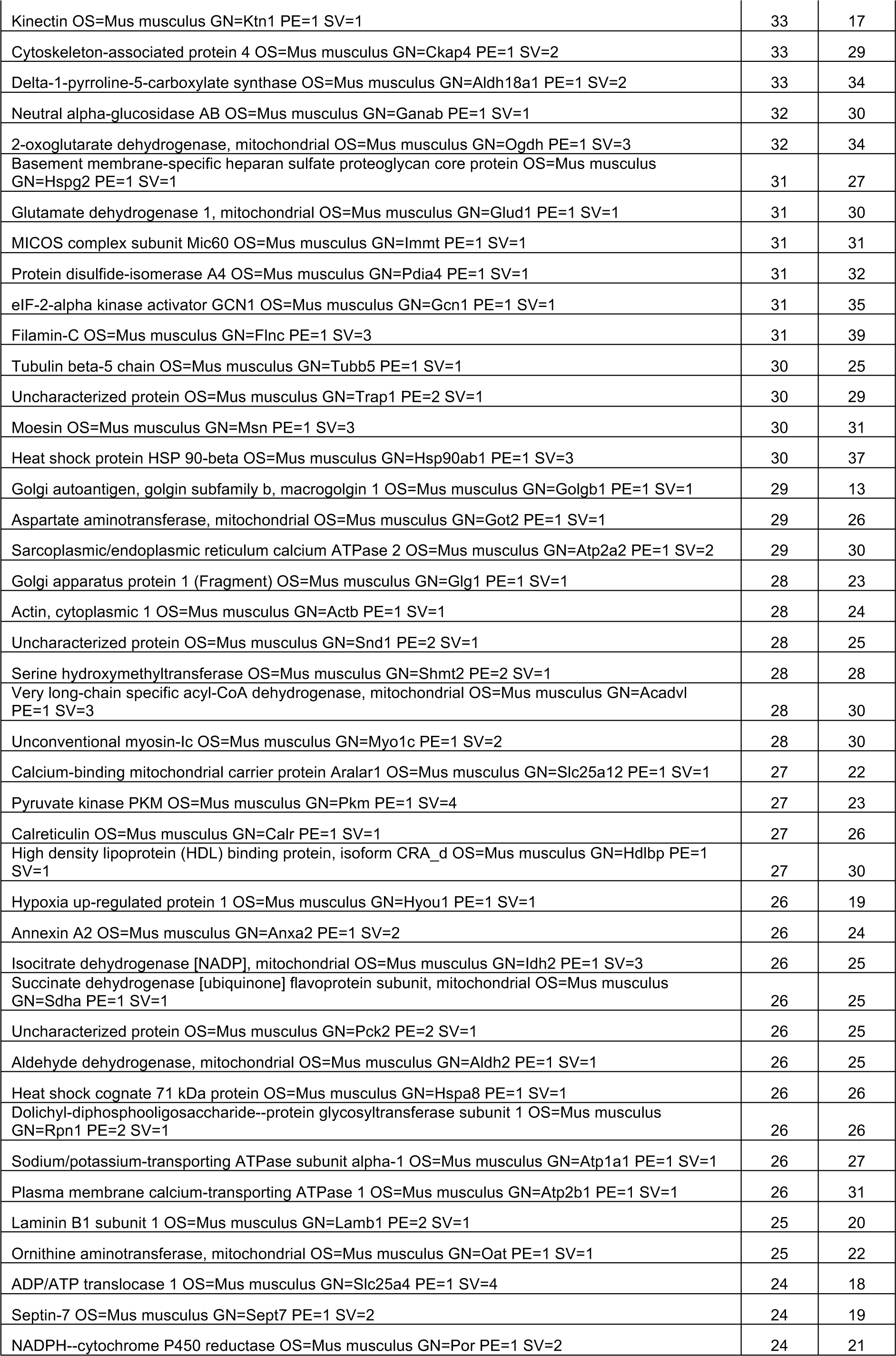
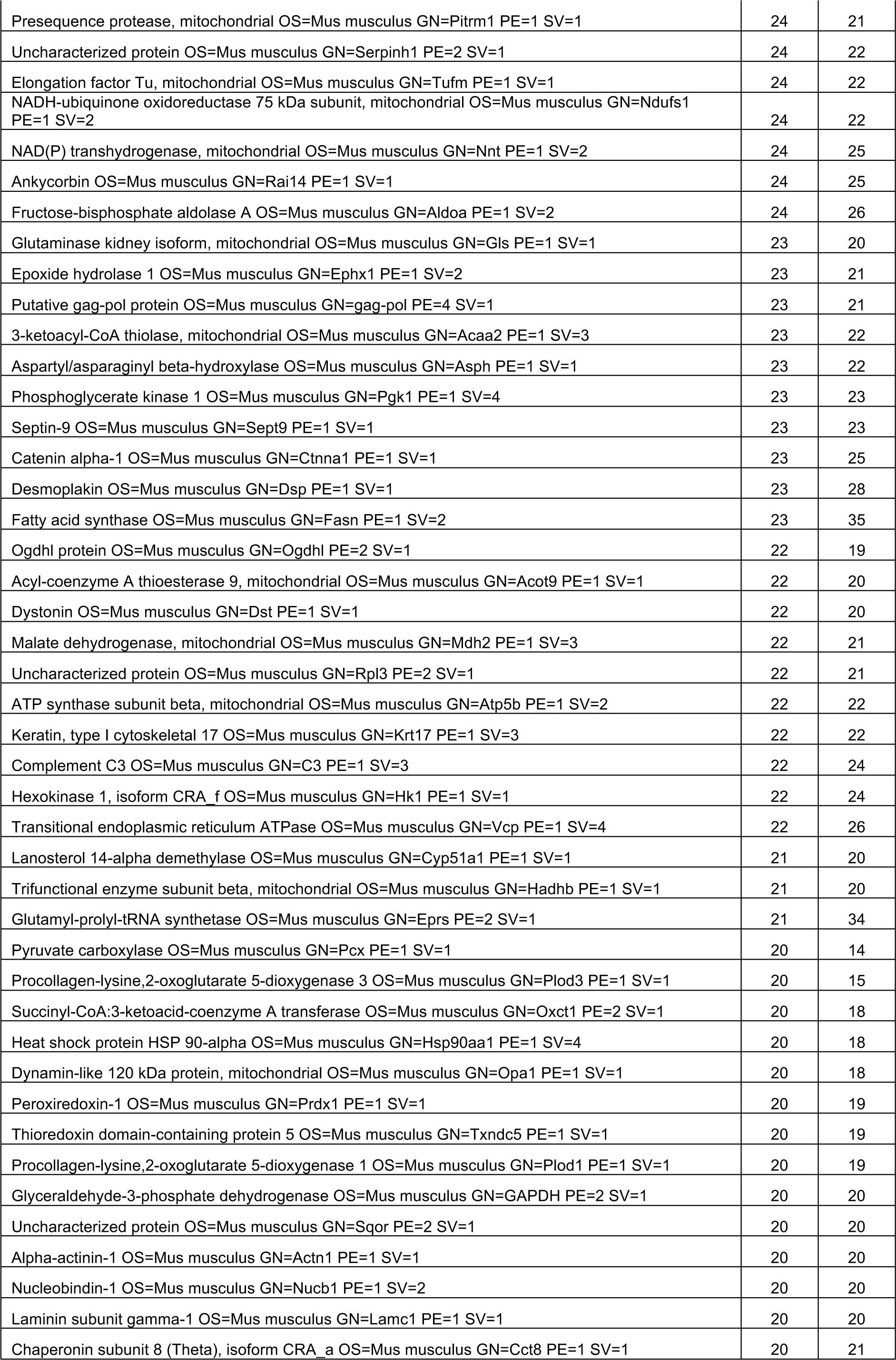
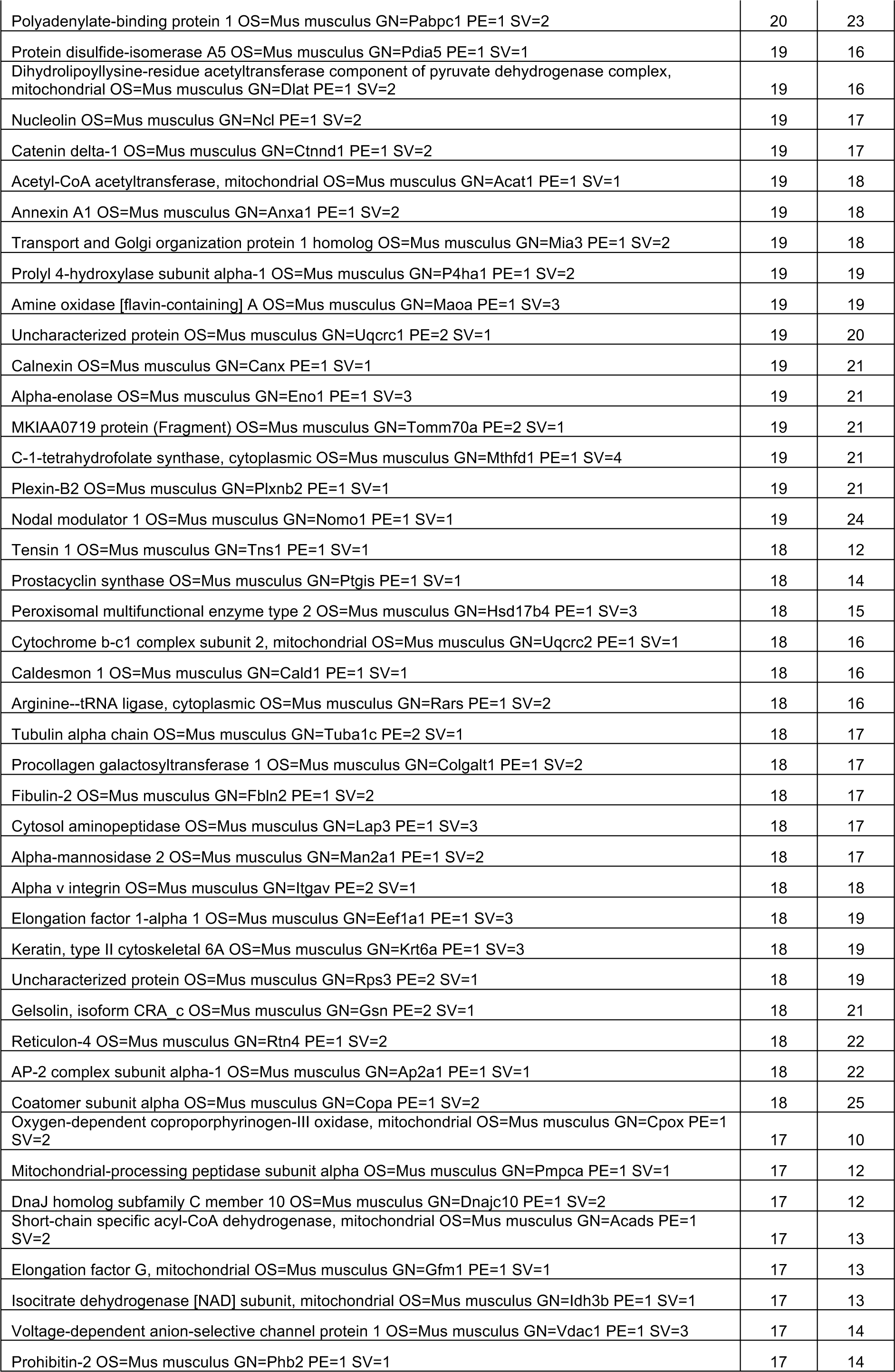
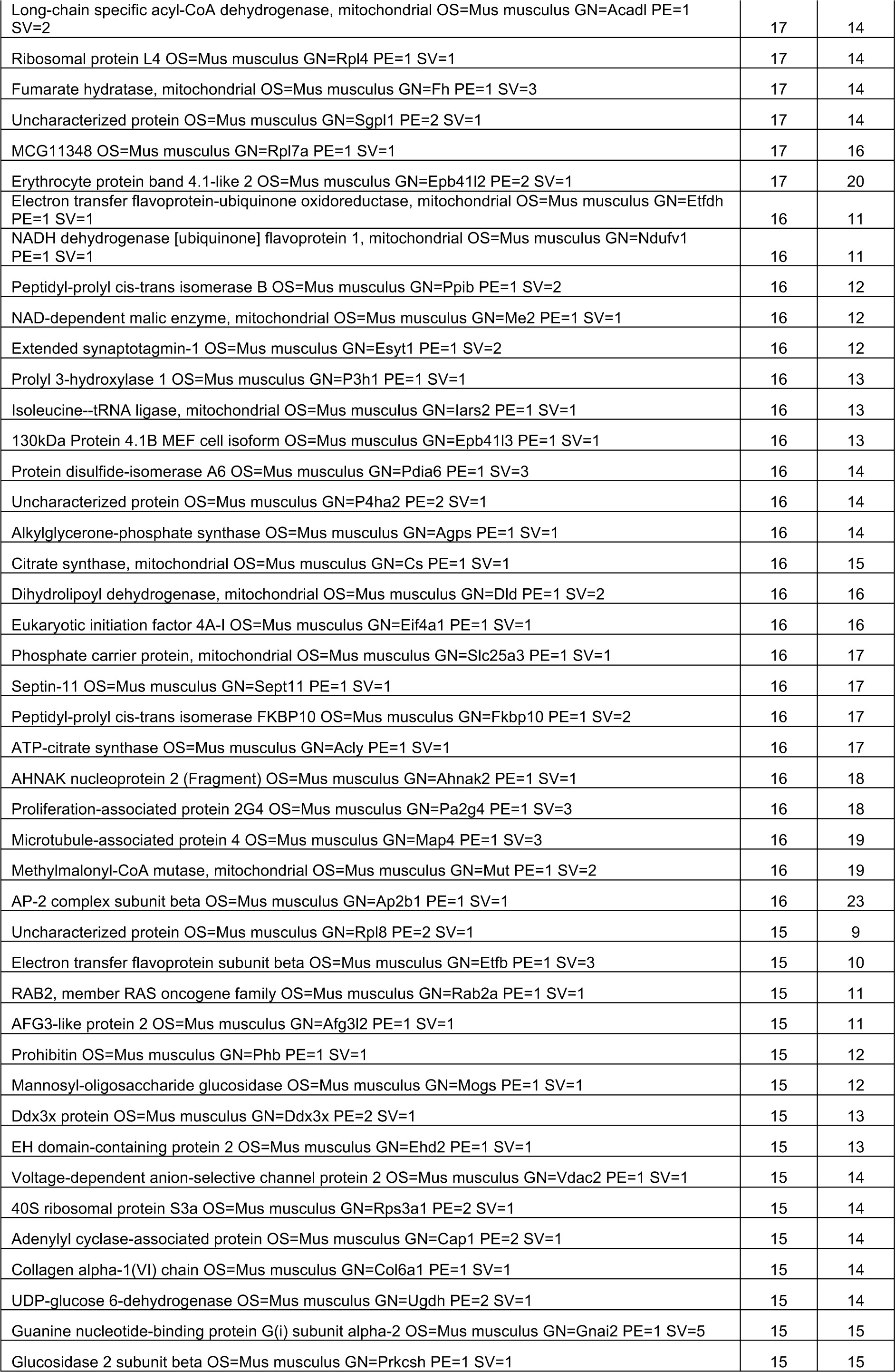
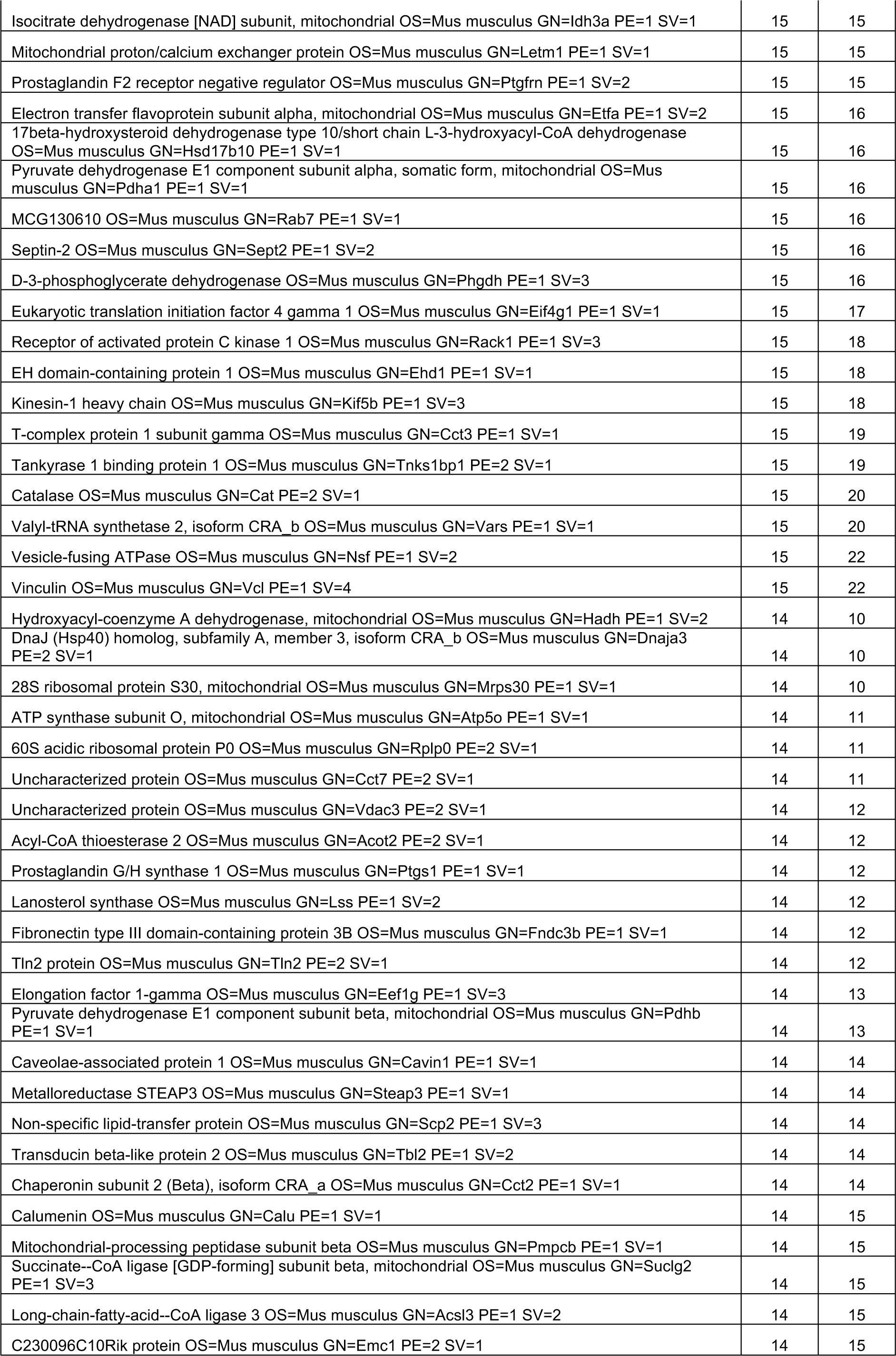
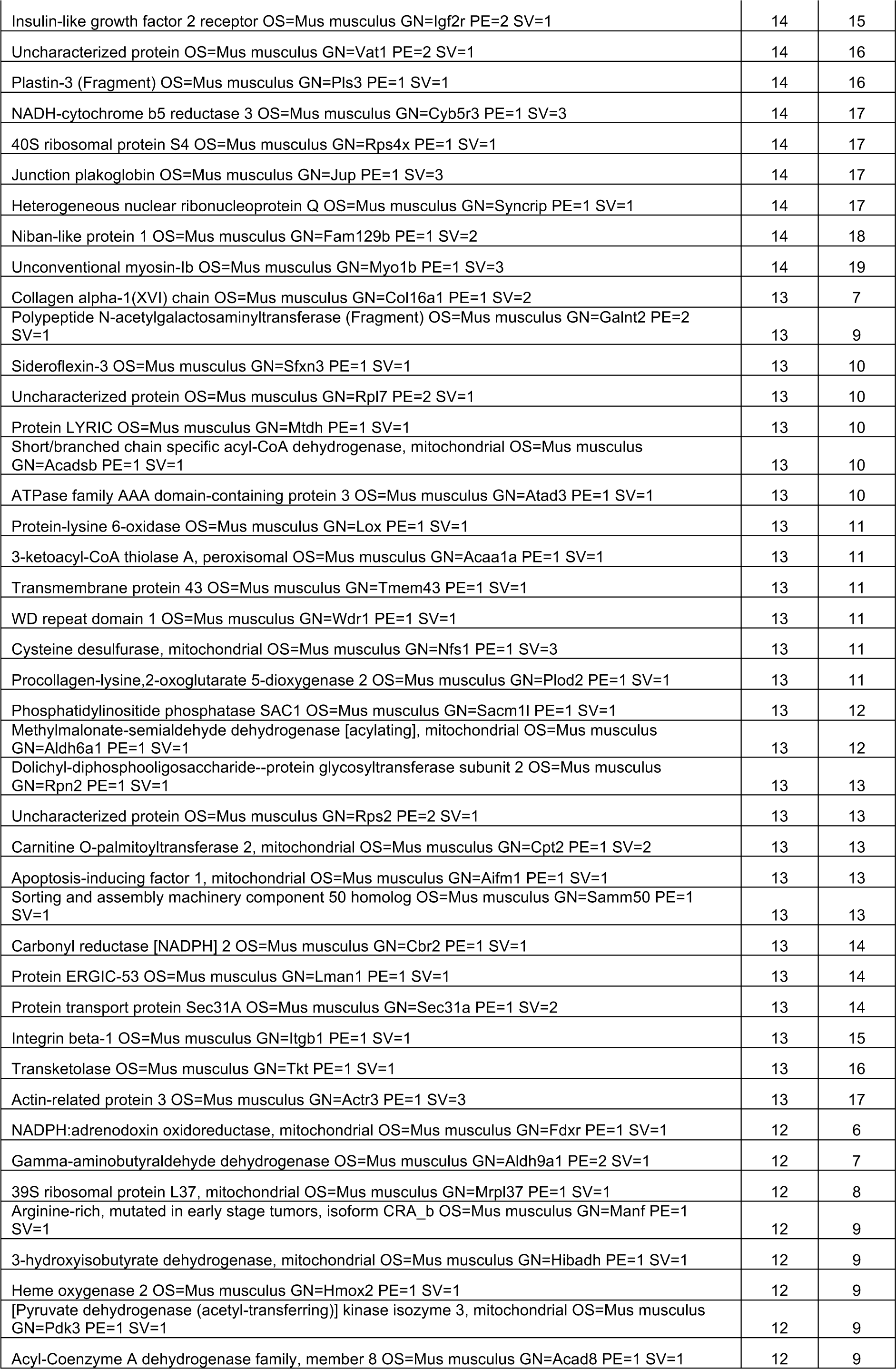
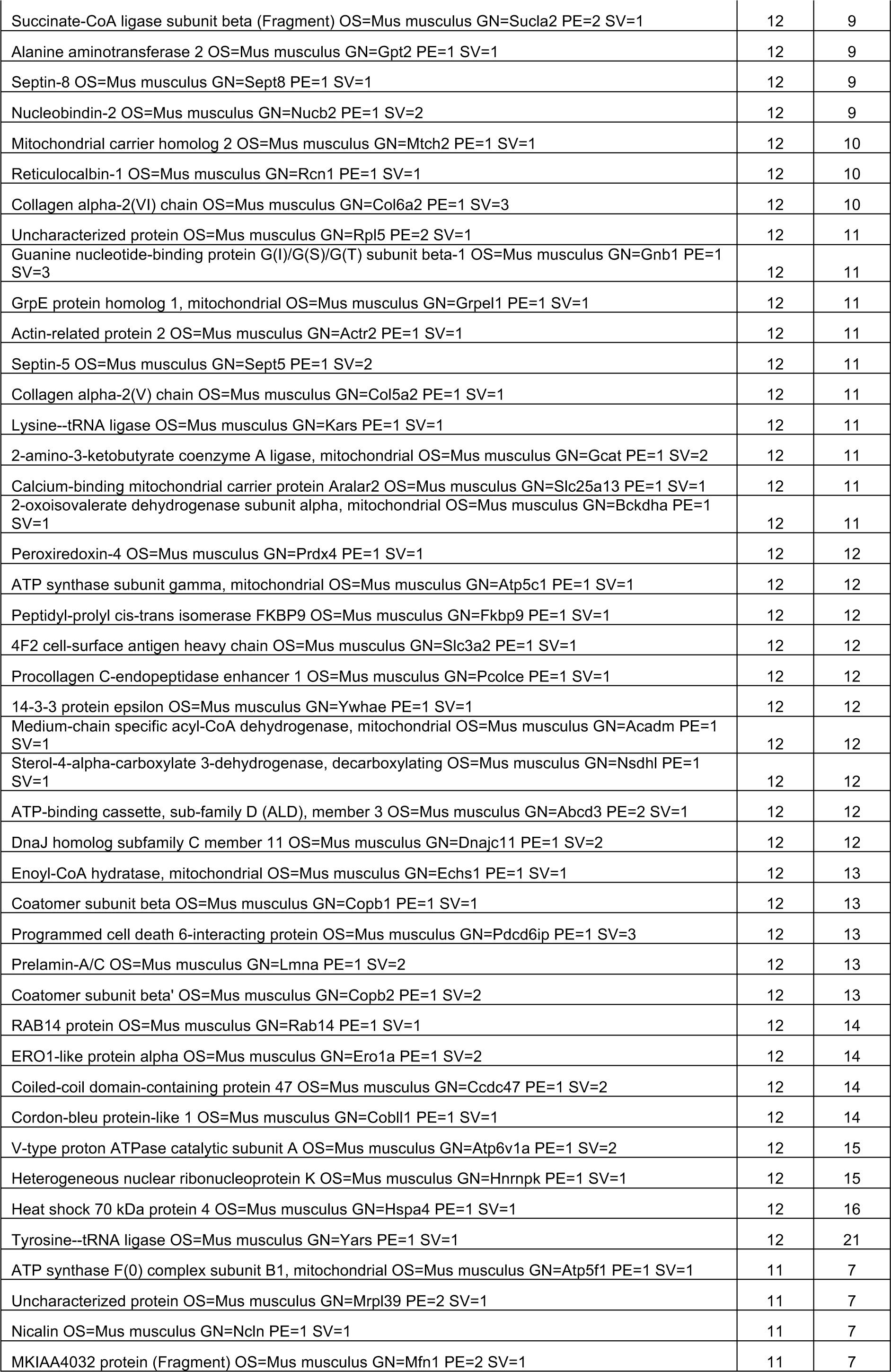
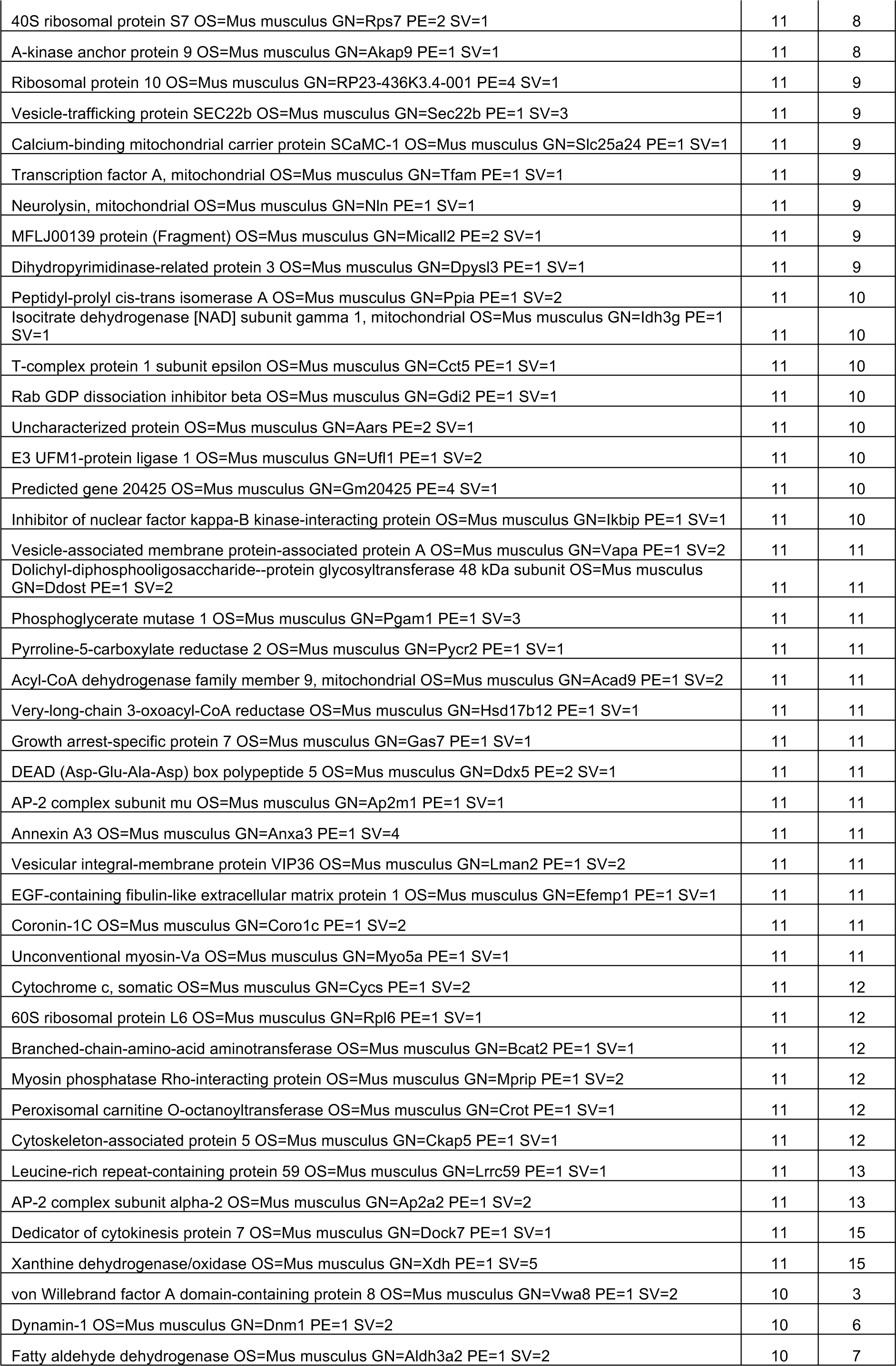
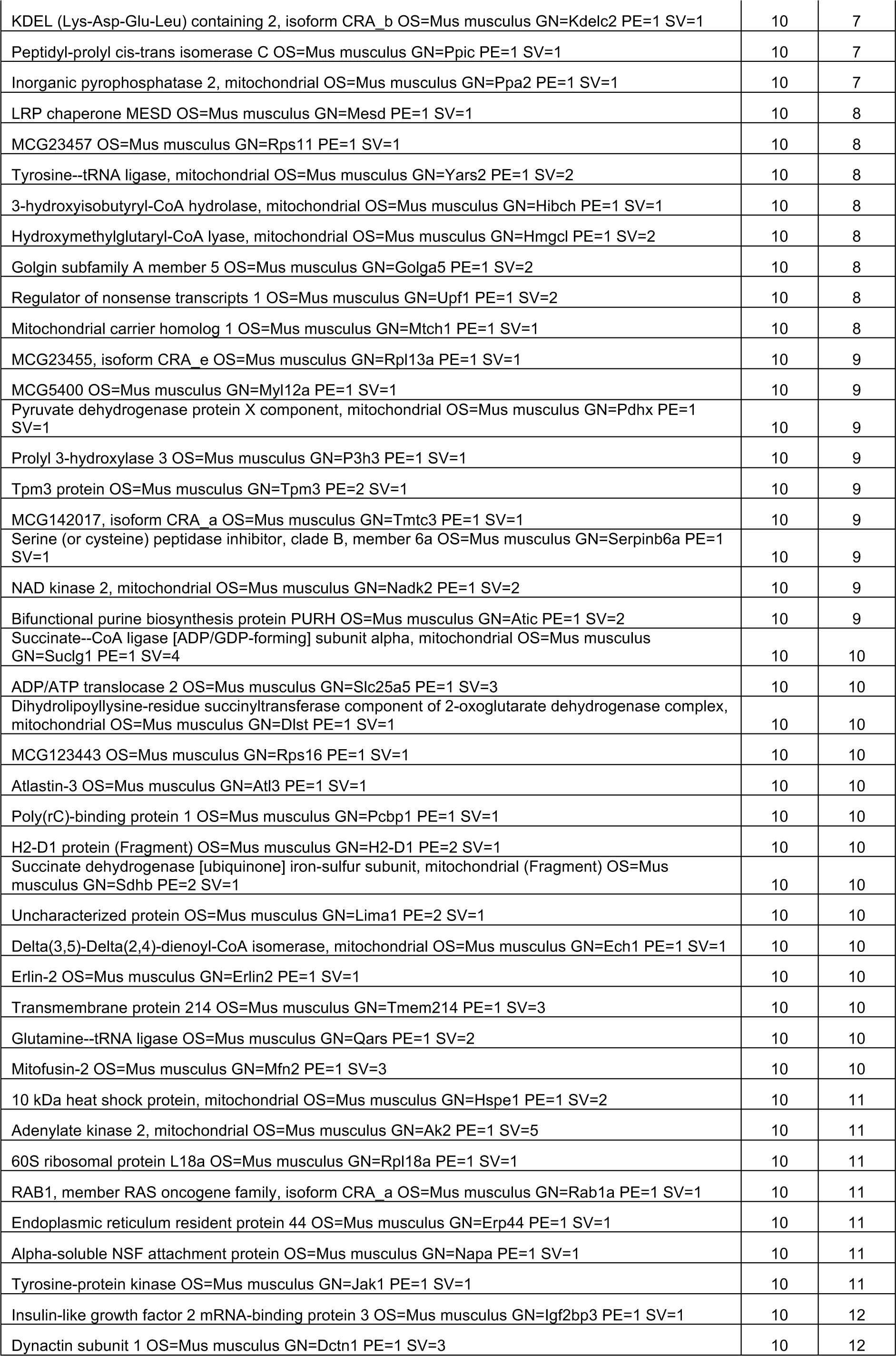
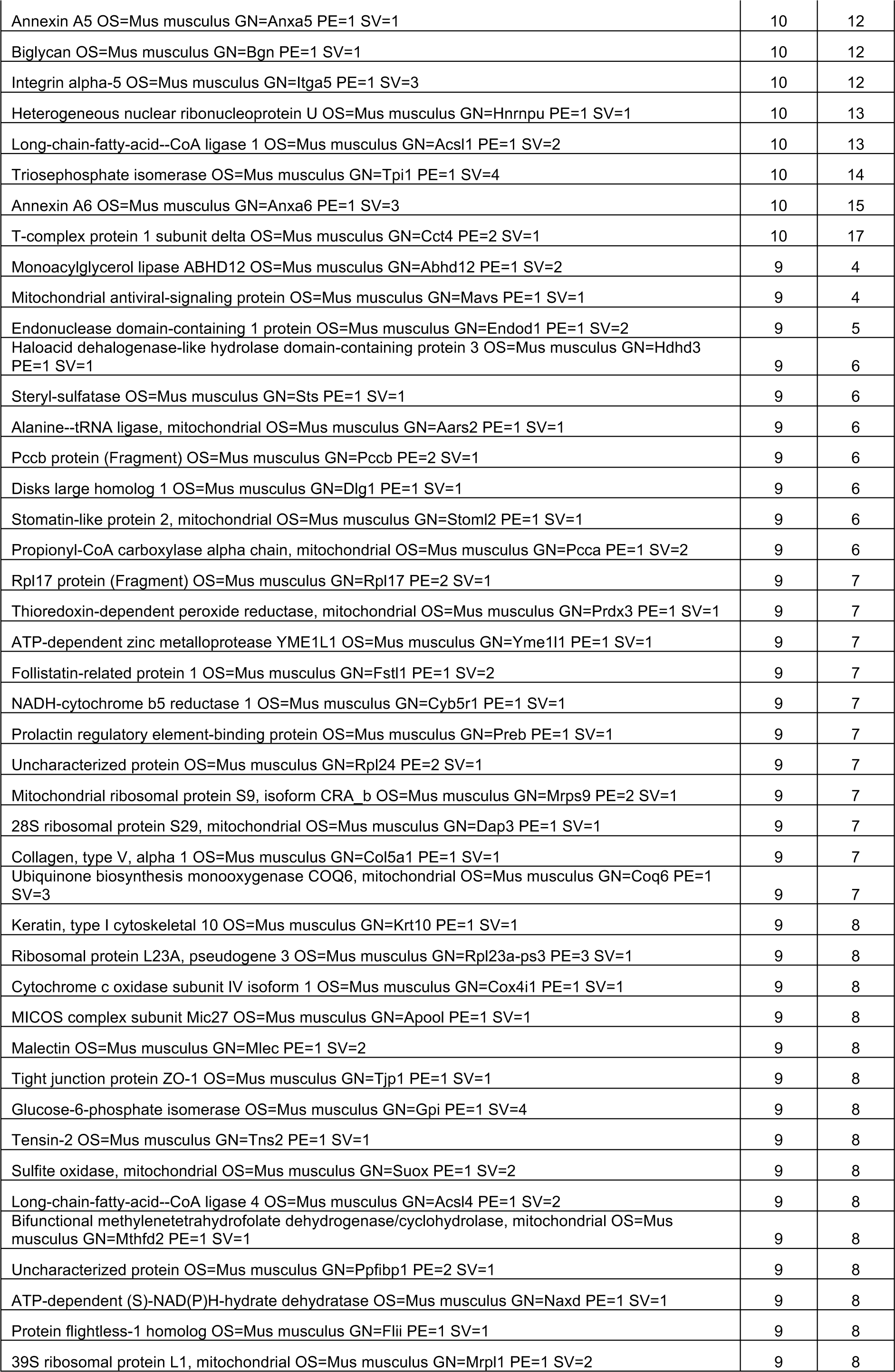
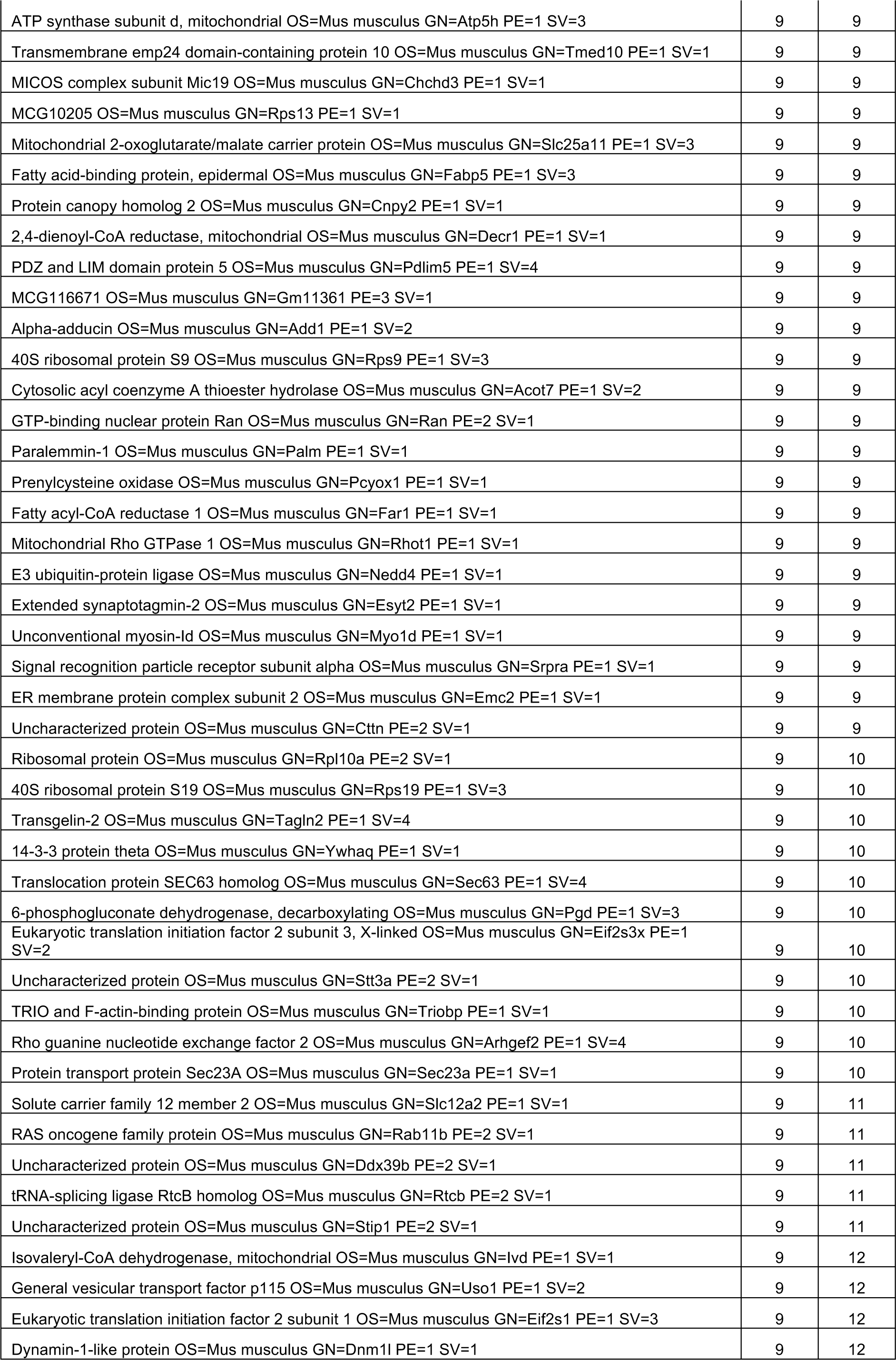
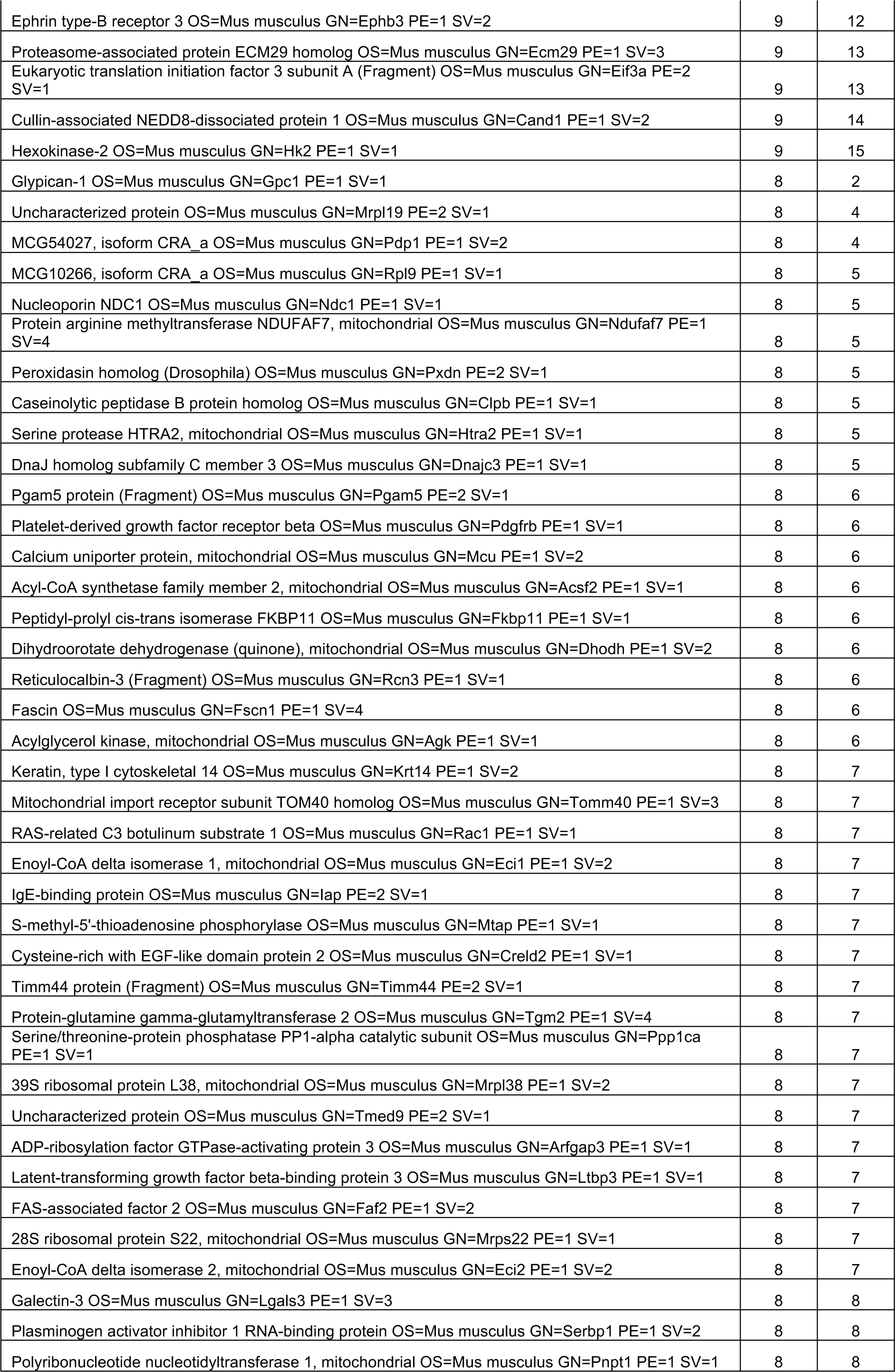
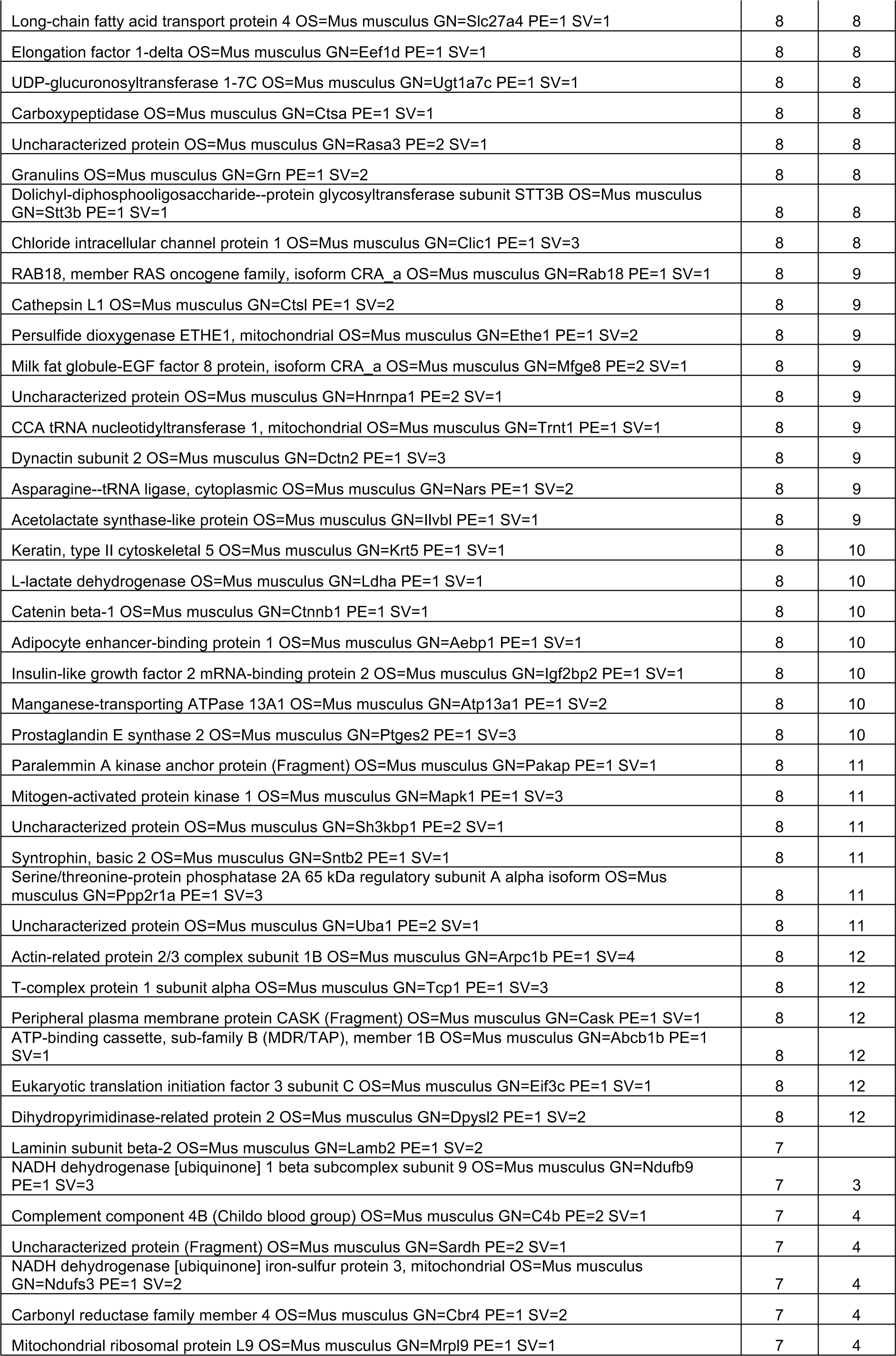
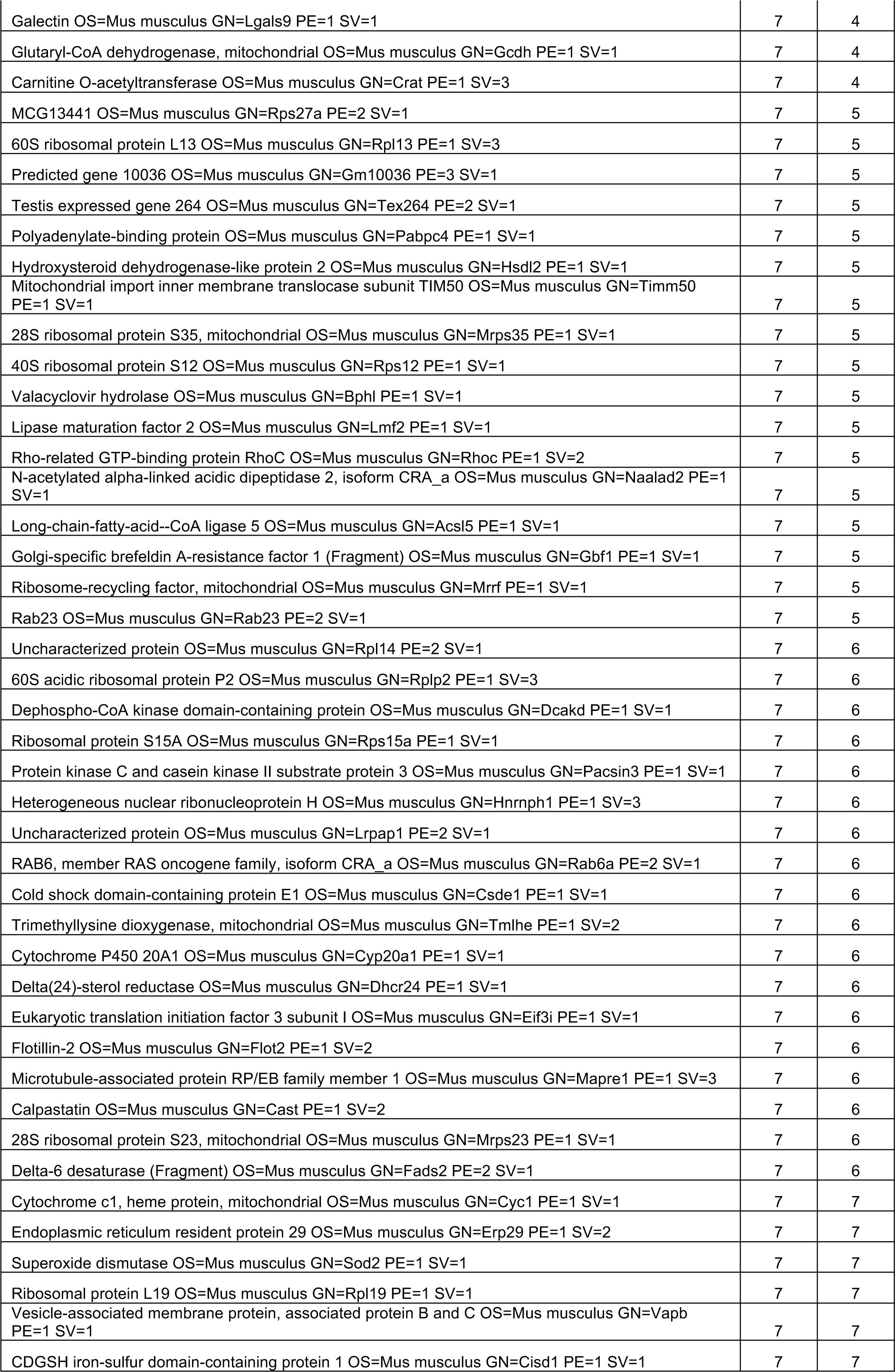
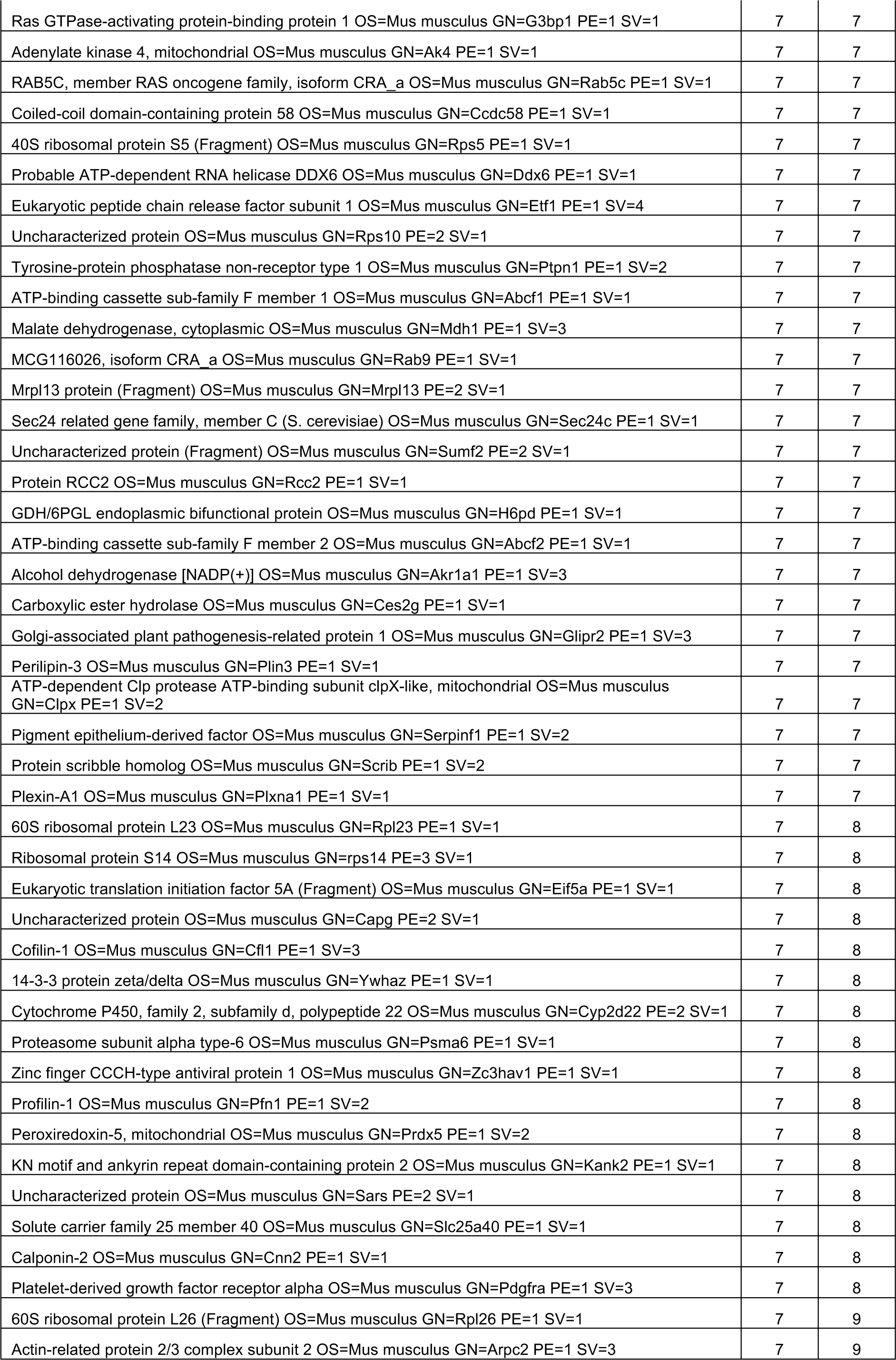
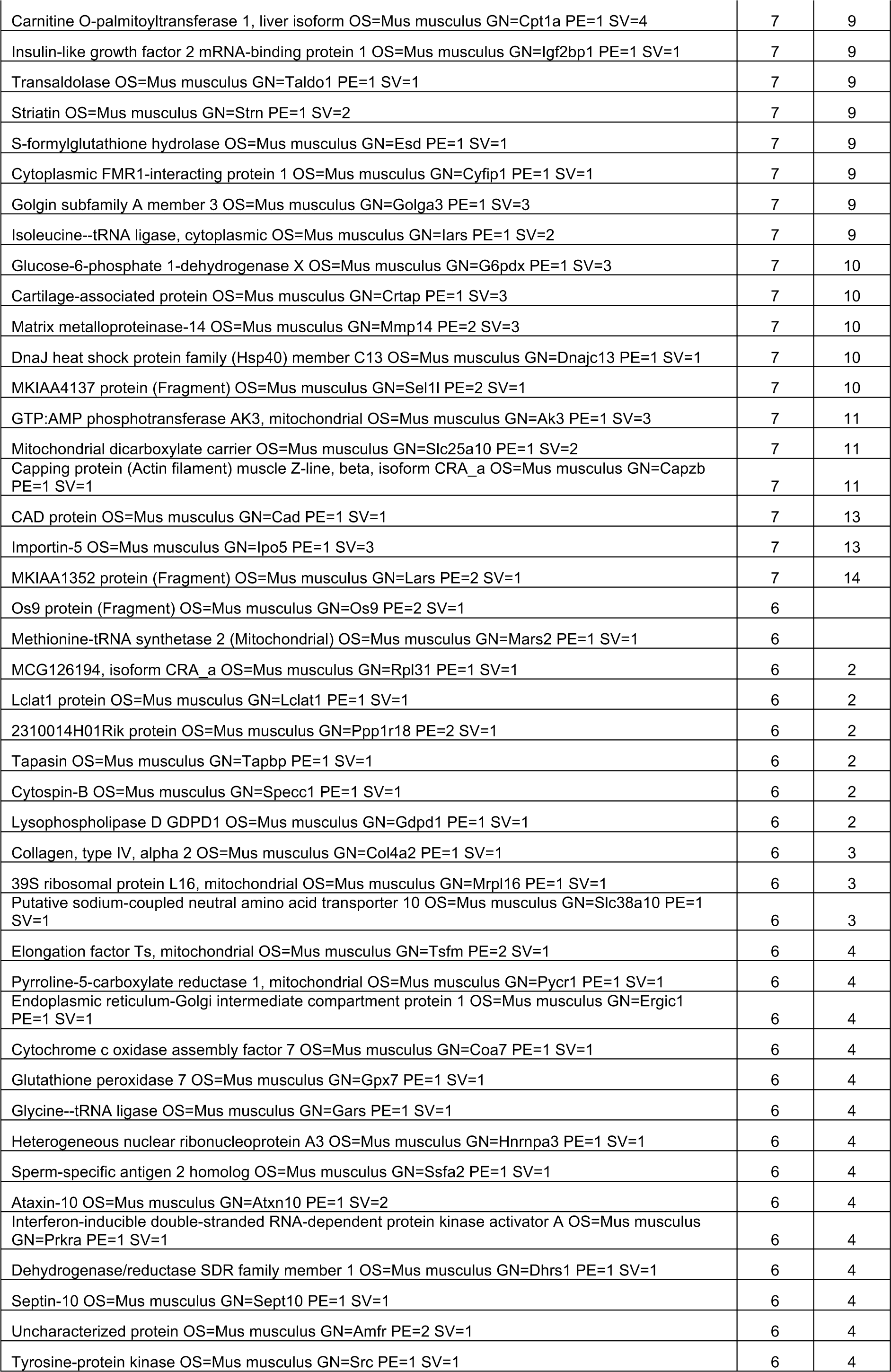
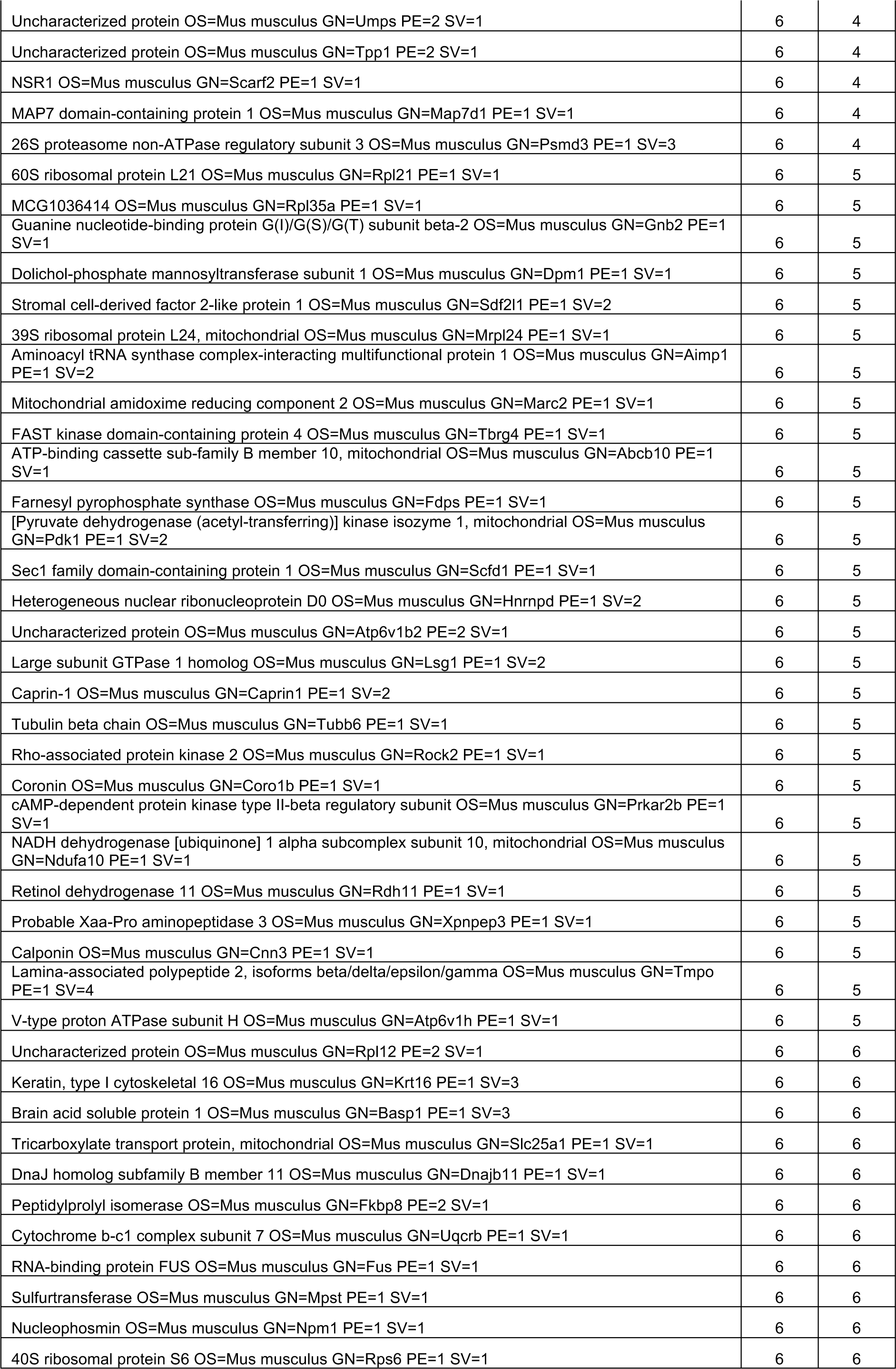
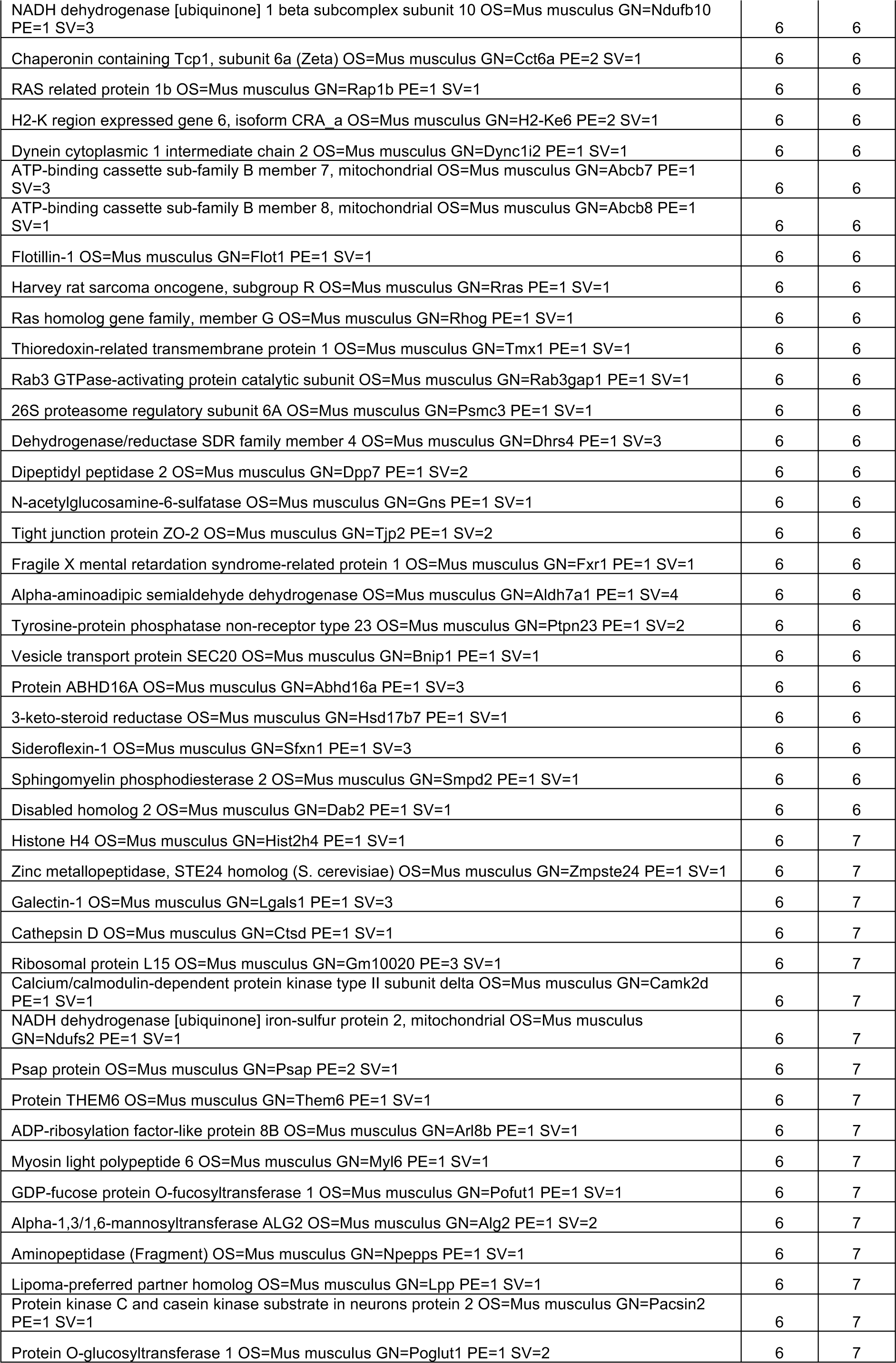
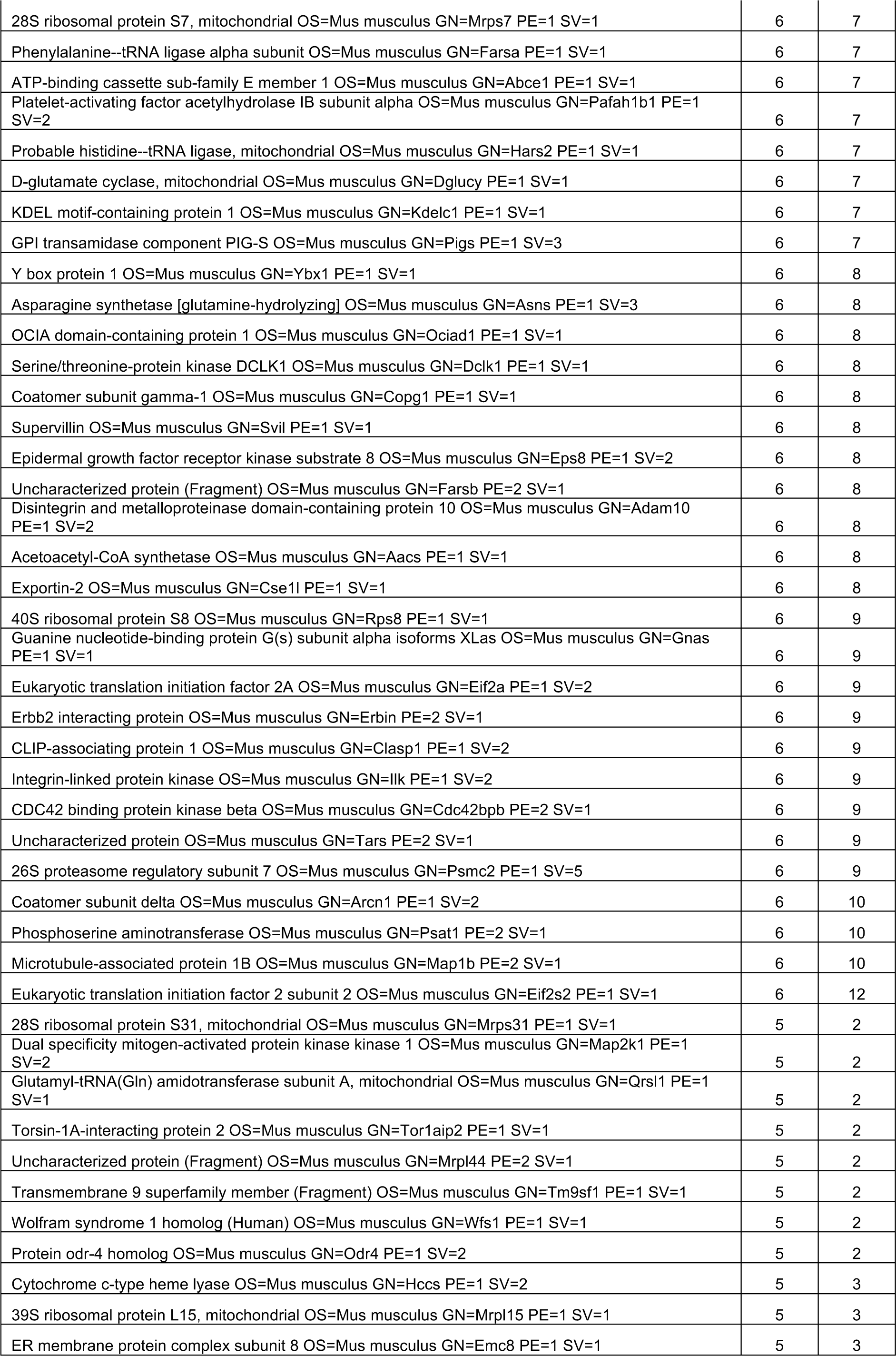
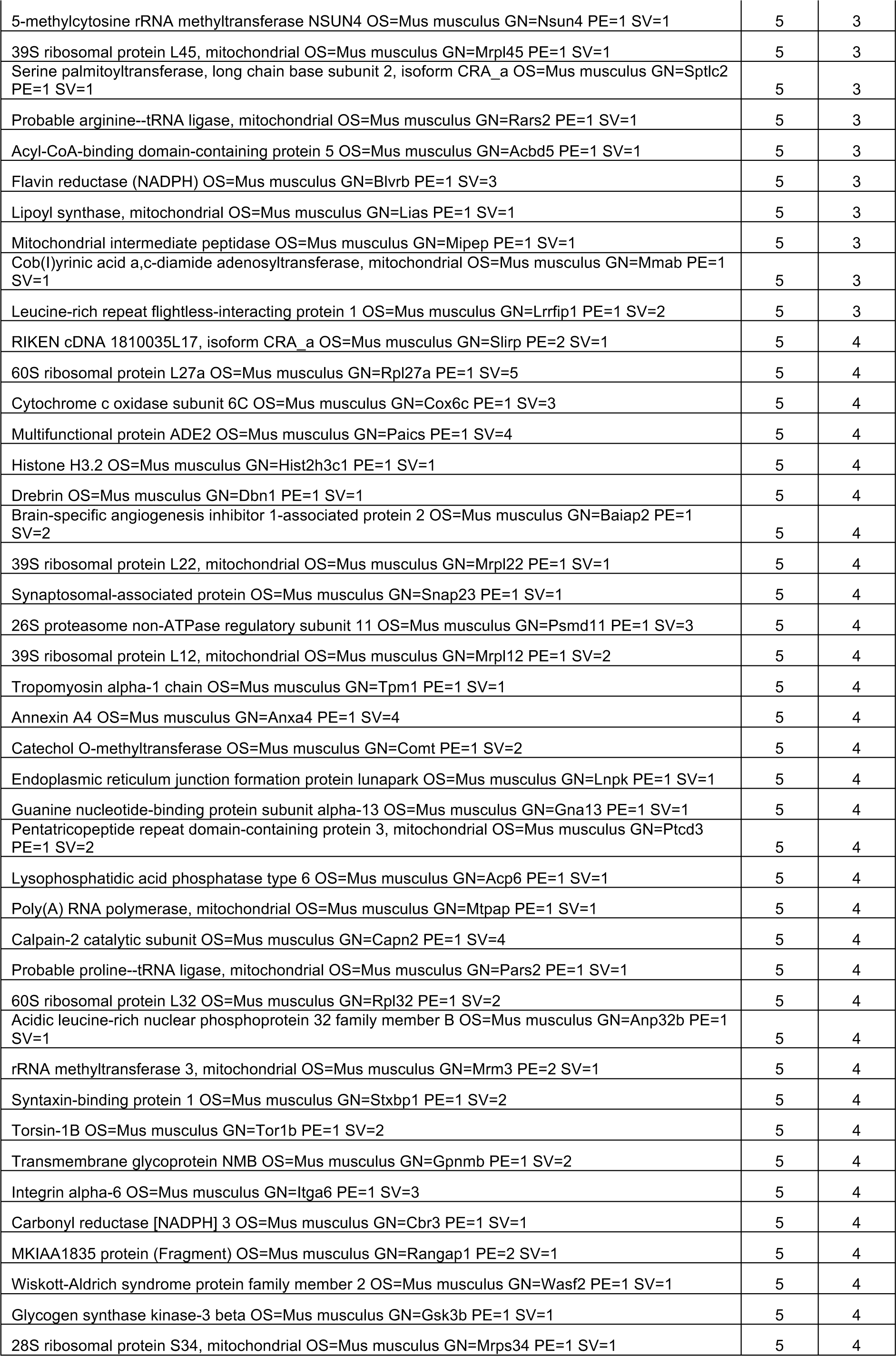
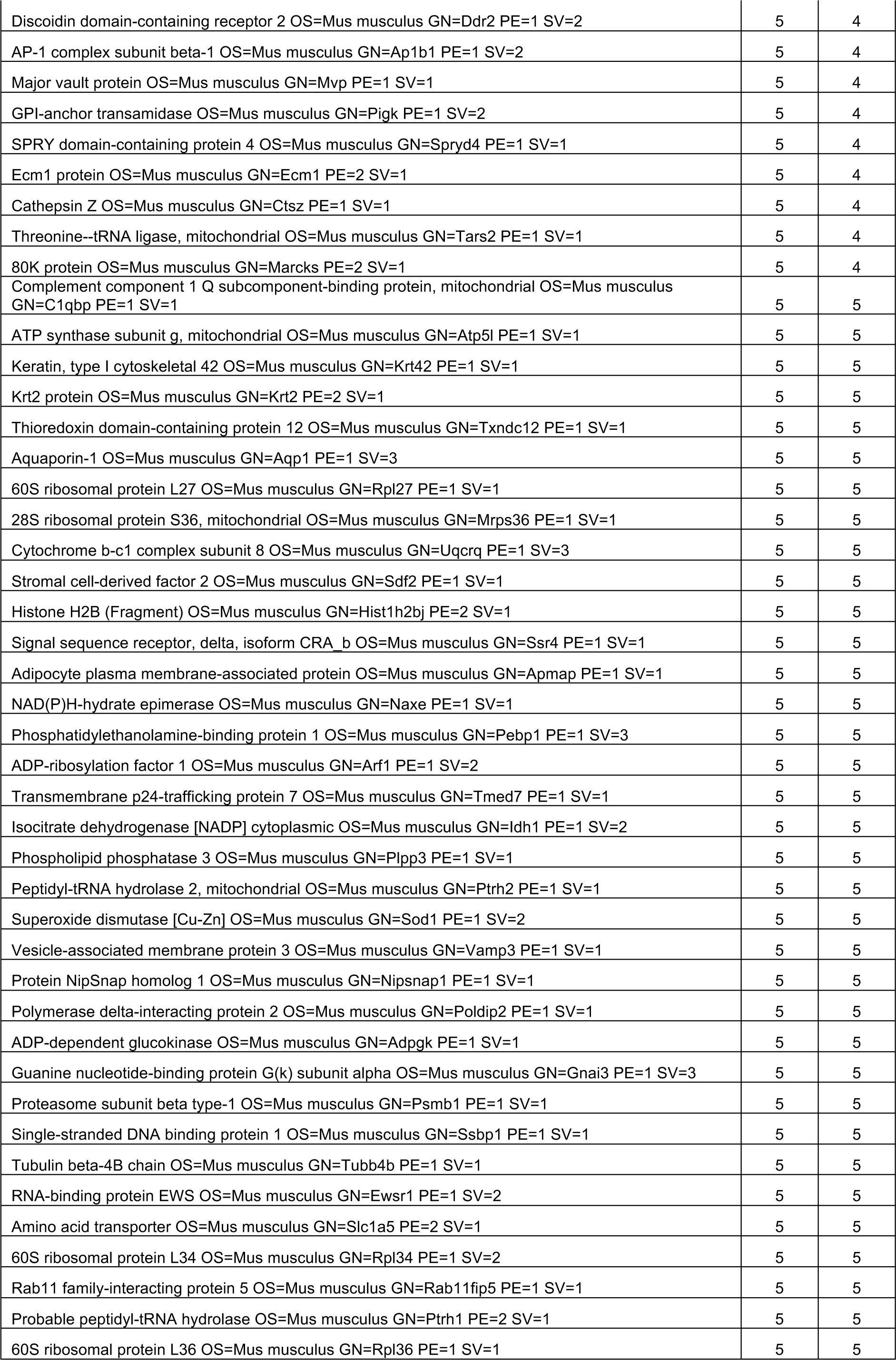
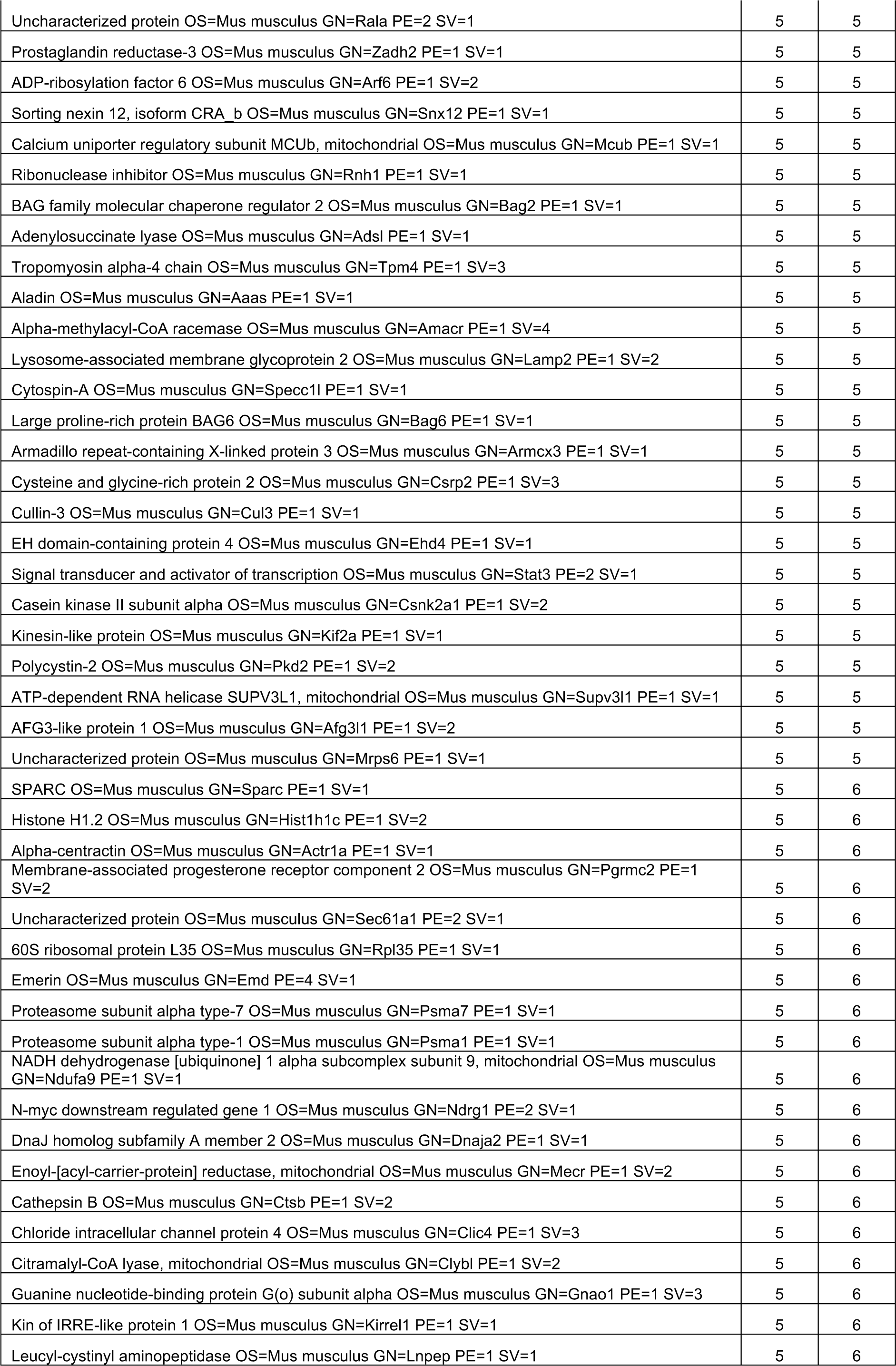
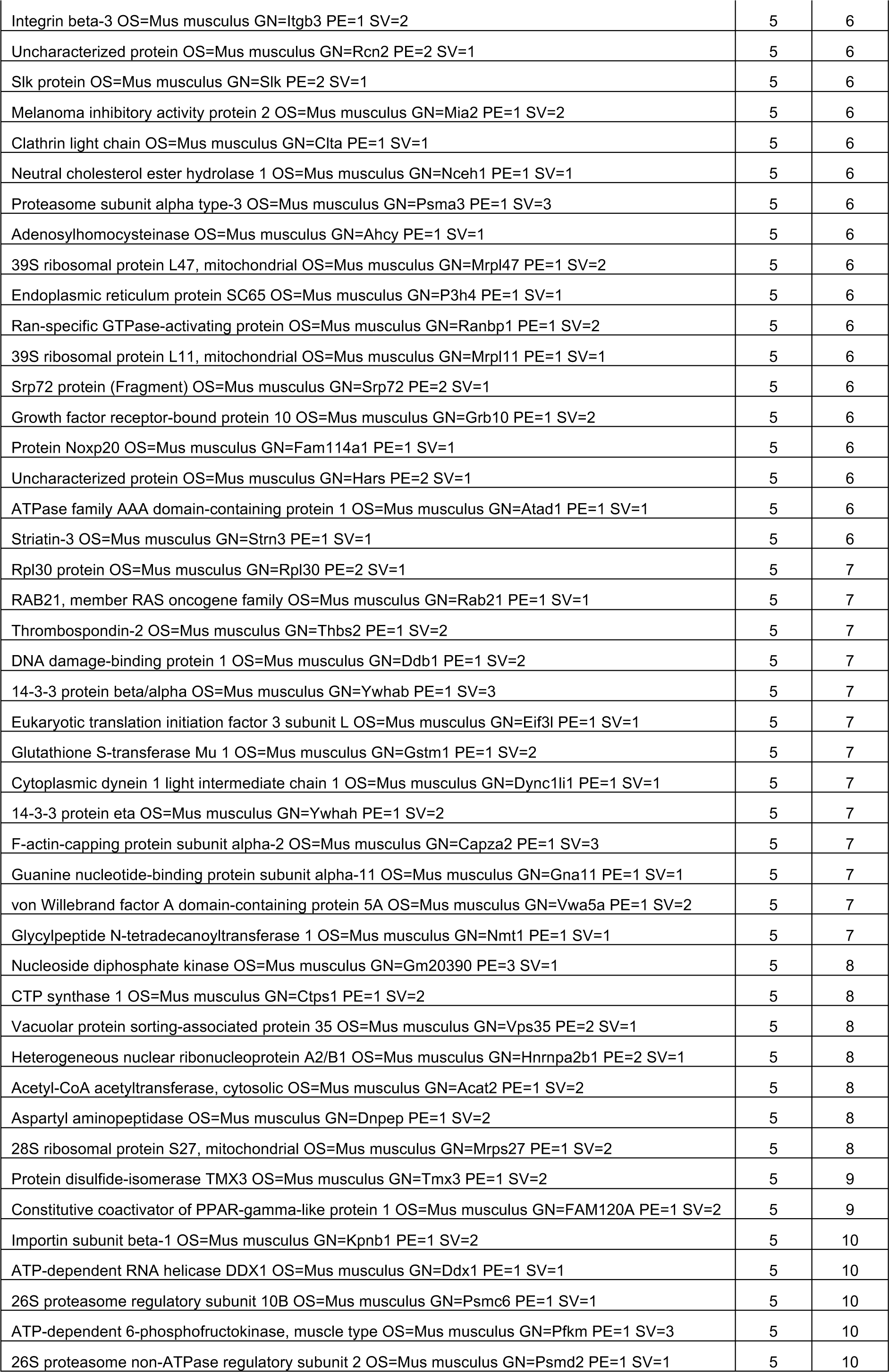
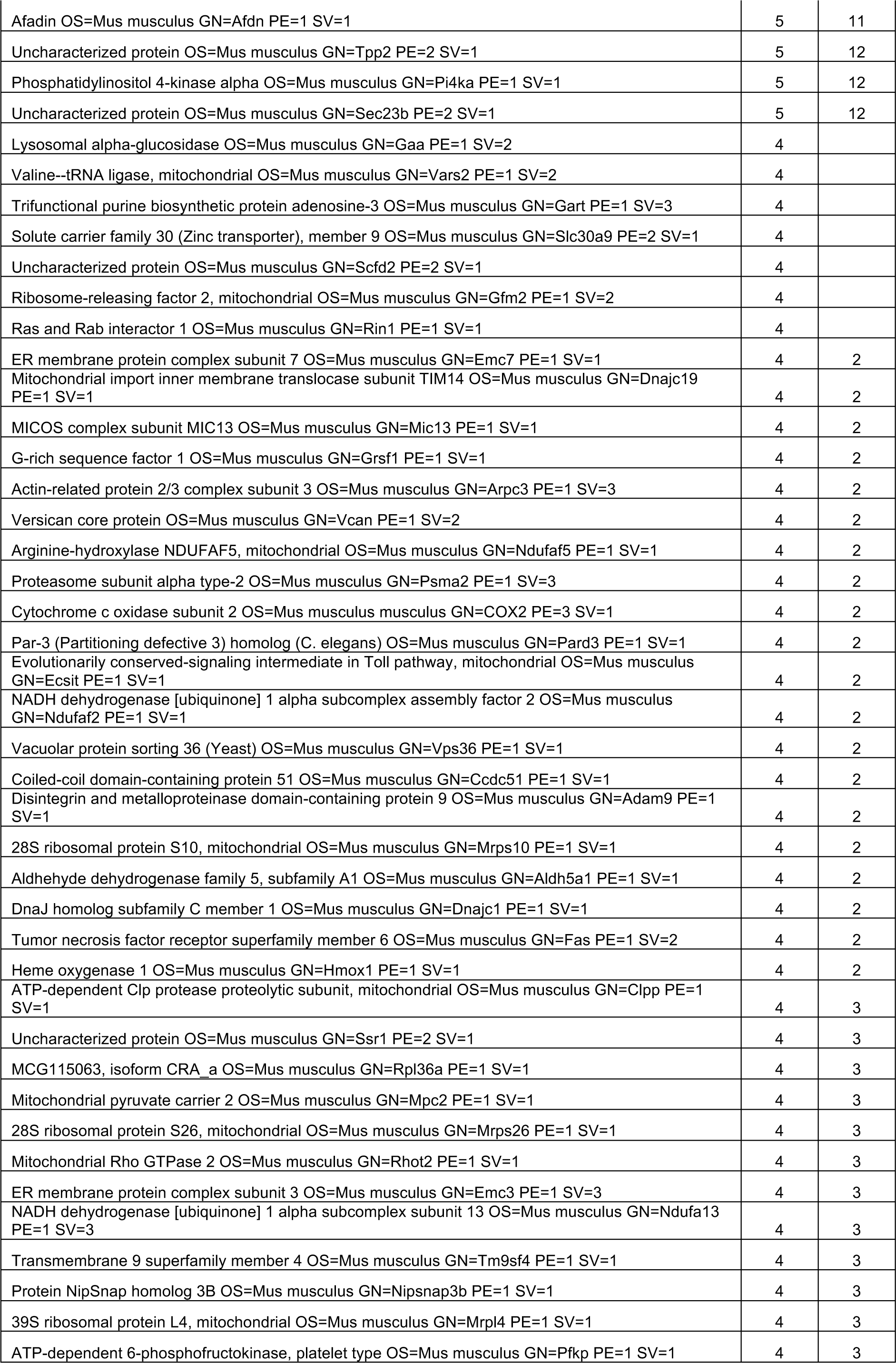
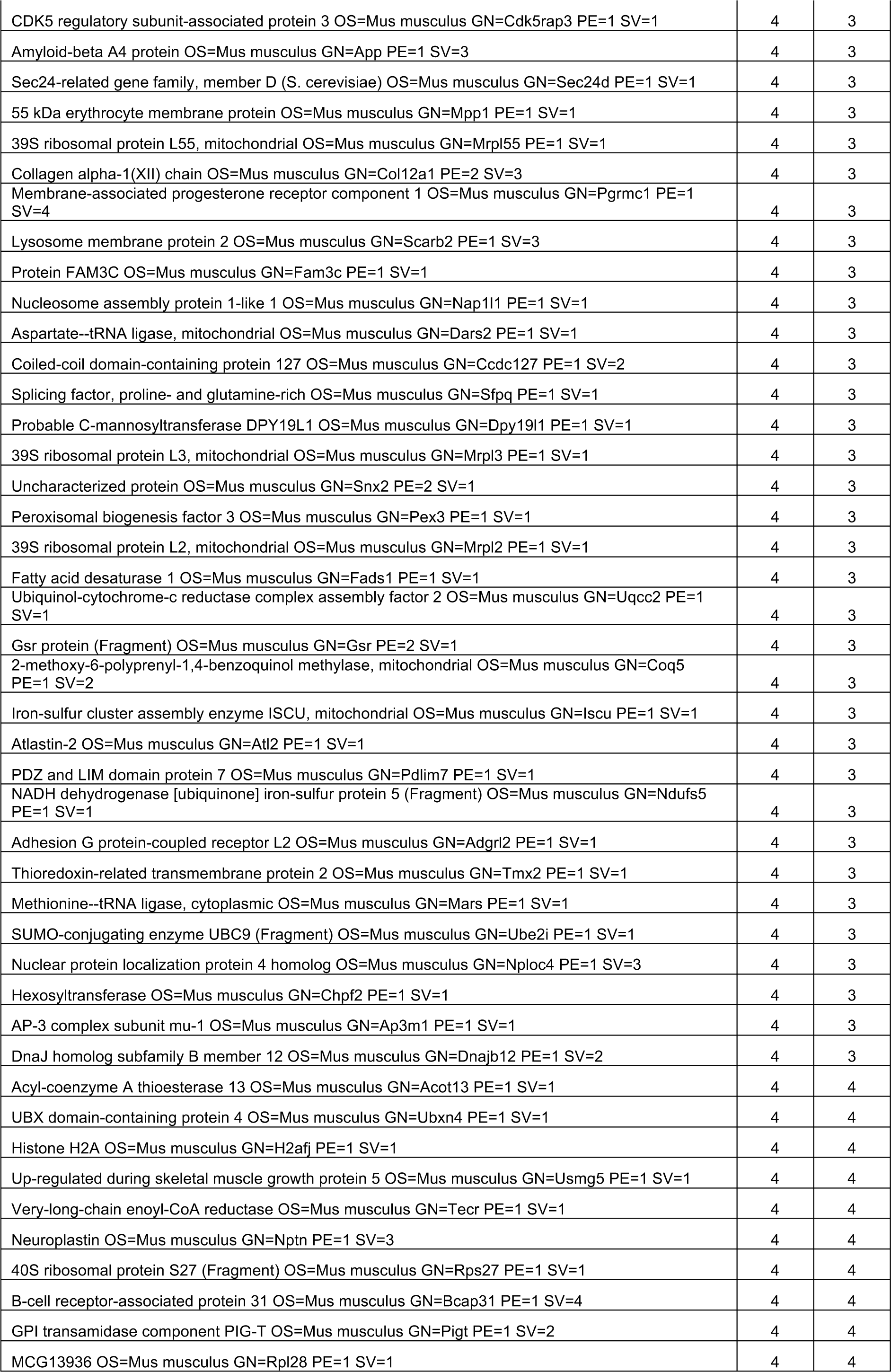
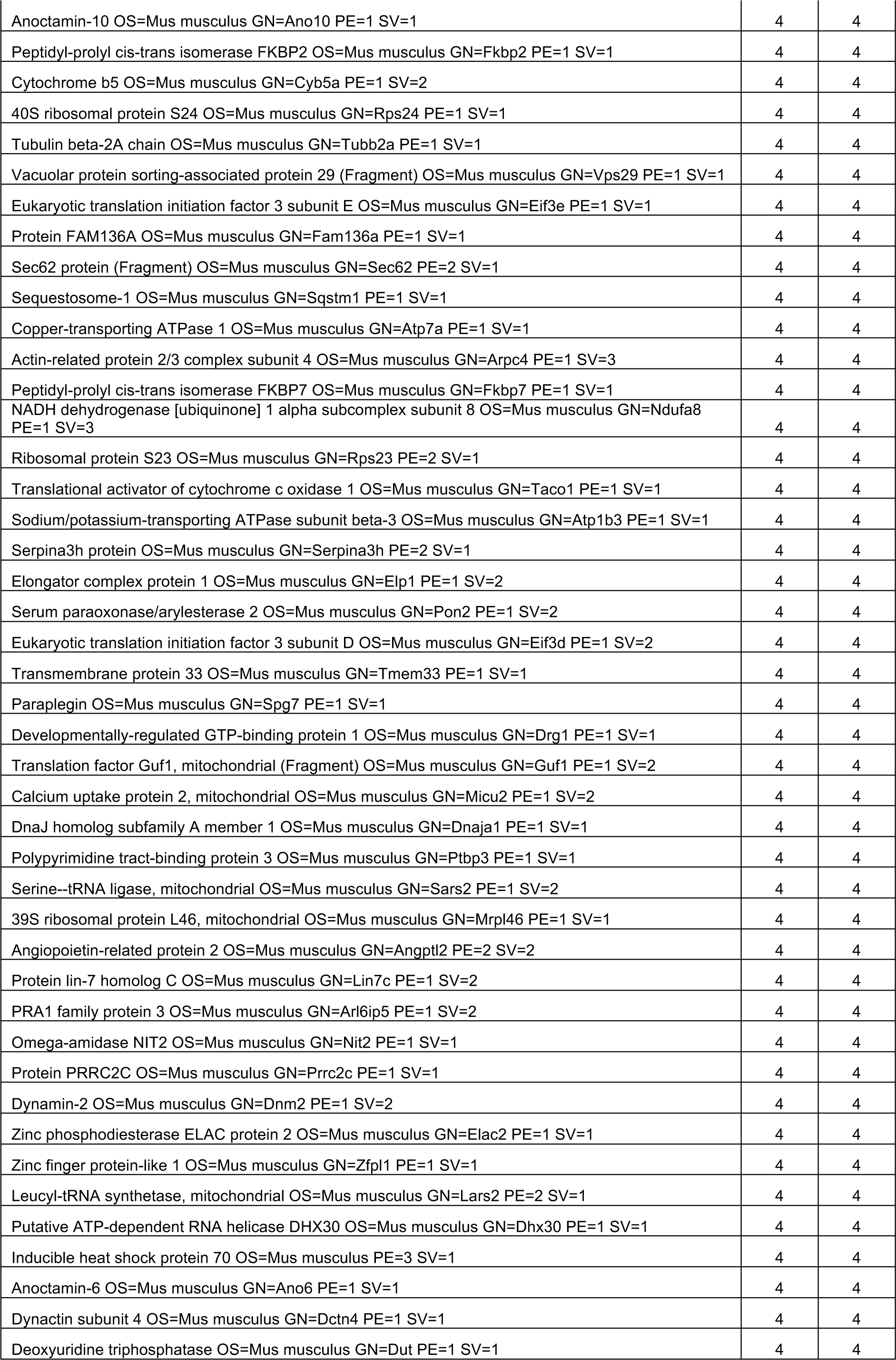
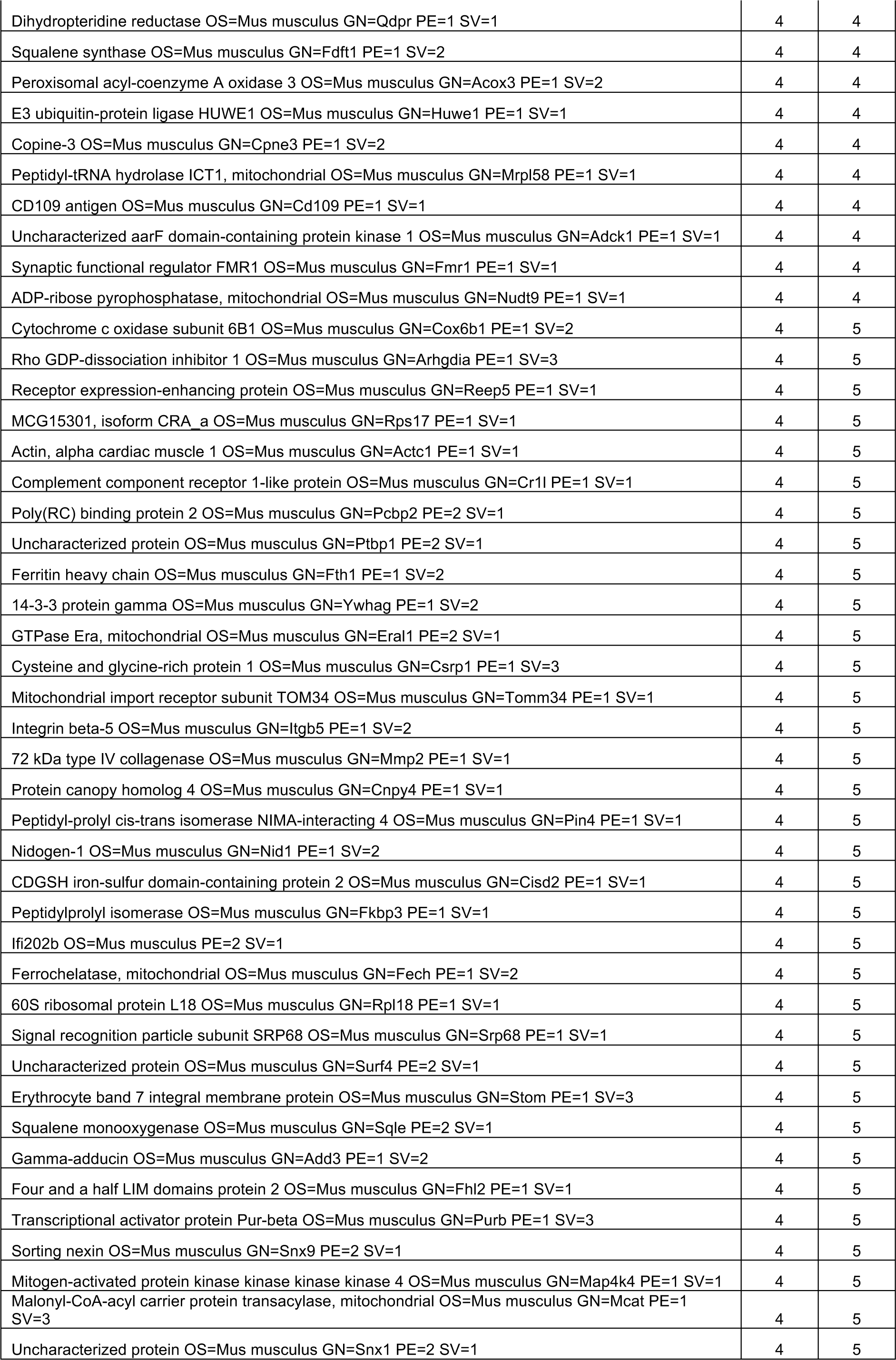
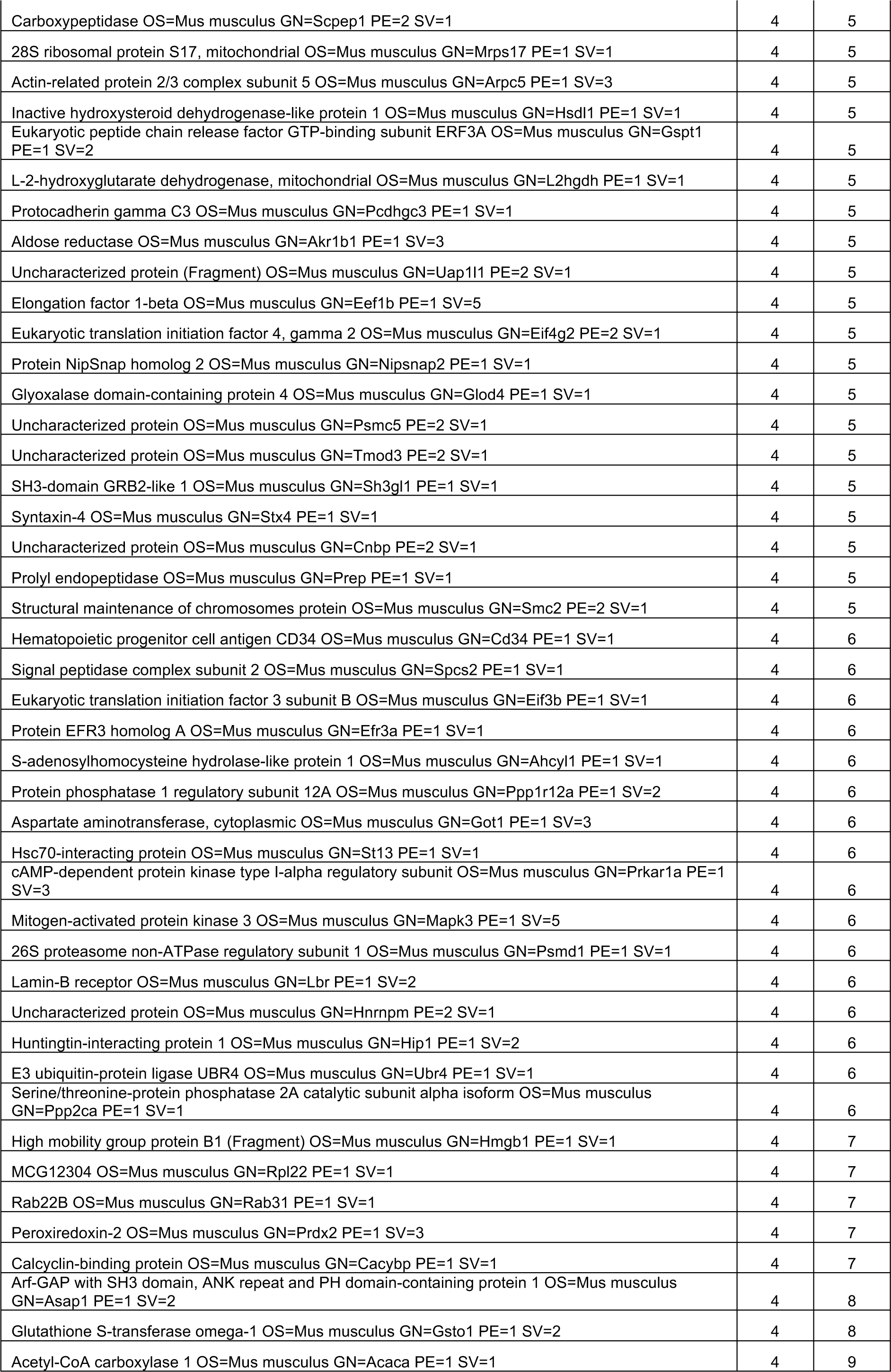
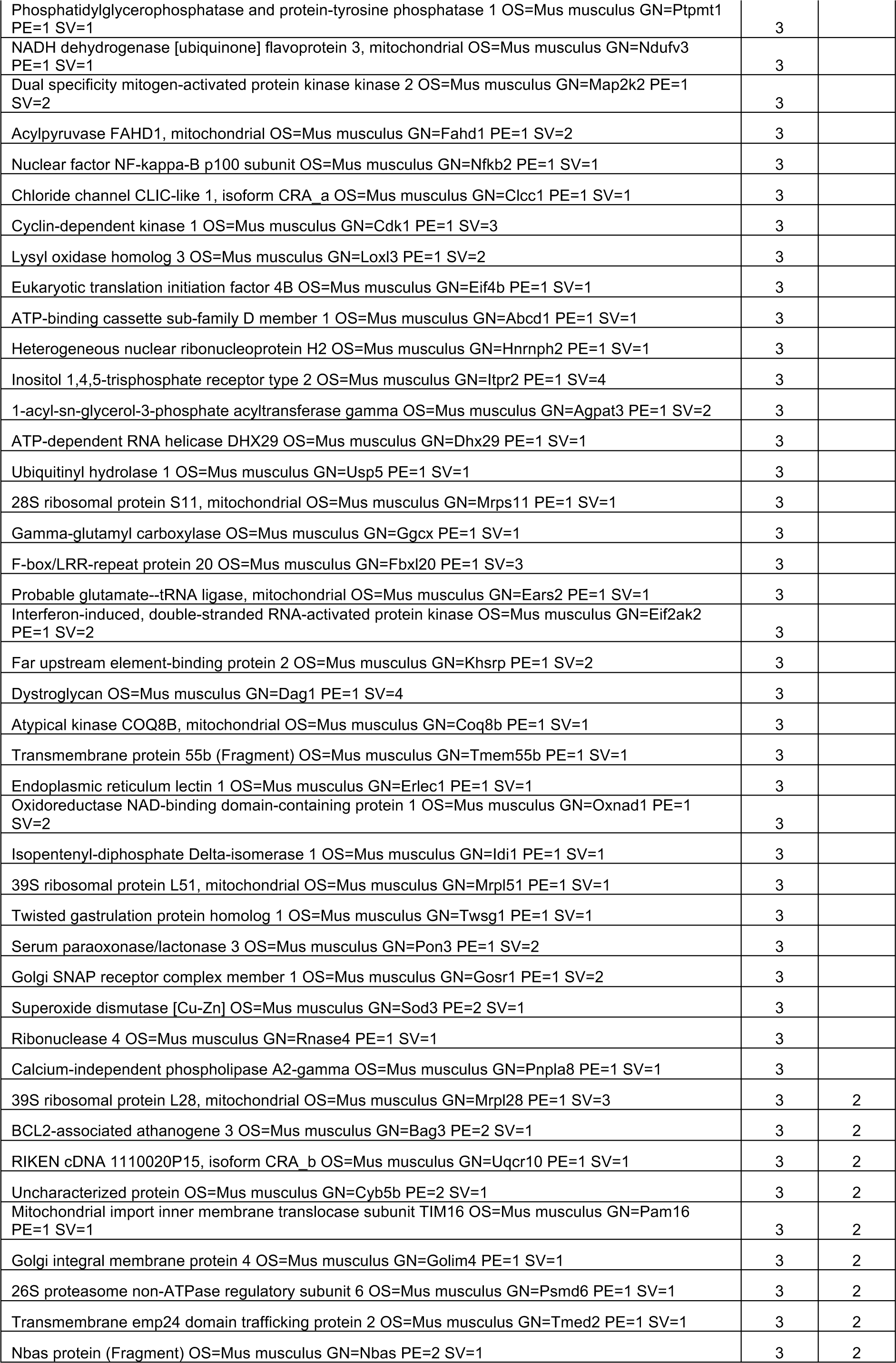
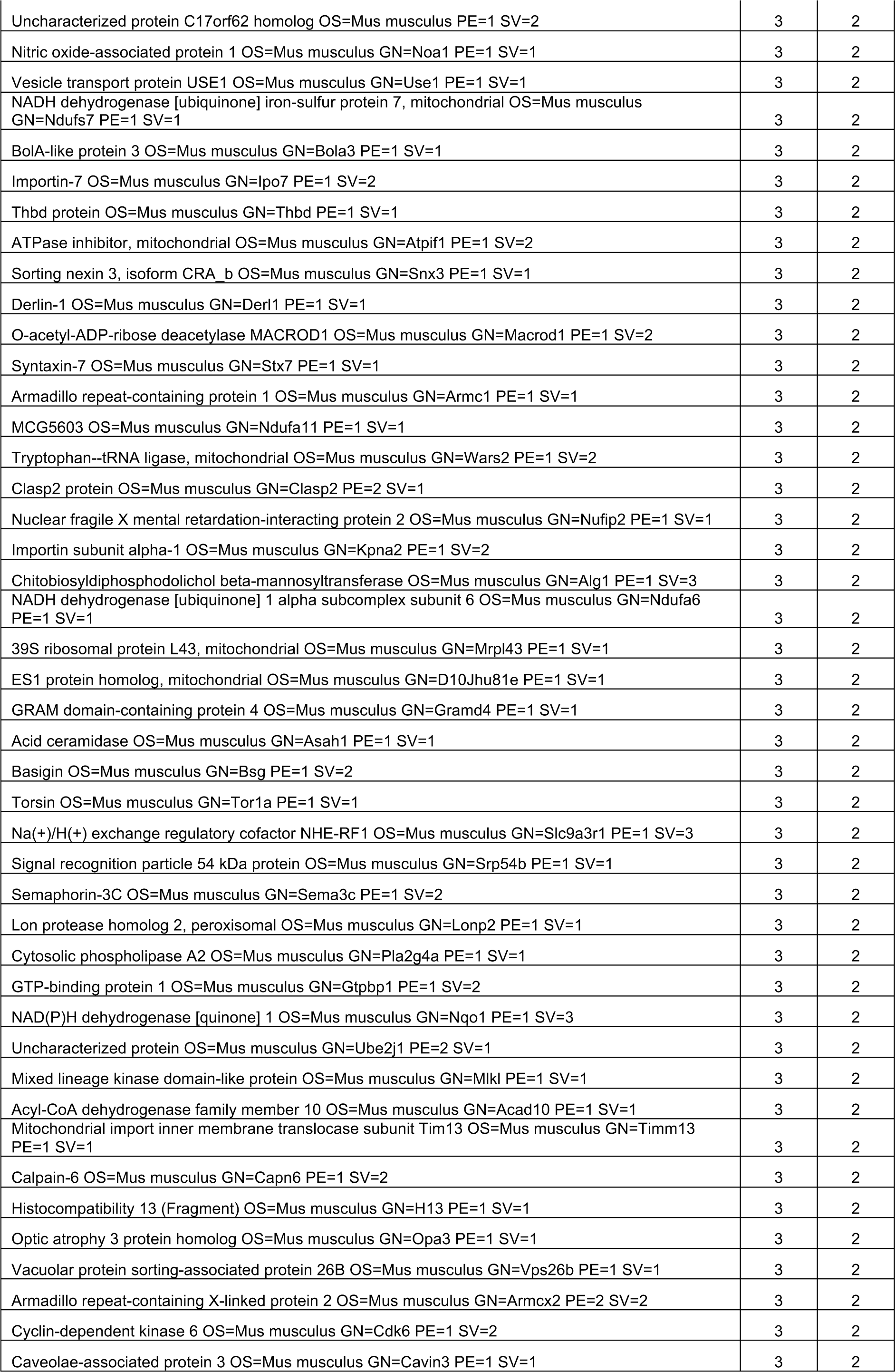
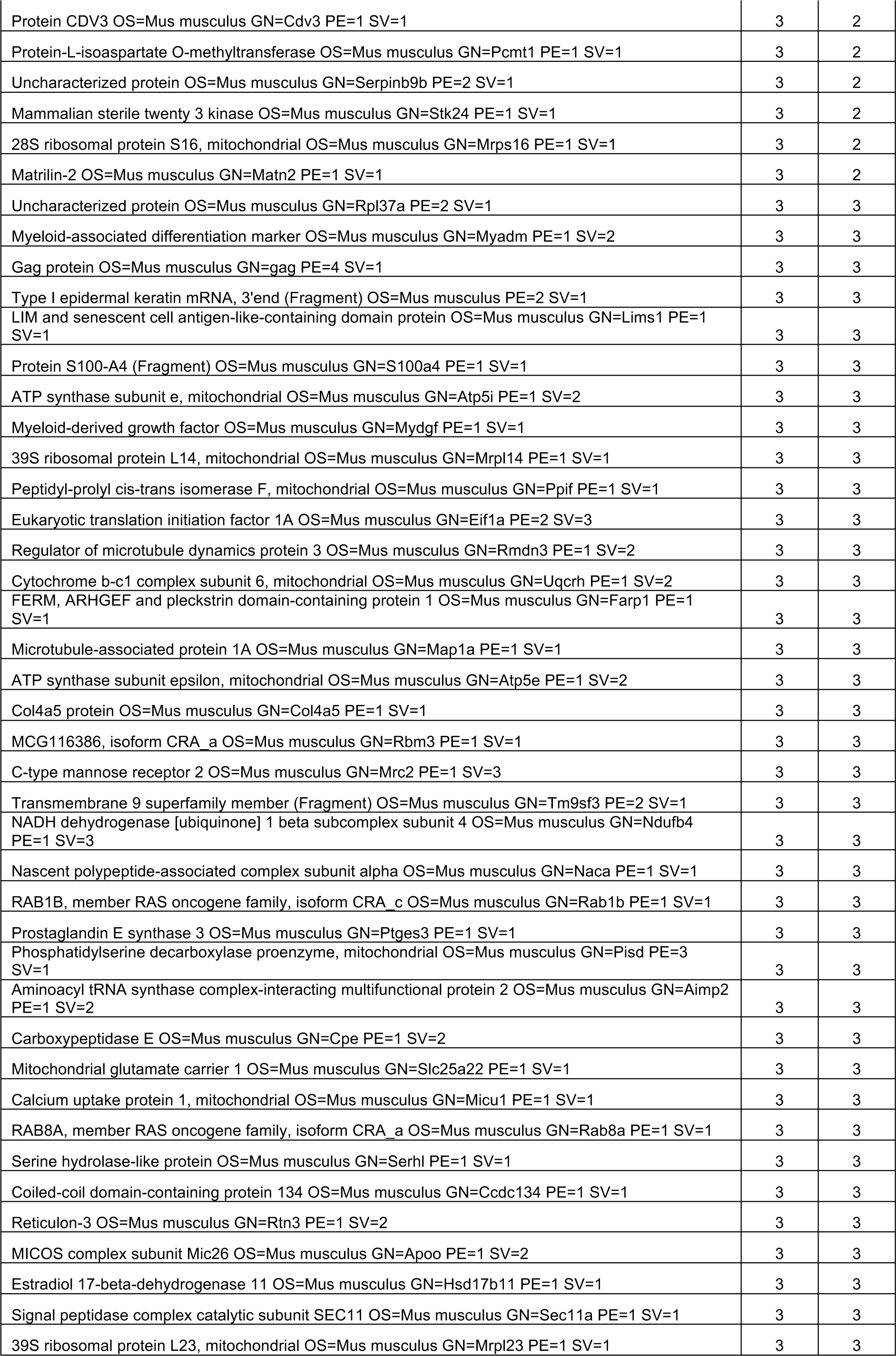
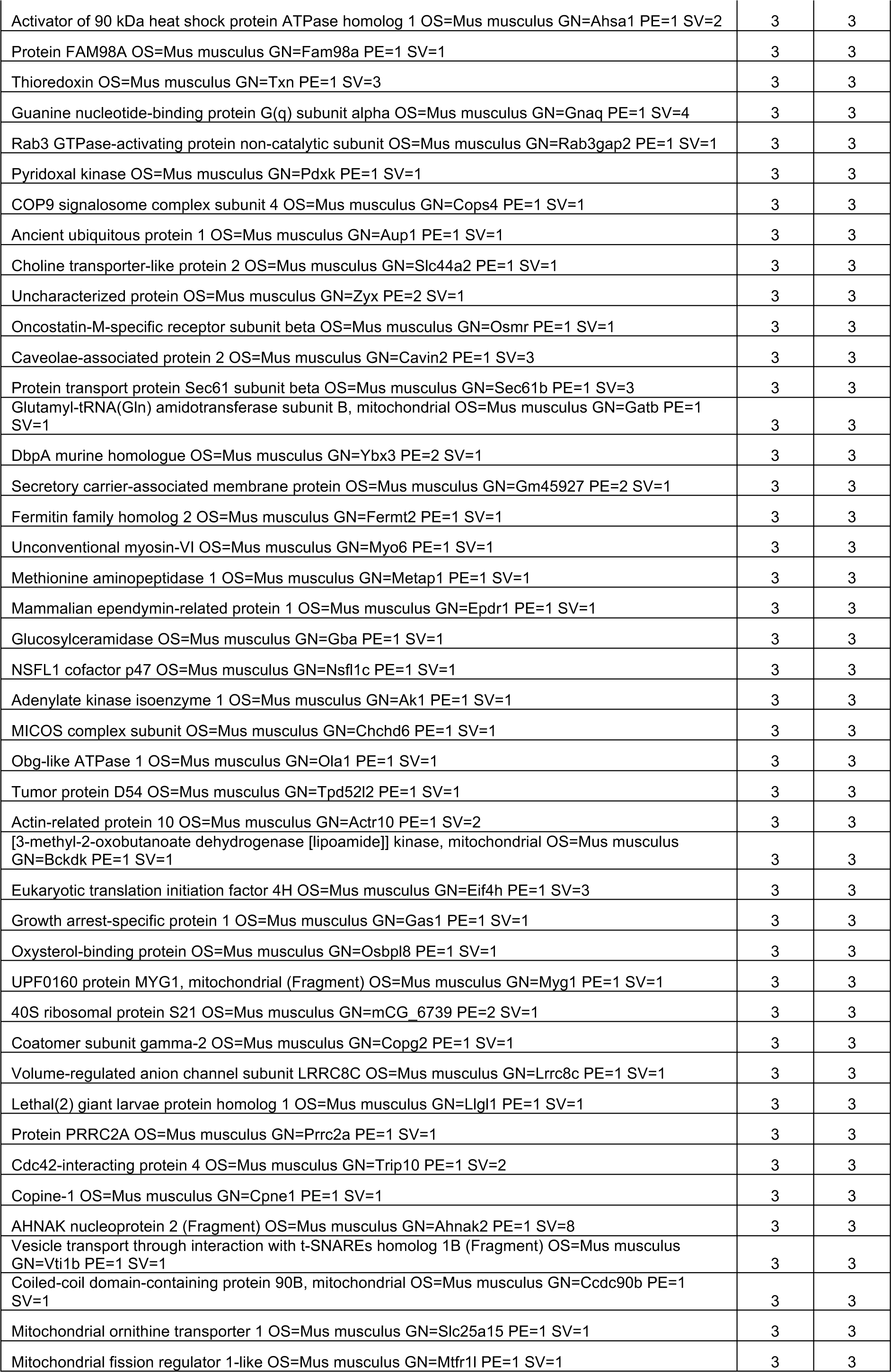
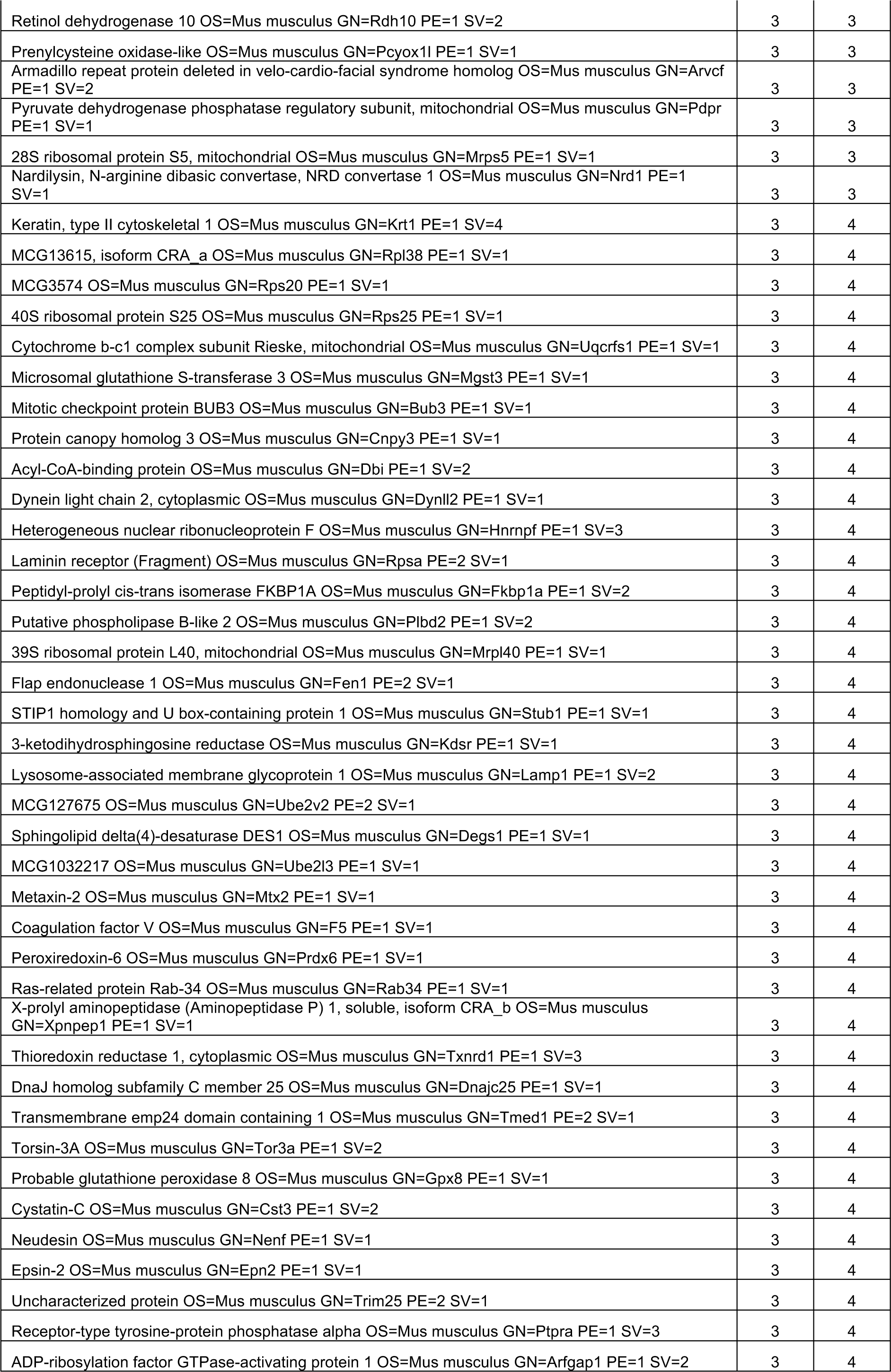
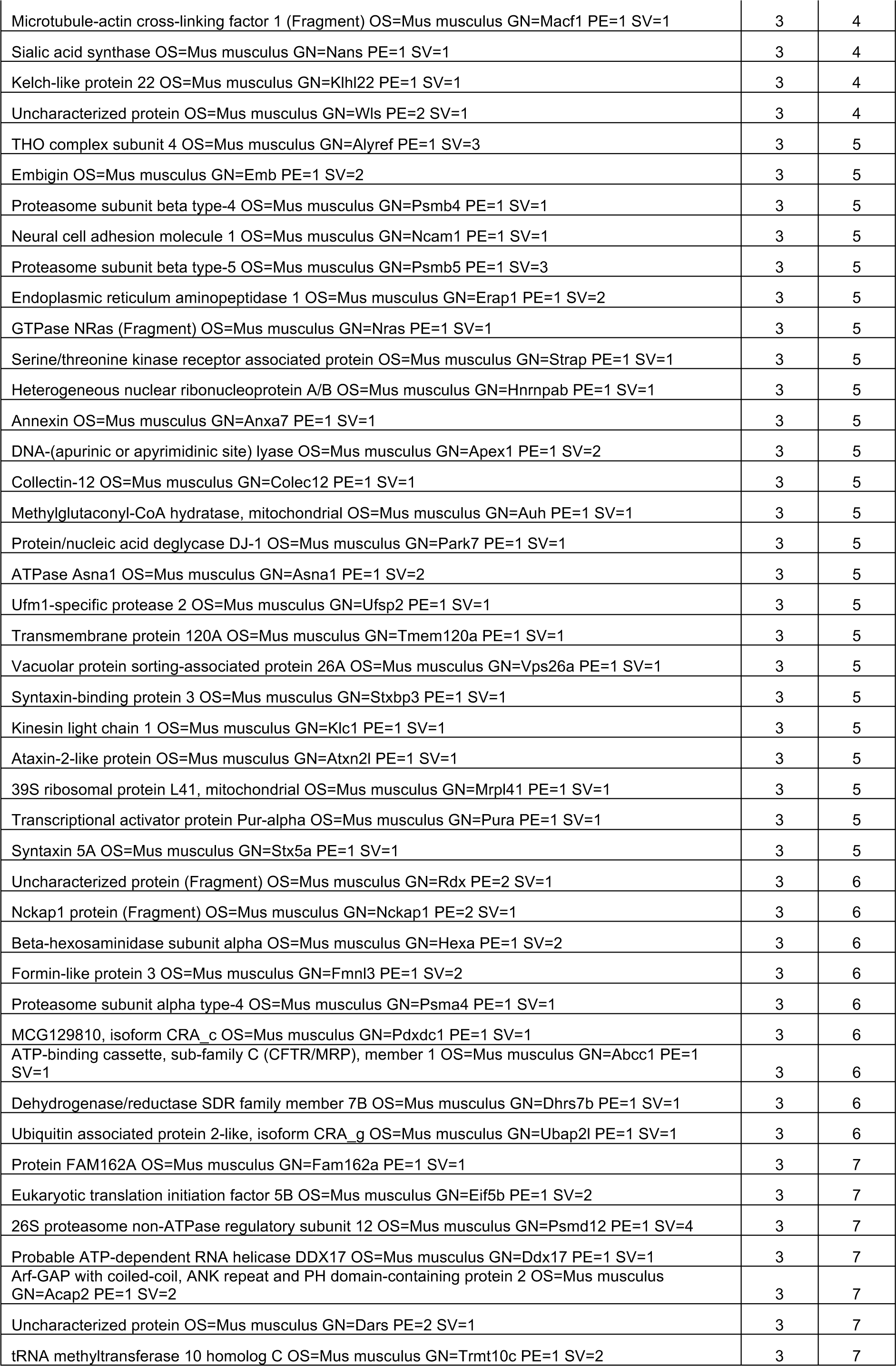
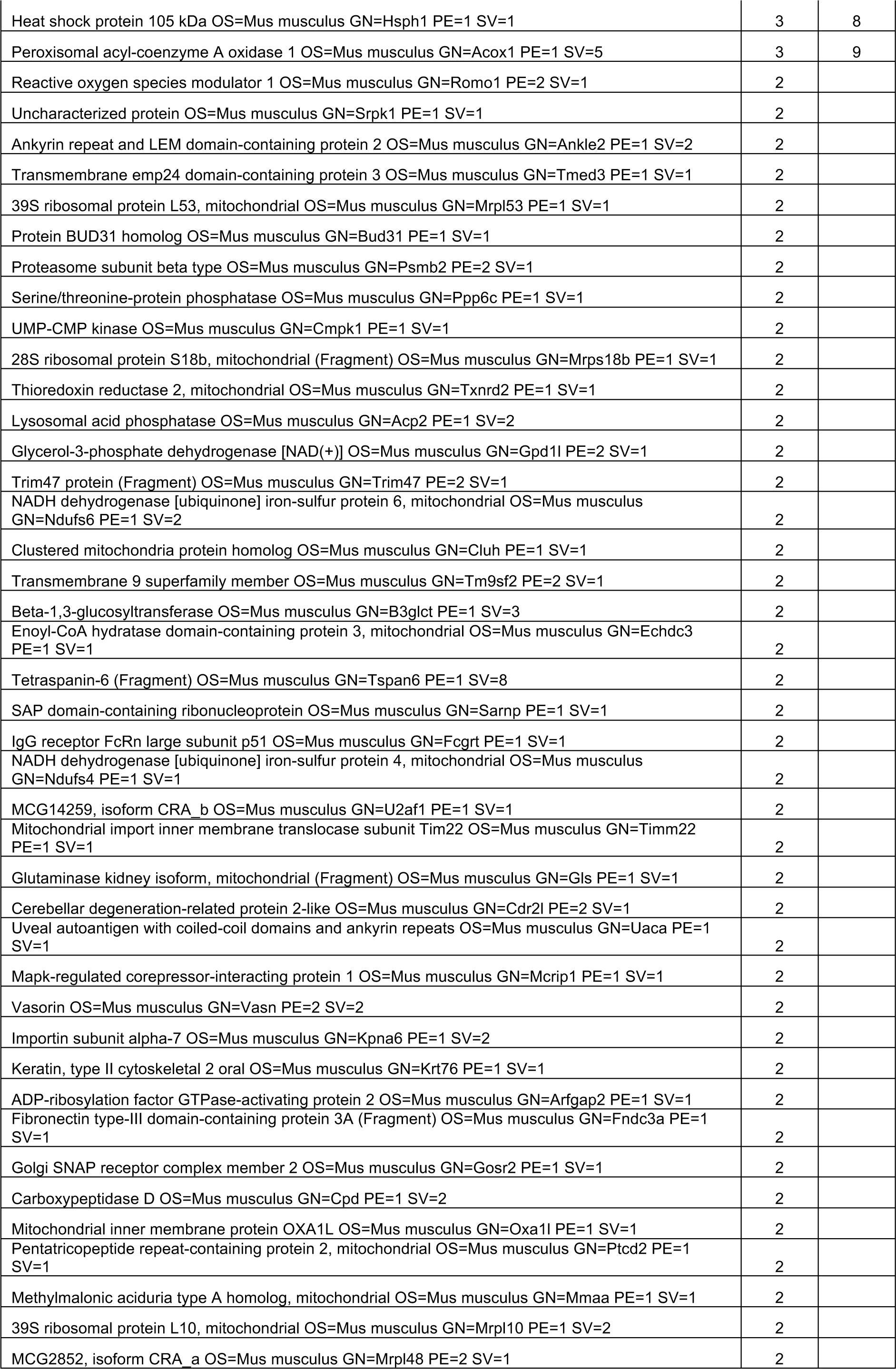
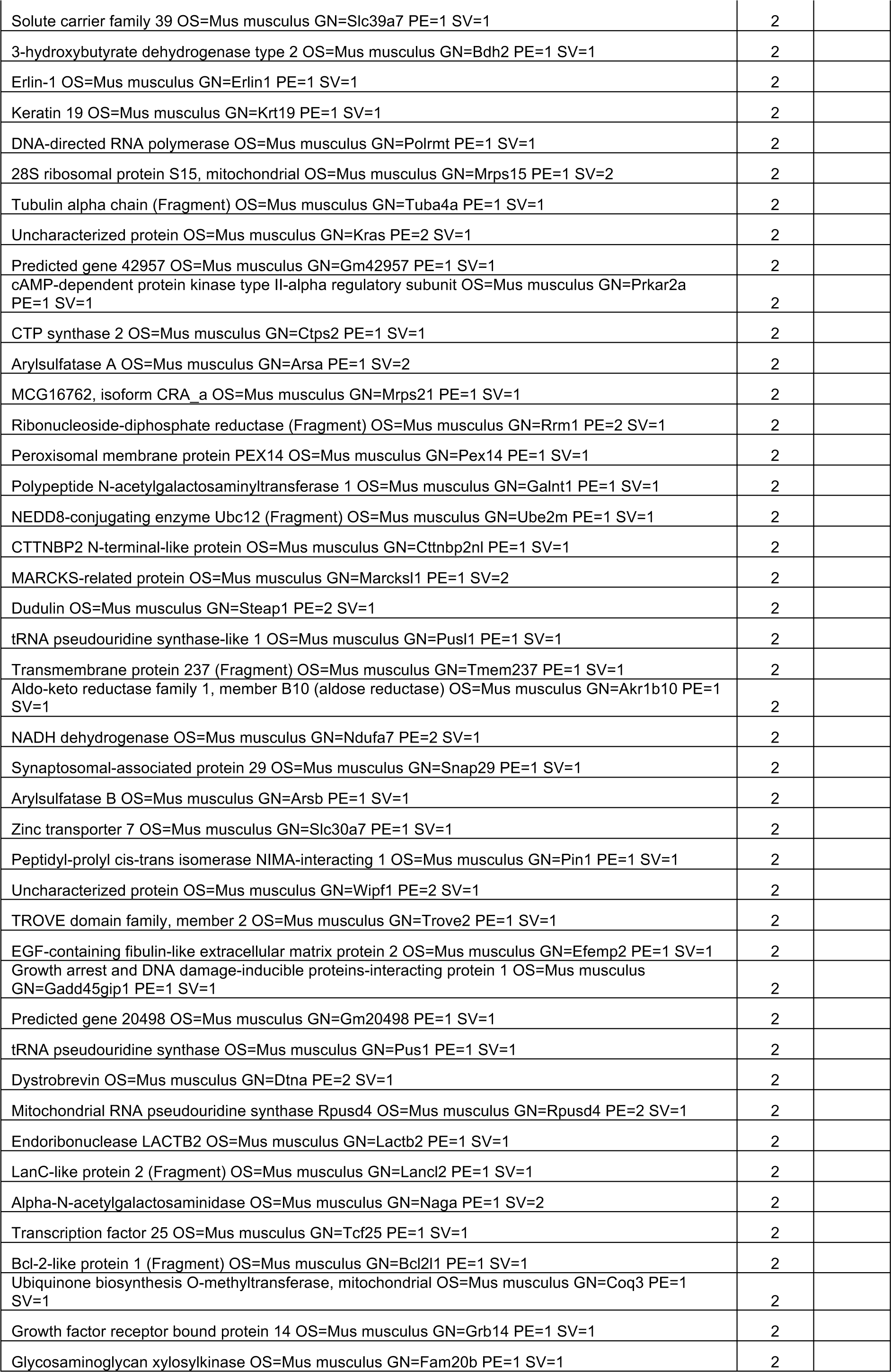
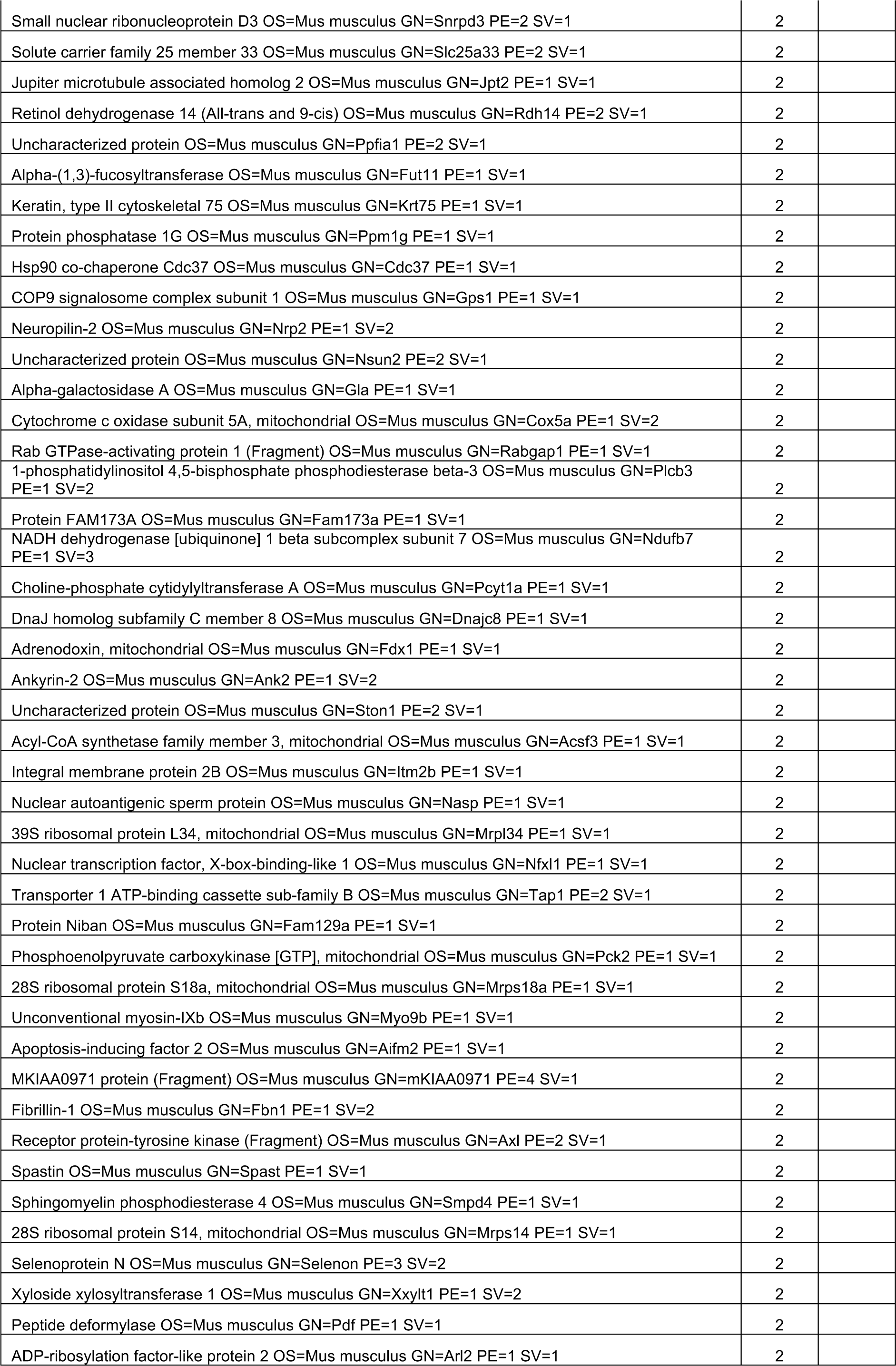
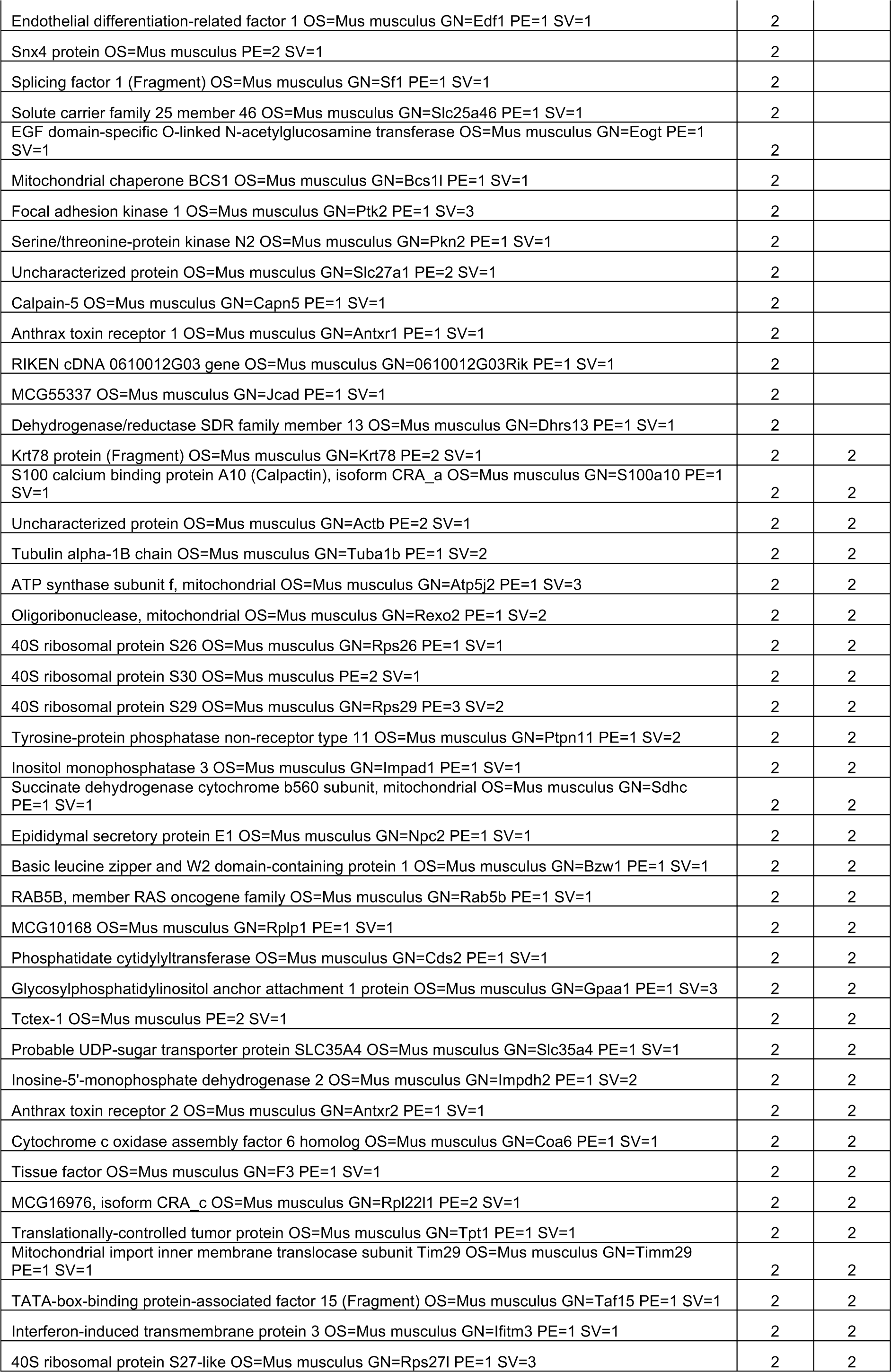
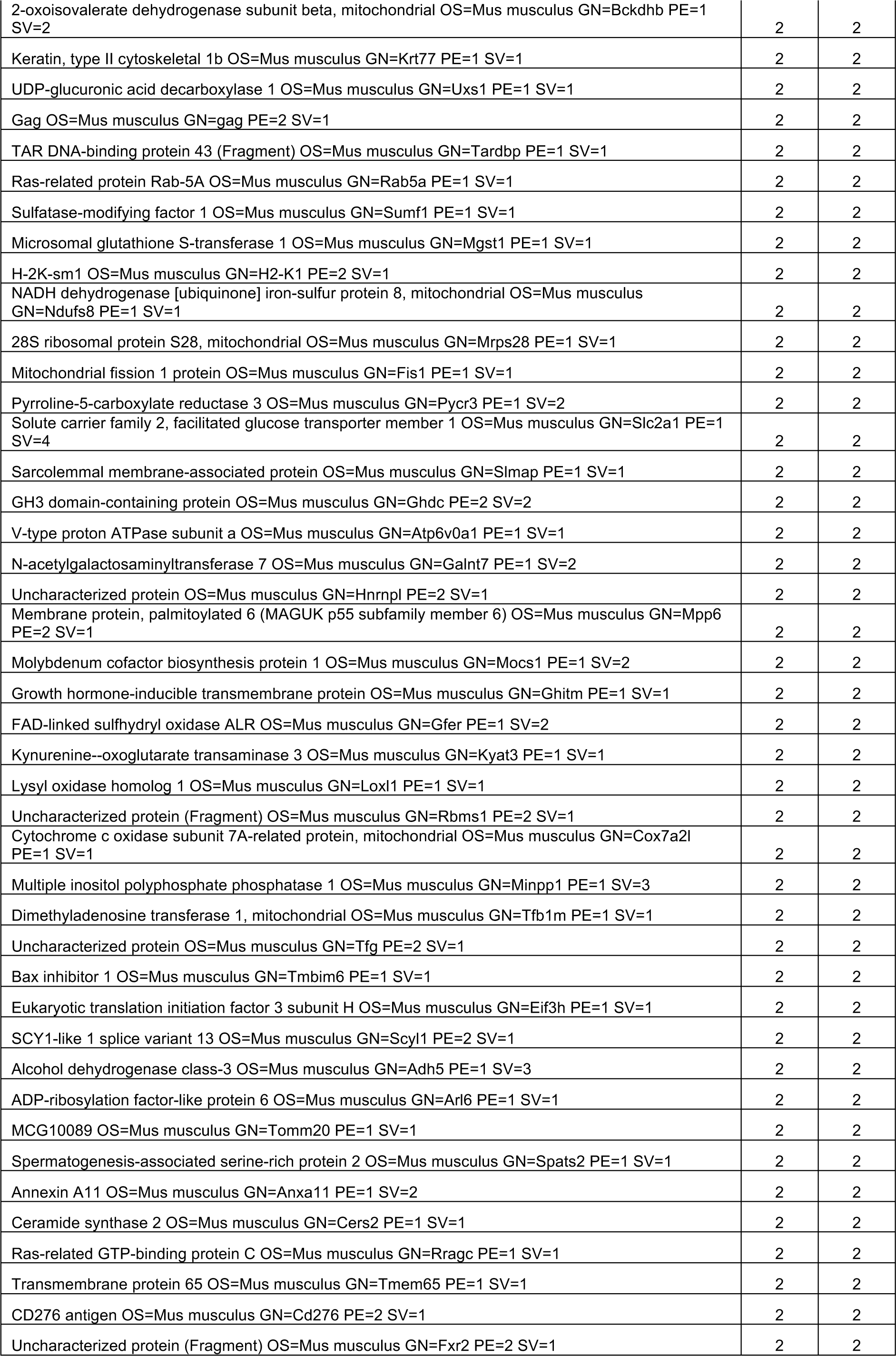
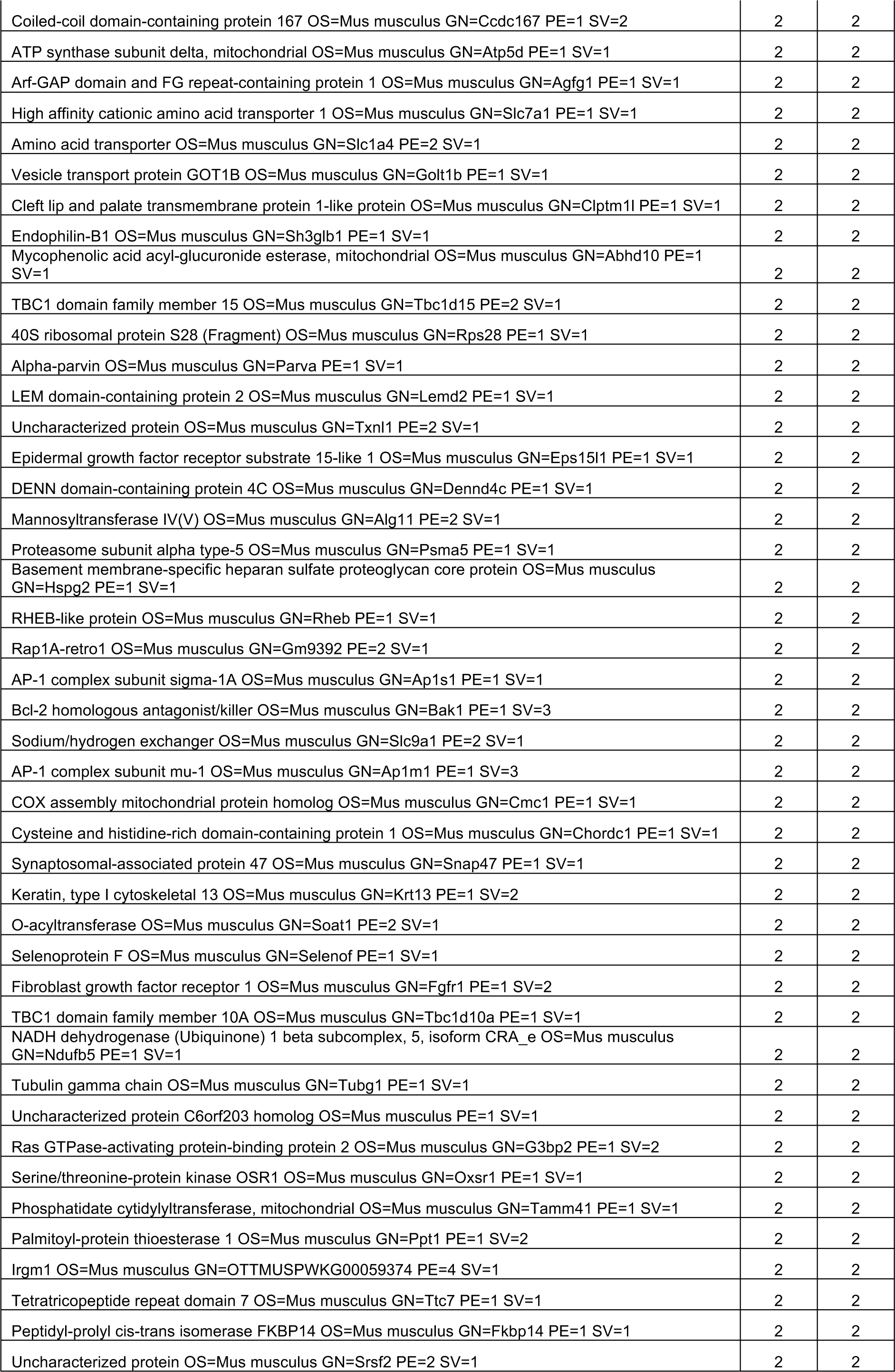
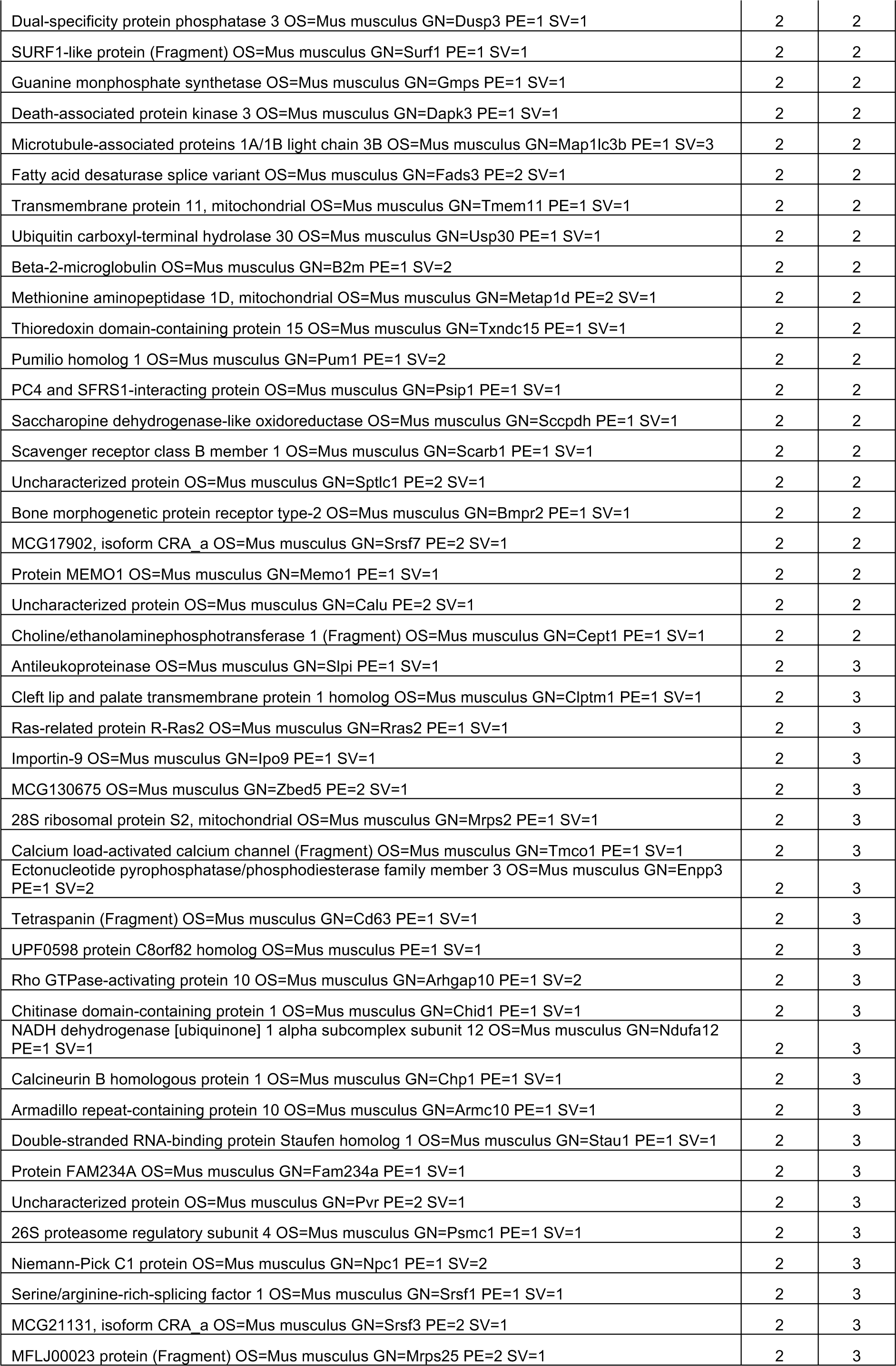
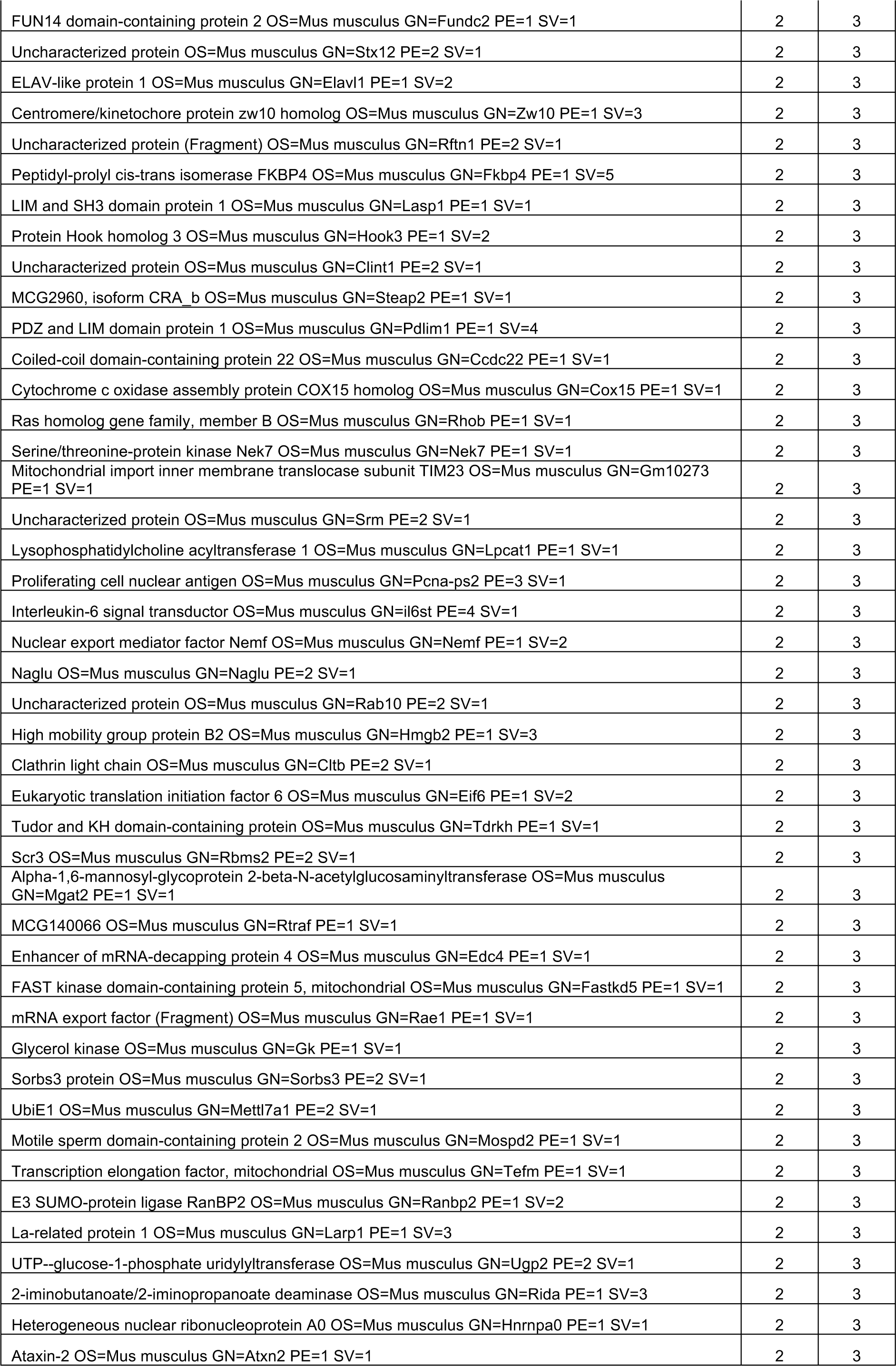
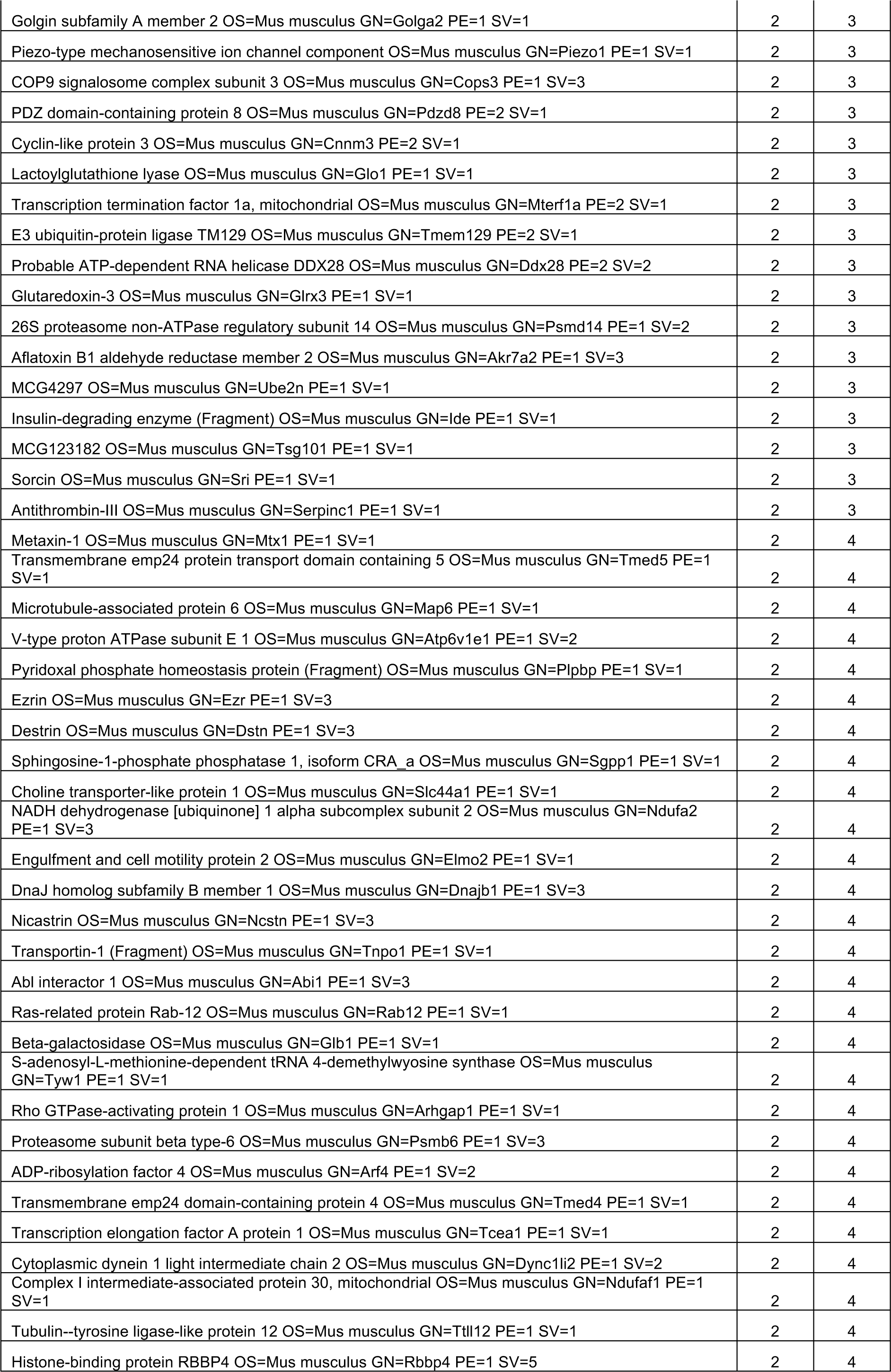
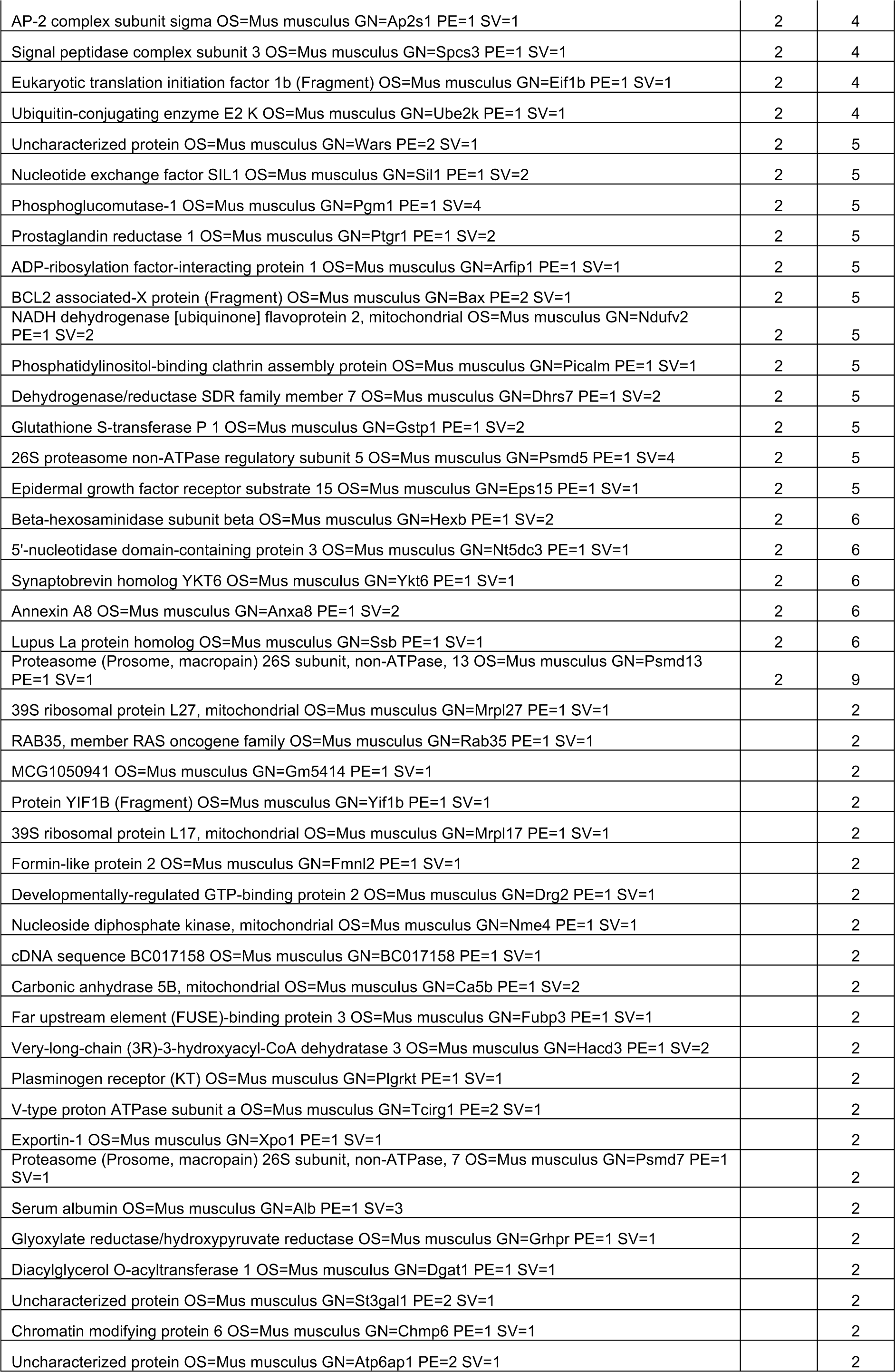
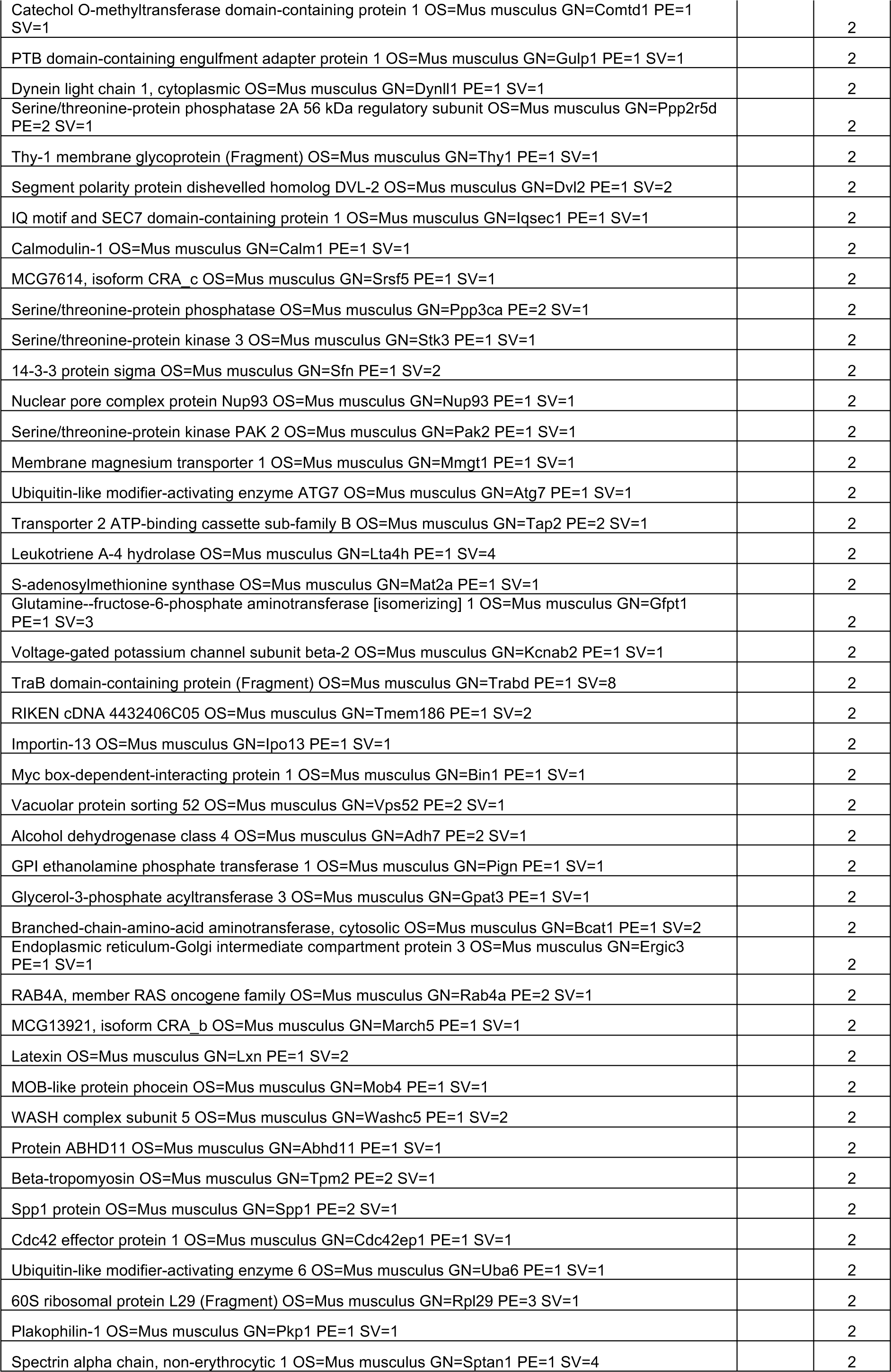
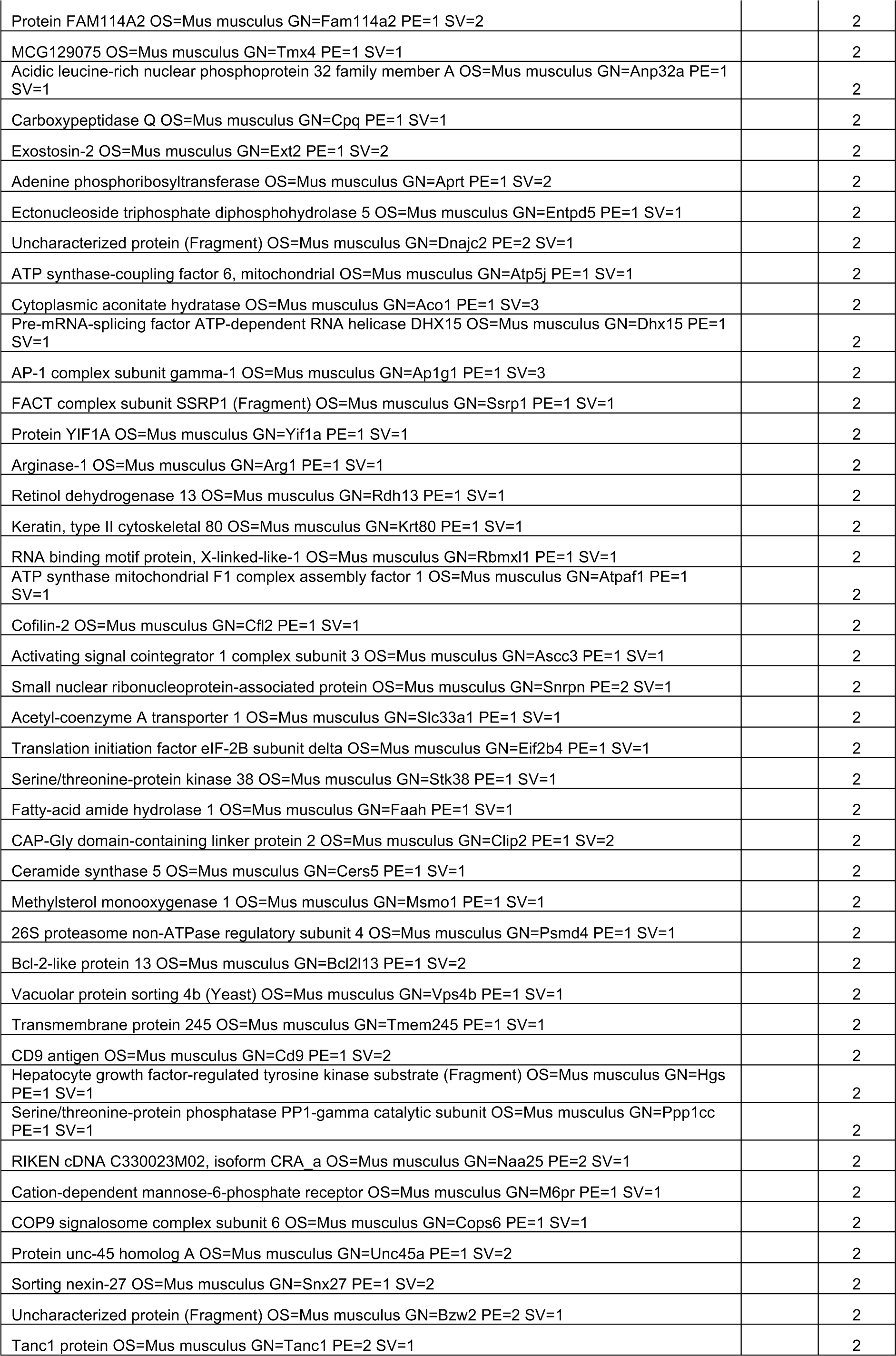
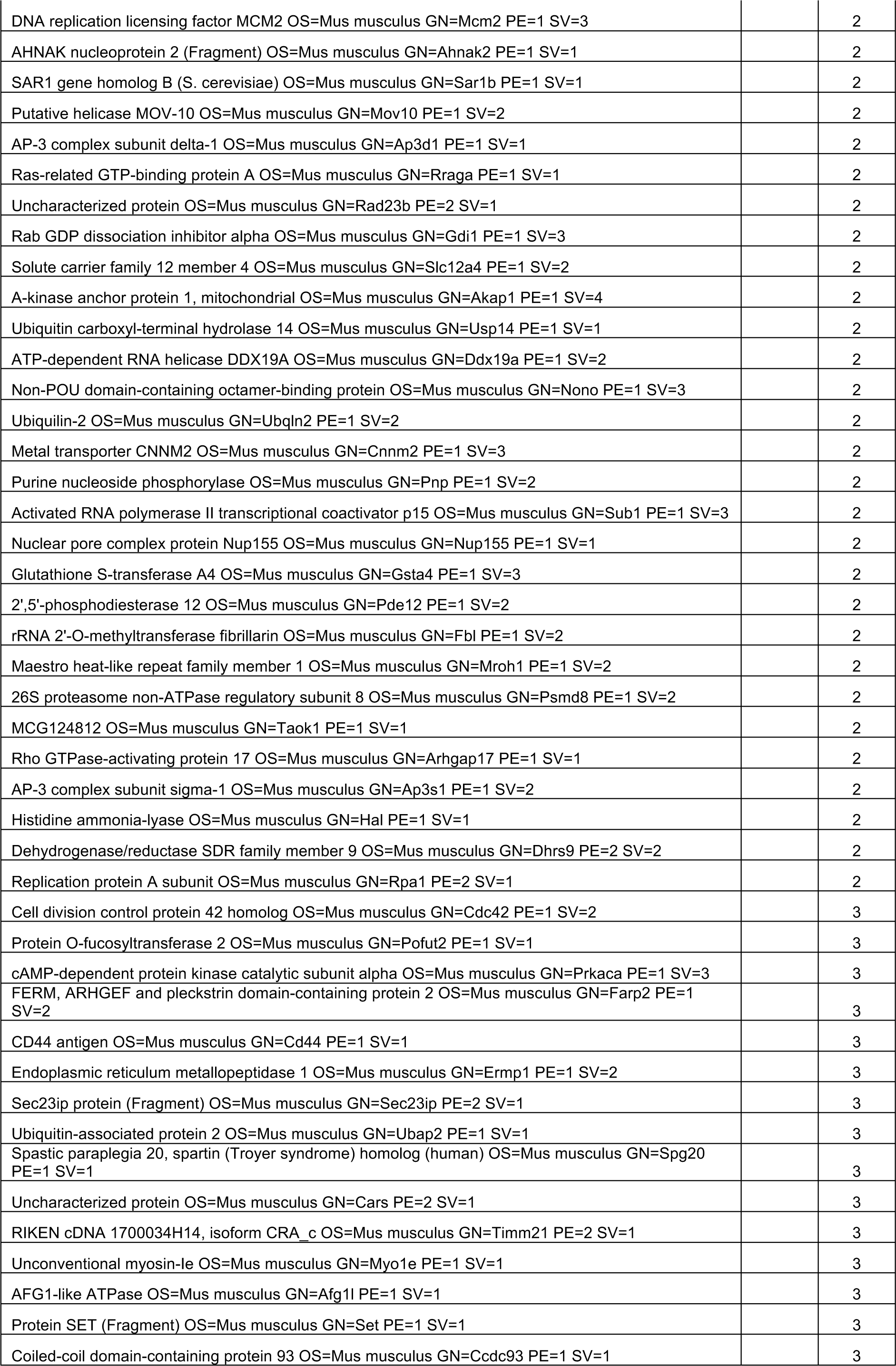
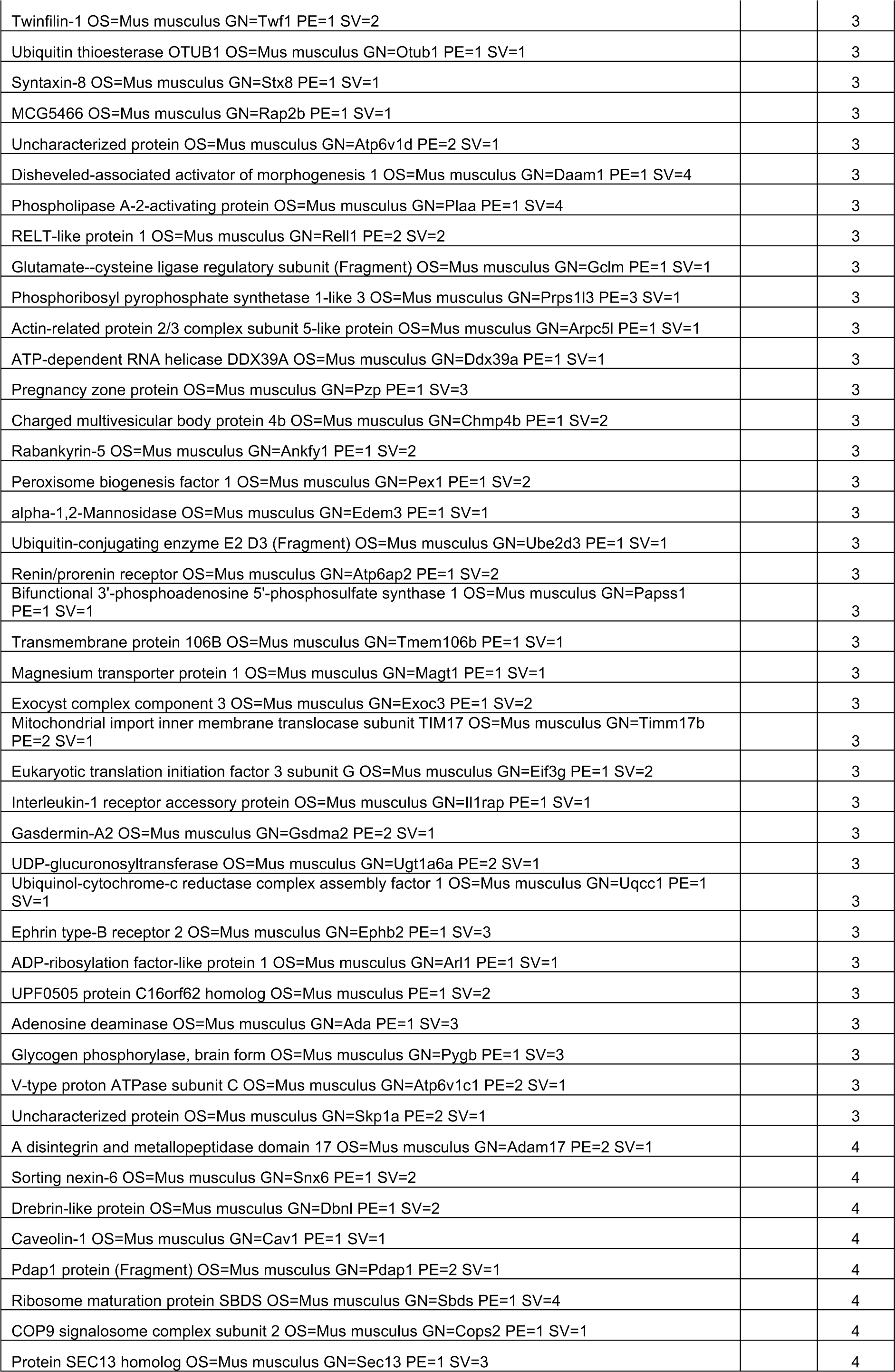
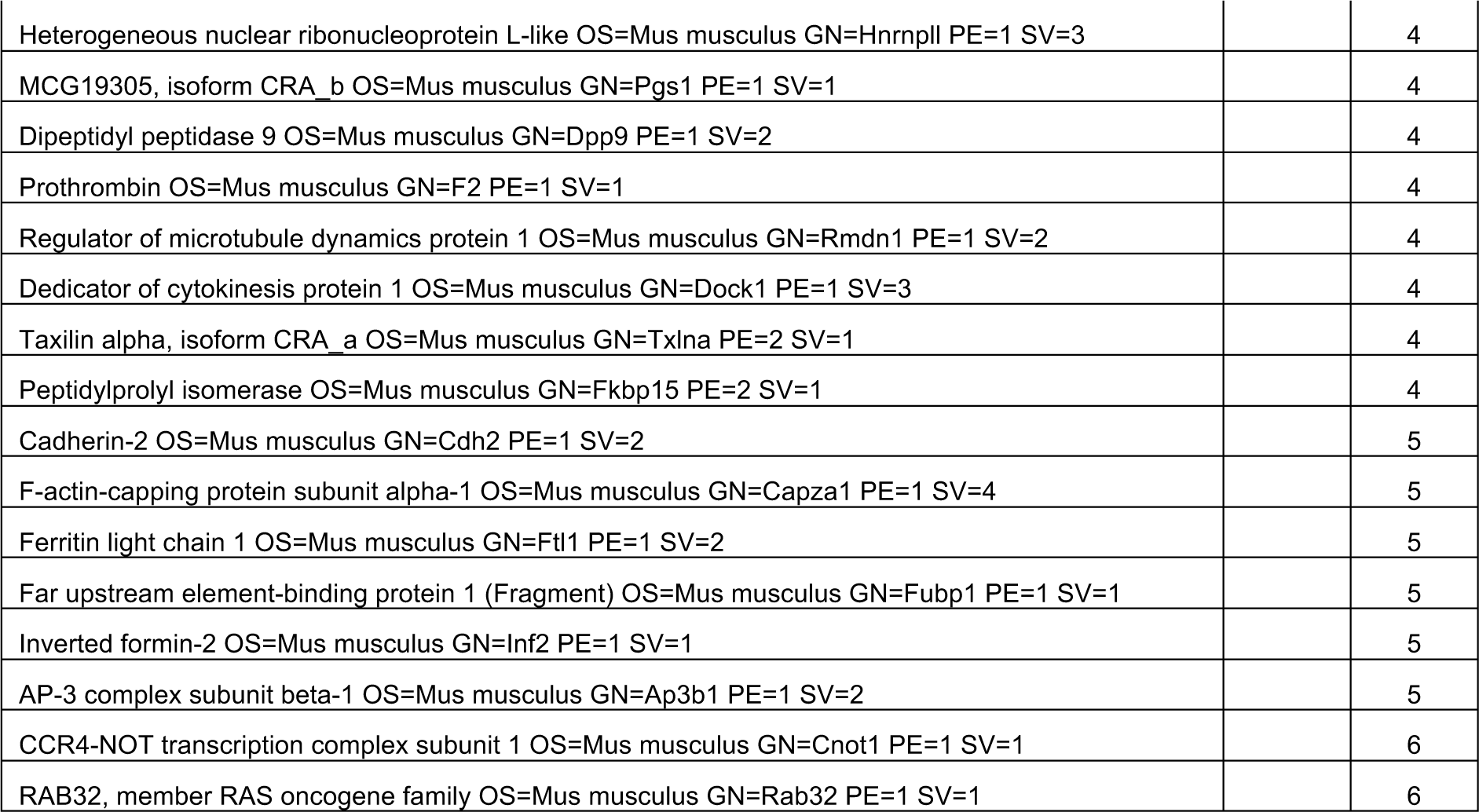
Proteins identified in lysosomal fractions.

The presence of PCI and LAMP1 in fractions 6 and 7 in the sedimentation studies, prompted us to investigate if these molecules could be co-localized using imaging approaches. Therefore, we expressed LAMP1-YFP in the Dendra2:Col1a2 cells. Dendra2-PCI was readily detected in LAMP1 positive compartments (**Fig. 7 G**). LAMP1 was also located at sites close to collagen fibrils (**Fig. S6**). Of interest, Hsp47-BFP co-localized with LAMP1-YFP (**Fig. 7H**). Co-localization of collagen-I with LAMP1 was also confirmed in both mouse and human fibroblasts (**Fig. S6**).

### Disrupted lysosome function results in defective deposition of type I collagen to the matrix

To explore further the possible role of lysosomal compartments in the deposition of type I collagen, we utilized dermal fibroblasts derived from patients with lysosomal storage disorders. Fibroblasts were isolated from patients with mucopolysaccharidosis (MPS) type I and type IIIA (MPS-I, W402X mutation in IDUA and MPSIIIA, R245H and c1284del11 mutations) mutations and lysosomal enzyme activity defects were confirmed in isolated fibroblasts (**Fig. S7**). These fibroblasts were allowed to assemble a matrix over 3 days. Detection of collagen fibrils by immune-staining using an anti-collagen I antibody identified reduced deposition of collagen fibrils in these cultures. Failed fibril nucleation events were observed in both MPSI and MPSIIIA patient derived fibroblasts (**Fig. 8A-C**, red boxes), implicating the lysosome as an essential compartment in the formation of collagen fibrils. Secretion was also assessed using CRISPR-Cas9 mediated Hibit tagging. This approach revealed unaffected PCI secretion in these fibroblasts (**Fig. 8D**). The failed fibril formation events were reminiscent of the early fibril nucleation sites seen in the Dendra2-PCI experiments shown in **Fig. 2**. In a final experiment we showed that ionomycin (a lysosome inhibitor) led to increased intracellular PCI compared to untreated cultures (**Fig. 8E**). Taken together, this final set of experiments point to a conclusion in which the lysosome, or lysosomal-like compartments, act as a store of PCI in readiness for fibril formation. When this store is perturbed, either by gene mutations in the case of lysosome storage disorders or by pharmaceutical intervention, fibril formation can be attenuated.

**Figure 8:**
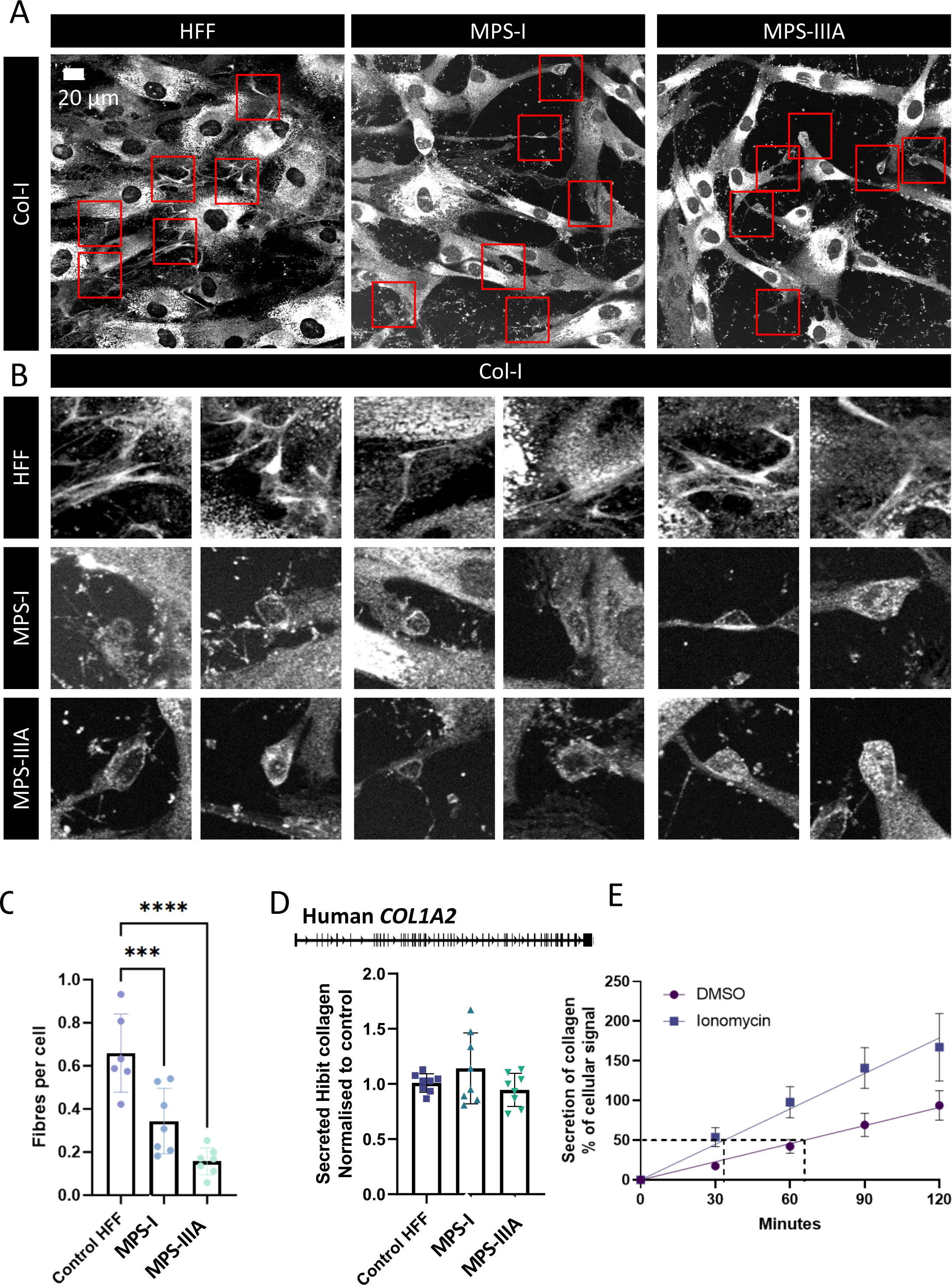
Disruption of type-I collagen fibril assembly in fibroblasts from patients with lysosome storage disorders. A) Immunofluorescence detection of type-I collagen in human skin fibroblasts taken from individuals with lysosomal storage disorders, mucopolysaccharidosis type 1 and type 3a (MPSI, and MPSIIIA). Scale bars, 20 µm. B) Regions highlighted in A show failed collagen assembly sites in MPS patient derived fibroblasts. C) Quantification of deposited collagen fibrils in control and MPS patient derived fibroblasts D) CRISPR-Cas9 mediated knock-in of the split nanoluciferase tag, Hibit, into the Exon 1 of Col1a2 in control HFF and MPS patient derived fibroblasts. Secretion rates of Hibit tagged collagen were normalized to cellular Hibit levels and indicated that there was no difference in the ability to secrete type-I collagen. E) Ionomycin triggered release of Nluc::Col1a2 from NIH3T3 cells. N=9 independent experiments, error bars represent SEM.

## DISCUSSION

In this study we established that cells control collagen fibril assembly by tethering one end of the fibril to the plasma membrane. Using a combination of gene editing to label collagen, combined with live light microscopy, we establish a fundamental distinction between collagen molecule synthesis and collagen fibril formation. We showed that collagen *molecule* secretion occurs via the conventional ER-Golgi secretory pathway at a rate of 100,000 PCI molecules/h but that collagen *fibril* assembly occurs at the plasma membrane every 24 h, via an intracellular pool. We provide evidence that collagen molecules can be captured from the extracellular space and repurposed into fibrils, via a route that is characterized by endosomal and lysosomal marker proteins. These findings help resolve a long-standing puzzle of how cells initiate a new collagen fibril and elongate one that has already been started. Our study highlights a critical distinction between collagen *molecule* secretion and collagen *fibril* assembly.

The use of CRISPR-Cas9 to introduce photoswitchable Dendra2 into PCI enabled us to distinguish newly-synthesized PCI (green) from pre-existing PCI (red, after photoswitching) within the cell. Using this approach, we showed that PCI is stored within the cell in a collection of vesicles with a t½ of 89 h. Rhythmically, every 24 h, the majority of PCI in the pool is directed to sites of fibril assembly at the plasma membrane. We considered that some of the features we had observed could be because of photobleaching. However, there was no bleed-through from the green channel into the red as the two were imaged on separate tracks on the confocal microscope with 488 and 561 excitations, respectively. Indeed, if there were any bleed-through in the time lapse images of photoswitched Dendra2-PCI this would be apparent in **Fig. 1**. Sufficiently low levels of light were used to excite the Dendra2 (at either wavelength) so as to minimise photobleaching. If there were any significant photobleaching the free radicals produced would have precluded cell survival over the 48 h of the experiment and both red and green signals from the Dendra2 would have been lost. The photoswitching of Dendra2 is not 100% efficient at the intensities used but sufficient photoswitched Dendra2 was visible to allow estimation of the half-life due to turnover and secretion. The fact that the Dendra2 signal diminished 5 times faster during fibril nucleation (e.g. see **Fig. 3**) supports the interpretation that the loss of photoswitched Dendra2 signal from the cells was predominately due to secretion rather than any significant contribution from photobleaching and protein degradation.

The spatial distribution of vesicles comprising the PCI pool did not conform with our current understanding of the ER or Golgi. Indeed, fibril formation (which draws PCI from the pool) was not inhibited by Brefeldin A or by reduction in Stx5. Instead, the inhibition of fibril formation by monensin suggested that the vesicles might share properties of LAMP1-containing lysosome-like compartments. The involvement of compartments containing PCI and lysosomal proteins, such as LAMP1, to generate fibrils, was unexpected. Non-classical routes of secretion have been described for various cargoes, but these typically carry proteins without signal peptides (for review see (Kim et al., 2018)). Some Golgi-bypassing mechanisms have been described e.g., ER targeted cargoes such as alpha integrins, the thrombopoietin receptor, and mutant forms of CFTR (reviewed by (Kim et al., 2018)). The term ‘secretory lysosome’ has been used to describe organelles that function both for degradation and storage of secretory proteins (reviewed by (Blott and Griffiths, 2002)). These secretory lysosomes are found in T cells, natural killer cells, mast cells and macrophages. Natural killer cells release toxic proteins by secretory lysosomes (reviewed by (van der Sluijs et al., 2013), and in cancer, secretory lysosomes can be a temporary store of immune checkpoint proteins (reviewed by (Wang et al., 2020)). Wounding of fibroblasts leads to the presentation of lysosomes to the cell periphery and is important to the repair of the cell membrane (Reddy et al., 2001). This presentation of lysosomes to the cell periphery is also observed during the differentiation of osteoblasts at times of enhanced collagen production (Beck et al., 2001; Nabavi et al., 2008). Thus, beyond the well documented role in protein degradation and recycling, lysosomal systems play a role in exocytosis for the purpose of cellular clearance and cell-cell communication (reviewed by (Buratta et al., 2020). In the context of collagen trafficking, the lysosome has typically been viewed as a degradative compartment via an autophagy pathway (Gorrell et al., 2021). We propose that fibroblasts use lysosomes, or lysosome-like compartments, to store and concentrate a pool of procollagen and collagen. Indeed, imaging approaches using Dendra2 showed the existence of the pool in the presence of active PCI synthesis and SILAC studies confirmed long-lived intracellular PCI with a t½ ∼ 68 h. The lysosome is an ideal storage compartment for type I collagen because the acidic pH prevents spontaneous fibril formation (Harris and Reiber, 2007) and is a stable environment for the collagen triple helix, which is highly resistant to degradation by proteinases such as pepsin. This storage function could represent a state of preparedness in the event of tissue damage or wounding. The rate of appearance of collagen in the lysosome compartment is slower than through the ER (Leblond, 1989) and others have demonstrated accumulation of collagen within the lysosome when applying high concentrations of inhibitors (Forrester et al., 2019). Recent studies following the trafficking of type I and II collagens in the presence of lysosomal inhibitors also suggest that collagen is trafficked to the lysosome (Forrester et al., 2019). Evidence from past studies suggest that not all collagen is rapidly secreted; labeling of newly synthesized collagen in rat fibroblasts appears in the lysosome at a slower rate than in the Golgi (Leblond, 1989). Together, these studies suggest that collagen may utilize this route during its exit from cells. In our present study the importance of the lysosome compartment in collagen fibril formation was demonstrated using lysosomal inhibitors and in fibroblasts derived from patients with lysosomal storage disorders, where lysosomal function is disrupted due to improper processing of glycoproteins. Importantly, these fibroblasts continue to secrete type-I collagen in the absence of fibril formation, which demonstrates the distinction between collagen secretion and fibril assembly. The defect in the ability to assemble an effective matrix warrants further investigation to understand the contribution to skeletal and connective tissue malformations associated with MPS.

Our experiments using a bioreactor showed that fibril nucleation (the start of a new fibril) and fibril propagation (the lengthening of an existing fibril) were separate processes. Under static conditions, fibrils were nucleated on the plasma membrane and these subsequently extended in length. However, when cells were put under flow, the number of fibrils on the cell surfaces did not change, whereas the fibrils were shorter. The reduction in fibril length could be attributed to either a reduction in soluble collagen from the culture medium or to a cellular response in response to flow. In the cell cultures used here, fibrils occurred exclusively on the underside of the cell where a microenvironment most probably exists to promote fibril elongation. However, it is unclear how flow across the apical surface changes the microenvironment at the basal surface. A feedback mechanism might therefore exist to divert secretion of collagen from the basal surface to the apical surface depending on how the cell senses flow. The pathway of rapid collagen egress from the cell is clearly dependent upon traffic from ER to Golgi, as described elsewhere (Bonfanti et al., 1998), therefore it remains to be understood how flow across the cell surface might affect ER-to-Golgi transport.

In conclusion, the use of CRISPR/Cas9 to tag procollagen, combined with absolute quantitation of collagen and high-resolution microscopy, has allowed visualization of collagen fibril formation by living cells. The identification of separate handling of collagen for fibril nucleation and propagation provides an explanation for how cells might regulate the number and growth of fibrils. Our study also brings together extensive data on fibril formation *in vitro* and electron microscopy of fibrils *in vivo* into one unified model of cell-regulated nucleation-and-propagation to explain how cells orchestrate tissue structure via control of collagen fibril assembly.

## ACKNOWLEDGEMENTS

KEK was supported by Wellcome (110126/Z/15/Z and 203128/Z/16/Z) and BBSRC (BB/T001984/1). JS was funded by a Biotechnology and Biological Sciences Research Council (BBSRC) David Phillips Fellowship (BB/L024551/1). BCC was supported by a Wellcome 4-year PhD studentship (210062/Z/17/Z). The Col1a1^F/F^ mice were generated with funding from NIH/NHLBI R01HL156998 and NIH/NHLBI R01HL153056. The CreER^T2^ mice were generated with funding from Crossley Barnes Studentships, University of Liverpool. The proteomics was performed at the Biological Mass Spectrometry Facility in the Faculty of Biology, Medicine and Health (University of Manchester) with the assistance of Stacey Warwood and Ronan O’Cualain, and electron microscopy was performed in the Electron Microscopy Facility, Faculty of Biology, Medicine and Health (University of Manchester). The imaging of MPS fibroblasts was with the assistance of Joan Chang. B.B. was funded by the UK MPS Society. The Bioimaging Facility microscopes used in this study were purchased with grants from BBSRC, Wellcome, MRC and the University of Manchester Strategic Fund.

## AUTHOR CONTRIBUTIONS

KEK and AP conceived the project. AA, AH, AP, BCC, CEH, DS, NH, RG, YL, OM, performed experiments and collected data. BCC performed the mathematical calculations. JS analyzed data. BB provided reagents and obtained patient consent under REC 08H101063. KKK and GB-G provided mice and advice. All authors contributed to editing the manuscript and support the conclusions. AP and KEK finalized the manuscript.

## SUPPLEMENTARY FIGURE LEGENDS

**Figure S1: Nluc-PCI and Dendra2-PCI generation**

A) Assessment of collagen secretion rate in CRISPR-edited Nluc::Col1a2 NIH3T3 cells. Nluc activity is readily detected in conditioned medium within minutes after medium change. Comparison to cellular Nluc::Col1a2 activity levels were used to assess the rate of type-I collagen secretion. N=4, individual data points are shown.

B) Paired T-test of Nluc activity at the indicated times after media change, each time is compared to T=0. Cellular Nluc activity is also shown. Data was used in Figure A.

C) CRISPR-Cas9 knock-in strategy to integrate Dendra2 encoding sequences into the *Col1a2* gene. A guide RNA immediately downstream of the signal peptide of the proa2(I) chain, directing Cas9 to exon 1 of Col1a2. A repair template encoding the Dendra2 coding sequence flanked by 800 bp homology arms.

D) PCR validation of Dendra2 integration into the *Col1a2* gene. DNA was extracted from a single cell clone identified as Dendra2 positive by FACS. Primers locations for PCR are as shown in C.

E) Real-time PCR primers were designed to detect Dendra2 knock-in, PCR products we subjected to Sanger sequencing.

F) Western blot validation of Dendra2 knock-in in the same single cell clone compared with un-edited NIH3T3.

G) The edited *Dendra2:Col1a2* locus retrained responsiveness to TGF-βI treatment and was induced transcriptionally following treatment with 2.5 ng/mL TGF-βI for 48 h, additional TGF-βI responsive transcripts were also measured. N = 3 independent repeats. Error bars represent the SEM.

**Figure S2: Rapid release of intracellular collagen at the time of fibril formation.**

A) Immunofluorescent detection of Dendra2 and collagen in *Dendra2::Col1a2* NIH3T3 cells grown for 3 days after passaging. Dendra2 positive fibrils are also positive for type-I collagen. Image captured at 1000x magnification. Scale bar represents 20 µm.

B) Airyscan microscopy of Dendra2:Col1a2 edited MC3T3 cells captured at 40x magnification, 41 h after seeding. Arrows indicate Dendra2 positive fibrils deposited by cells. Scale bar represents 20 µm.

C) Electron micrograph of MC3T3 cells and Dendra2:Col1a2 edited MC3T3 cells focusing on the collagen fibrils formed beneath the cells.

D) Quantification of cellular Dendra2:Col1a2 signal intensity in the cell body prior to the onset of fibril formation, termed nucleation, cellular Dendra2::Col1a2 signals drop dramatically during nucleation but then later recover as fibrils continue to grow. Nucleation was set at the frame prior to the first detection of Dendra2 positive fibrils, this was designated as T=0, and the cellular Dendra2 signal was set as 1, background pixel intensity was set as 0. Data from individual cells are shown, the average data is shown in Figure 2B.

E) The loss of cellular Dendra2 signal observed in Figure2C demonstrates rapid release of cellular collagen at the onset of fibril formation. The rate of loss is compared to the rate of loss of photoswitched Dendra2 signal in the absence of fibril formation (Figure 1D). n=5 individual cells, error bars represent SD.

**Figure S3: Endocytosed collagen contributes to fibril assembly.**

A) 10 µg/mL Cy3 labeled rat tail collagen was added to cultures of human lung fibroblasts for 1 or 18 h demonstrating that fibroblasts were able to endocytose type-I collagen. Uptake was significantly enhanced in cells cultured for 72 h compared to those cultured for only 24 h.

B) Quantification of Cy3 positive cells after exposure to 10 µg/mL Cy3 labeled rat tail collagen added to human foreskin fibroblasts n=3 independent experiments, error bars represent SEM.

C) Assessment of Cy3 labeled rat tail collagen uptake into SAOS2 cells for 1 h, demonstrating that these cells are able to endocytose type-I collagen with increased uptake with prolonged culture.

D) Tendon fibroblasts were fed with conditioned medium from NIH3T3 cells supplemented with varying concentrations of recombinant Nluc (rNLuc). The amount of Nluc activity taken up by tendon fibroblasts after 18 h was measured, the amount of Nluc activity taken up was directly related to the amount of rNLuc added to the medium. N=3 independent experiments, error bars show SD.

**Figure S4: Non-conventional PCI trafficking at sites of fibril nucleation.**

A) Cell viability as assessed by Prestoblue metabolism after 72 h treatment of tendon fibroblasts with increasing doses of chloroquine (Blue) and bafilomycin (purple), viability to compared to equivalent doses of DMSO (black).

B) Confocal microscopy of immunofluorescent detection of type-I collagen in DMSO, chloroquine and bafilomycin treated tendon fibroblasts, upper. Enlarged regions showing type-I collagen fibrils, lower.

C) Quantification of type-I collagen fibrils per cell, n=4 independent experiments, at least 2000 cells were scored per treatment. Error bars represent SEM.

D) Measurement of type-I collagen fibril length. *** represents p<0.001, Students T-test, unpaired. Error bars represent SD.

E) Hydroxyproline quantification of collagen in matrix deposited by tendon fibroblasts grown for 72 h with 10 µM chloroquine or 1 nM bafilomycin. N=3 independent experiments for each treatment. Error bars represent SD.

**Figure S5: Quantitative real time PCR of syntaxin-5 knock down.**

A) Quantitative real time PCR to detect Stx5 and Col1a1 transcripts in NIH3T3 cells treated with 100 pmol control scrambled (siScr) Stx5 targeting siRNA. Stx5 quantification is from n = 3 independent experiments each performed in triplicate. Error bars show SD. P value for Student’s T-test is shown.

B) Quantification of Col1a1 transcription is based on the means of N=3 independent experiments. Error bars show SEM. P value for a paired Student’s T-test is shown.

**Figure S6: Dendra2-PCI colocalizes with ER and lysosomal-like compartments**

A. On gridded 35 mm dishes Dendra2::Col1a2 3T3 cells were transfected with LAMP1-YFP overnight with Fugene 6, cells were then grown in full medium with ascorbic acid for 48 h. LAMP1-YFP positive cells were then photoswitched for 30 sec on a Nikon Eclipse microscope before imaging with Zeiss Airyscan 880 in super-resolution mode.

B. LAMP1-negative cells (Region 1) and LAMP1-positive cells (Region 2) were imaged with Zeiss Airyscan 880 in super-resolution mode. Photoswitched Dendra2-Col1a2 fibrils can be observed in Region 1 demonstrated by the detection of a red fluorescence signal. In a neighboring LAMP1-YFP positive cell, Region 2 the photoswitched Dendra2-PCI signal is located within the lumen of the lysosome. The highlighted region 2 shows the area magnified in **Figure 7G**. Scale bar represents 10 µm.

C. Co-localization of collagen-I and LAMP1 in mouse fibroblasts. Immunofluorescence imaging of type I collagen in mouse embryonic fibroblasts with the lysosome marker LAMP1. Scale bar represents 20 µm.

D. Co-localization of collagen-I and LAMP1 in human fibroblasts. Immunofluorescence imaging of type I collagen in human skin fibroblasts with the lysosome marker LAMP1. Scale bar represents 20 µm.

**Figure S7: Confirmation of mutations in MPS I and MPS IIIA fibroblasts**

A) The W402X mutation is located in exon 9 of the IDUA gene. The exon was amplified by PCR. The amplicons (243 bp) were run on a 1.5% agarose gel, at 70 V for 1 hour.

B) The amplicons were then purified and sequenced; the sequencing analysis confirmed that the MPSIH fibroblast line carries the single base substitution that introduces a stop codon at position 402 (W402X).

C) Lysosomal function was assessed in NHDF and MPS I fibroblasts by measuring IDUA activity at p.8 (n = 3 per group).

D) The R245H and c1284del11 mutations are located in exon 6 and 8, respectively, of the SGSH gene. The exons were amplified by PCR. The amplicons (224 bp) were run on a 1.5% agarose gel, at 70 V for 1 hour.

E) The amplicons were then purified and sequenced; the sequencing analysis confirmed that the MPS IIIA fibroblast line is a compound heterozygous for the R245H/c1284del11 mutation.

F) SGSH activity was also measured in NHDF and MPS IIIA fibroblasts at p.8 (n = 3 per group)

## VIDEOS AND TABLES

**Video 1: Intracellular collagen turnover over 2 days**

A Dendra2::Col1a2 edited NIH3T3 cell was photoswitched with 30 s exposure to 400 nm light. Cells were imaged every 20 minutes for 2 days. The disappearance of photoswitched collagen occurs prior to visualization of new green Dendra2-PCI. The video shows combined images captured using excitation and emission filters required to detect GFP and mCherry.

**Video 2: Deposition of Dendra2 tagged type I collagen into the matrix**

Time lapse microscopy of a Dendra2::Col1a2 edited NIH3T3 cell imaged 24 h after seeding. The cell traced over a further 24 h, with images taken every 20 minutes. Fibrils appear at approximately 36 h after seeding and then grow rapidly.

**Video 3: Dendra2 positive fibrils detected on the basal surface of cells**

Dendra2::Col1a2 NIH3T3 cell imaged after 48 h in culture. The video shows a step through of 51 Z planes starting from the basal surface. The nucleus of the cell is observed by the lack of Dendra2 signal within the cell body.

**Video 4: Photoswitched Dendra2 positive fibrils detected on the basal surface of cells**

A 3D reconstruction of images of Dendra2::Col1a2 NIH3T3 cells transfected with BFP-KDEL after photoswitching. Dendra2 positive fibrils (red/green) are formed on the basal surface of cells.

**Video 5: Photoswitched Dendra2 positive fibrils detected on the basal surface of cells**

A 3D reconstruction of images of Dendra2::Col1a2 NIH3T3 cells transfected with BFP-KDEL after photoswitching. Looped structures in Dendra2 positive fibrils (red/green) are engulfed by the cell with the endoplasmic reticulum (BFP-KDEL) arranged around the looped fibril structure.

**Video 6: Fibripositors are regularly contacted by endoplasmic reticulum and electron dense vesicles**

3view electron micrograph taken through embryonic mouse tendon showing a single fibripositor is regularly contacted by both endoplasmic reticulum and electron dense vesicles.

**Video 7: The Golgi apparatus does not contact fibripositors**

3view electron micrograph taken through embryonic mouse tendon showing that the Golgi apparatus does not contact fibripositors.

---

**Table 2:**
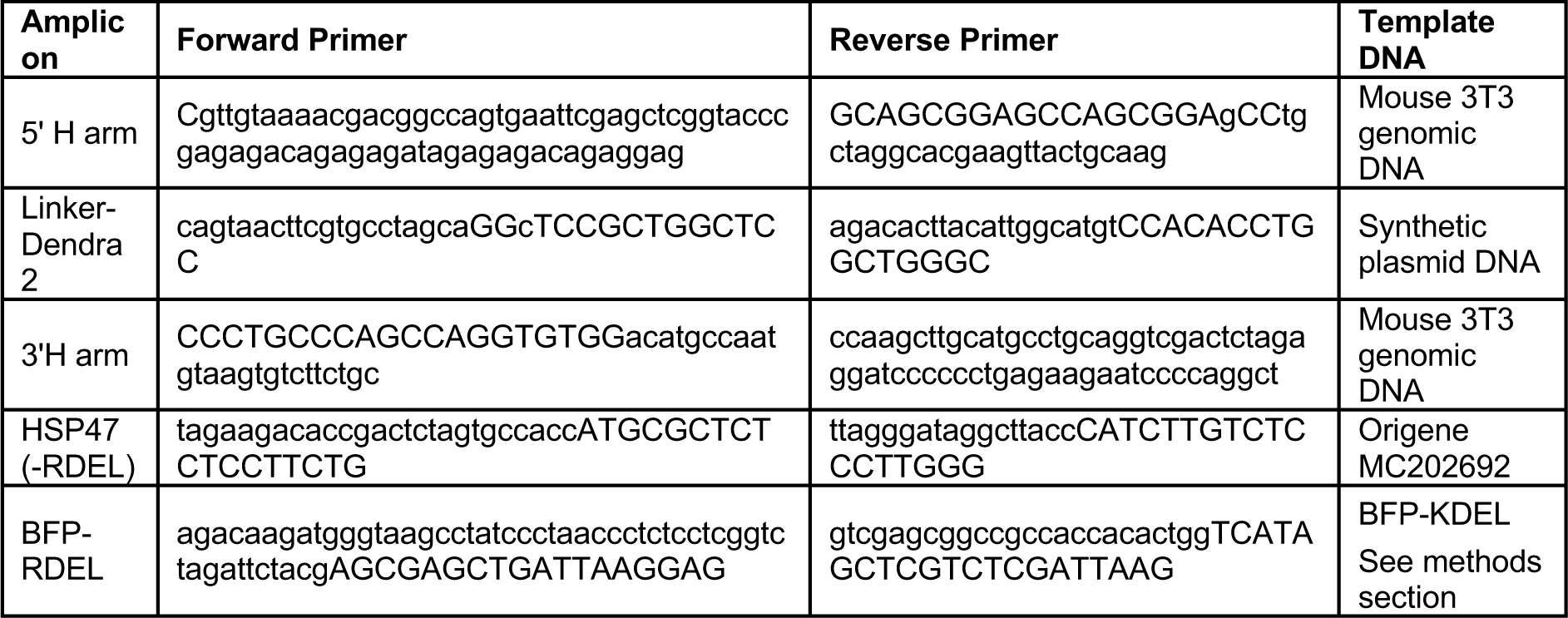
Primers used for generating repair templates and BFP tagged HSP47.

**Table 3:**
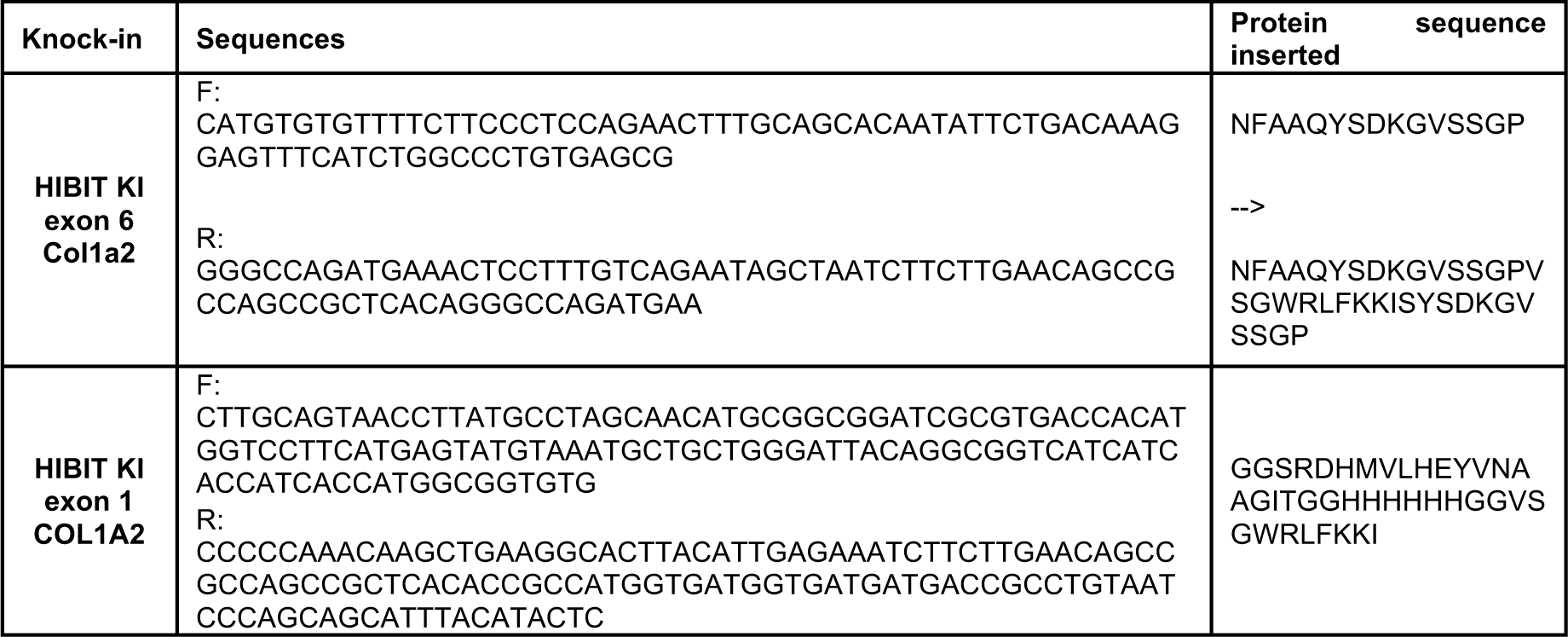
Primer sequences used to create split nanoluciferase knock-in repair templates.

**Table 4:**
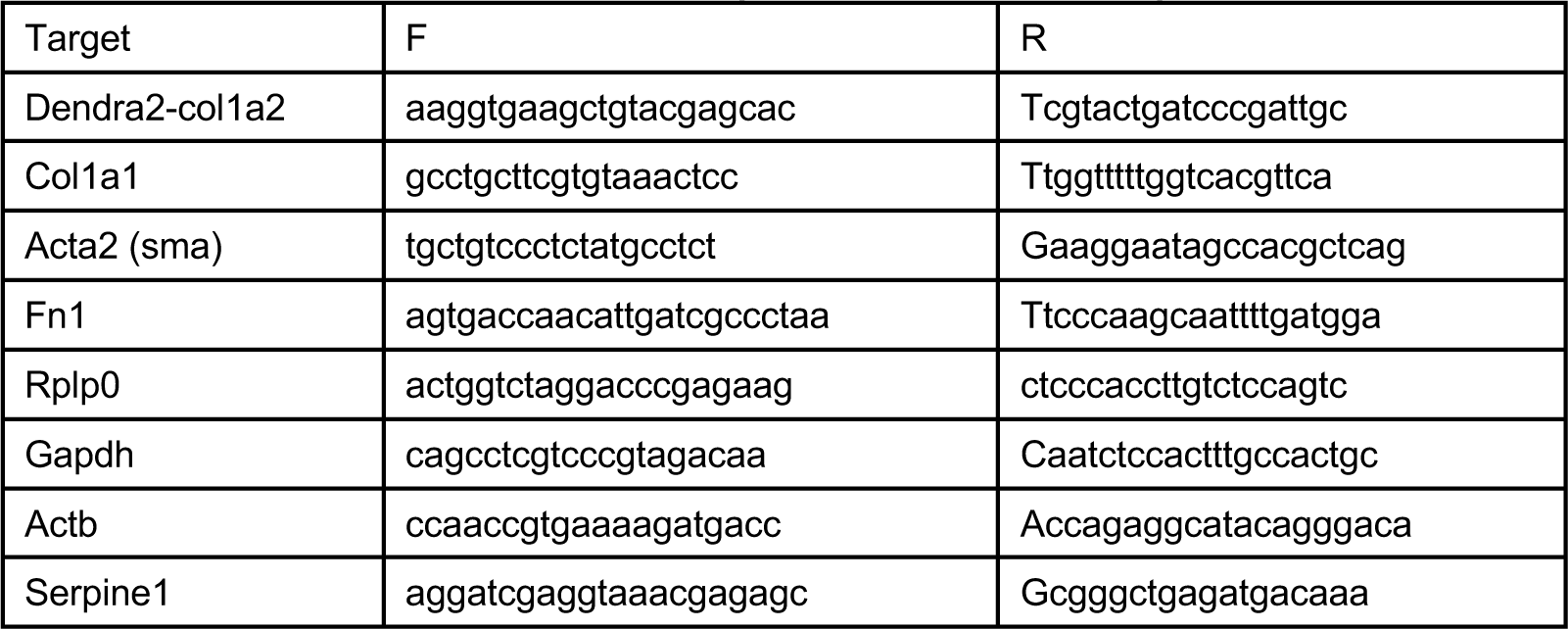
Primers used for real time PCR quantification of transcripts.

## MATERIALS AND METHODS

### Mice

The care and use of all mice in this study was carried out in accordance with the UK Home Office regulations, UK Animals (Scientific Procedures) Act of 1986 under the Home Office Licence (#70/8858 or I045CA465). The permission included the generation of conditional knock out animals. Col1a1^F/F^ mice were crossed with Col1a2-ER^T2^ mice to generate a tamoxifen-inducible Col1a2-ER^T2^:: Col1a1^F/F^ strain. Ten-week old Col1a2-ER^T2^:: Col1a1^F/F^ mice were treated with tamoxifen by intraperitoneal injections according to approved UK Home Office regulations (Project licence number I045CA465; Personal licence number P08B76E2B). Mice were humanely sacrificed by experienced personal in the animal facility.

### Cell culture and treatments

Human cells were obtained and used for this study under ethical approval REC08H101063 approved by the Northwest Research Ethical Committee, UK. Immortalized mouse embryonic fibroblasts, NIH3T3, were maintained in DMEM supplemented with 10% newborn bovine calf serum and penicillin and streptomycin. The preosteoblast cell line MC3T3 were maintained in alpha modification MEM supplemented with 10% fetal bovine serum and penicillin and streptomycin. Mouse tendon fibroblasts were cultured in DMEM:F-12 (1:1) medium supplemented with 5% L-glutamine, 10% dialyzed fetal bovine serum (FBS) and 10,000 U ml−1 penicillin/streptomycin) and at 37 °C with 5% CO2. For imaging, cells were grown on ibitreat µ-Dish (Ibidi, Germany). Where indicated NIH3T3 and MC3T3 cell lines were cultured with 20 and 200 µg/mL L-ascorbic acid, respectively, to induce collagen fibril formation. Immortalized tail tendon fibroblasts were generated as previously described (Chang et al., 2020; Pickard et al., 2019) and were maintained in DMEM/F12 1:1 mixture supplemented with 10% FBS, penicillin and streptomycin. For treatments, the following concentrations were used: bafilomycin A 100 nM, brefeldin A 100nM, monensin 1 µM, Chloroquine 1 µM and ionomycin 10 µM. Cell survival was performed using Prestoblue blue (Thermo Fisher) after 72 h treatment, fluorescence measurements were collected on the Synergy Neo2 Multi-Mode Reader (Biotek) at Excitation: 555/20 nm, Emission: 596/20 nm (Xenon flash, Lamp energy low, Gain 100 and read height 4.5 mm, 10 measurements per data point). For siRNA treatments, 200,000 cells were seeded in 6 well plates overnight and then transfected with 100 pmol siRNA,using RNAiMAX, protein, RNA and coverslips were harvested. Plasmid transfections were performed using Fugene 6 using 1 µg plasmid and 3:1 (Fugene 6:DNA) ratio. BFP-KDEL was a gift from Gia Voeltz (Addgene plasmid # 49150) (Friedman et al., 2011). HP47-BFP-RDEL was generated using PCR products of HP47 (PCR amplified from plasmid MC202692 (Origene)), and BFP-RDEL (PCR amplified from BFP-KDEL plasmid) using primers detailed In **Supplementary table 2**. These were then assembled into a pLenti CMV V5-LUC Blast (w567-1) digested with BstXI (a gift from Eric Campeau (Addgene plasmid # 21474). Cy3 labeled collagen was prepared as previously described (Doyle, 2018). Cy3 collagen was added directly to medium at the indicated concentrations and mixed immediately.

### CRISPR/Cas9 knock-in design and vector construction

We used CRISPR/Cas9 to generate the Col1A2-Dendra2 knock in cell line. Using the Sanger CRISPR design webtool we selected a guide RNA (*ACTTACATTGGCATGTTGCT* <u>AGG</u>) targeting exon 1 of the genomic sequence in order to generate a double strand break immediately after the sequence encoding the signal peptide. The guide was delivered as RNA oligos (integrated DNA technologies, Coralville, US) in complex with Cas9 protein. A double stranded DNA repair template was generated by PCR amplification of the 5’ and 3’ homology arms (800 bp each) from mouse genomic DNA. The Dendra2 mouse optimized coding sequence with a flexible linker sequence was synthesized (Genscript, US) and assembled by Gibson Assembly (NEB), primers are shown in **Supplementary table 2**. Knockin of nanoluciferase to Col1a2 was previously described (Calverley et al., 2020).

To knock-in split nanoluciferase (Hibit) into mouse *Col1a2* exon 6 we used gRNA: *GCTGCTCAGTATTCTGACAA*. For human *COL1A2* exon 1 knockin of Hibit we used gRNA: *CAAGCTGAAGGCACTTACAT*. Repair templates for CRISPR-mediated knock-in of Hibit are detailed in **Supplementary table 2**. Briefly the two primer sequences were annealed to generate hybrid ssDNA:dsDNA repair templates with 29 bp homology arms.

### CRISPR/Cas9 Delivery

For knock-in of Dendra2, NIH3T3 and MC3T3 cells were seeded 24 h prior to transfection of the repair template, using Fugene 6 (Promega) at a ratio of 3 µL per 1 µg DNA, cells were maintained in transfection mixture overnight and then fresh medium was added for approximately 6 h. Cells were then transfected overnight with 100 pmol of tracrRNA and crRNA complexed with 4 pmol recombinant cas9 protein using RNAiMAX (Thermo Fisher). Medium was then replaced and cells were grown for 48 h before assessing fluorescence and sorting.

For Hibit knockin crRNA were purchased from IDT, sequences are shown in **Supplementary table 3**. 0.25 µL crRNA (100 µM) was annealed with equimolar amounts tracRNA (IDT), and complexed with Cas9 (1.5 µL 1 µM, NEB). CRISPR/Cas9 complexes and 100 pmol annealed repair template were electroporated (Neon transfection system, Thermo). Electroporation conditions for NIH3T3 were 1400 V, 20 ms, 2 pulses using 10 µL tips and 50,000 cells and for HFFs were 1700 V, 30 ms and 1 pulse using 10 µL tips and 100,000 cells. Electroporated cells were grown in 12 well plates.

### CRISPR/Cas9 validation

Dendra2 positive populations were sorted by FACS and grown to sufficient numbers for validation of editing. Cells then sorted into single cells clones. All assays and imaging were performed on single cell sorted populations. Individual cells were then grown to sufficient numbers for validation. Primers for validation of the knockin into genomic DNA (PR1 and PR2) were as follows: F-GGCAAGGGCGAGAGAGG and R-TTTTCTCCGACAGATTAGAGGGC. Real time validation of Dendra2::Col1a2 transcripts were assessed in single cell clones treated with 2.5 ng/mL TGF-b1 for 48 h using the primers indicated in **Supplementary table 4**, real time PCR was normalized to the geometric mean of Rplp0, Gapdh and Actb. The sequence of the dendra2-col1a2 transcript was then validated by Sanger sequencing. Western blotting was performed on 4-12% tris-glycine gels with 25 µg cell lysate. Antibodies used in this study were Collagen (Gentaur, OARA02579, dilution 1:2000 (WB), 1:500 (IF)), Dendra2 (Origene, TA180094, dilution 1:500), GAPDH (Sigma, G8795, dilution 1:10,000) Calreticulin (Stressgen; SPA-601, dilution 1:1000), Lamp1 (Santa Cruz, sc-20011, 1:500), Vincullin (Chemicon; CBL233, 1:2000), Hp47 (Santa Cruz, sc-398579, 1:1000 (WB), 1:500 (IF)), Syntaxin 5 (Santa Cruz, sc-365124, 1:500(WB)) PDI (Abcam, ab180993, dilution 1:500 (IF)).

### Imaging

For immunofluorescence detection of cellular and extracellular proteins, cells grown on coverslips were fixed with 4% paraformaldehyde for 20 minutes at room temperature and washed with PBS before permeabilization with 0.2% triton-x-100 in 10%FBS for 15 minutes. Antibodies used in this study are detailed above.

Figures 1C, **E and F, Video 1,** Figures 2B were acquired on an Eclipse Ti inverted microscope (Nikon) using a 60x objective, the Nikon filter sets for GFP and mCherry and LED (Lumencor) fluorescent light sources each with 300 ms exposure. Photoswitching was performed using a 30s exposure to UV LED light source (400 nm). The images were collected using a Retiga R6 (Q-Imaging) camera, and captured using NIS Elements AR.46.00.0 software. Pixel intensity was analyzed using FIJI ImageJ (http://imagej.net/Fiji/Downloads). Cells were maintained at 37°C and 5% CO_2_.

Images from **Supplementary** figures 2A and **5C** were collected on a Zeiss Axioimager D2 upright microscope using a 100x objective and captured using a Coolsnap HQ2 camera (Photometrics) through Micromanager software v1.4.23. Following 4% PFA fixation cells were permeabilized and stained with antibodies to Dendra2 (dilution 1:100) and collagen (dilution 1:200), secondary antibodies used were goat-anti-mouse488 (Cell Signaling; 4408, dilution 1:400) and goat anti-rabbit-Cy5 (Invitrogen; A10523, dilution 1:400). Specific band pass filter sets for FITC and Cy5 were used to prevent bleed through from one channel to the next. All images were then processed and analyzed using Fiji.

Images for Figurse 2A, 3A, 3B, 4F, 5B, 6D, 6E, 6H, 7A, 7G, 7H. Supplementary Figures 2B, 4B, 6A, 6B, Videos 2-5 were acquired with either a Fluar 20X 0.75NA objective at 1024x1024 resolution, or 100x using a Zeiss LSM880 equipped with Airyscan detector set to super-resolution mode. Green fluorescence was excited at 488nm and collected through a 495-550nm filter. The 32 phase images were recombined using the Airyscan processing tool in the Zeiss Zen 2 software and the image brightness and contrast adjusted using BestFit.

Figures 5A, **7C, 9A** and **B** were collected on a Leica TCS SP5 AOBS inverted confocal using an [63x / 0.50 Plan Fluotar] objective. The confocal settings were as follows, pinhole [1 airy unit], scan speed [1000 Hz unidirectional]. Images were collected using [PMT] detectors with the following detection mirror settings; [FITC 494-530 nm; Cy5 640-690 nm] using the [488 nm (20%) and 633 nm (25%)] laser lines respectively. To eliminate crosstalk between channels images were collected sequentially.

### Electron microscopy

Transmission electron microscopy: Dendra2::Col1a2 edited MC3T3 were grown for 7 days in the presence of 200 µg/mL L-ascorbic acid, replenishing medium and ascorbic acid every 2 days. Following fixation in 2.5% glutaraldehyde/100 mM phosphate buffer (pH 7.2) for 30 minutes at RT cells and matrix were scraped from the culture dish. After 2 h at 4 °C the samples were washed in 100 mM phosphate buffer (pH 7.2). After immersion in 2% osmium/1.5% potassium ferro-cyanide in cacodylate buffer (pH 7.2) for 1 h at RT, samples were washed in ddH2O, and fixed in 1% tannic acid/0.1 M cacodylate buffer (pH 7.2) for 2 h at 4 °C. Samples were then thoroughly washed in ddH2O and incubated in 2% osmium tetroxide/ddH2O for 40 min at RT, before washing again in ddH2O at RT. This was followed by a final incubation step at 4 °C in 1% uranyl acetate (aqueous) overnight. Samples were then washed before infiltrated with a series of propylene oxide and TAAB 812 resin kit mix, with increasing resin concentration (2 h in 30% resin, 2 h in 50% resin, 2 h in 75% resin, 3 x 1 h in 100% resin). Samples are then embedded in capsules and cured at 60 °C for 12 h. Sections (80 nm thick) were cut from the sample blocks and examined using a Tecnai 12 BioTwin electron microscope.

Serial block face SEM: Tendons were prepared as described previously (Starborg et al., 2013). In brief, tendons were fixed in 1% osmium and 1.5% potassium ferrocyanide in 0.1 M sodium cacodylate buffer for 1 h, washed with ddH_2_O water, then incubated with 1% tannic acid in 0.1 M cacodylate buffer for 1 h, washed, then incubated with 1% osmium tetroxide in water for 30 min. Samples were then washed with ddH_2_O water and stained with 1% uranyl acetate in water for 1 h, then dehydrated in acetone and embedded into resin.

ImmunoEM: Post-embed labeling was used to detect type I collagen using a rabbit anti–chicken collagen-I antibody (Biodesign International) at a dilution of 1:500 followed by a gold-conjugated goat anti–rabbit antibody (British Biocell International) at a dilution of 1:200. All sections were subsequently stained with uranyl acetate and examined using a Tecnai 12 BioTwin electron microscope.

### Atomic force microscopy

Atomic force microscopy was performed using a JPK NanoWizard IV (Bruker Nano Inc., Karlsruhle, Germany, previously JPK Instruments AG, Berlin, Germany) mounted on a Zeiss AX10 (Carl Zeiss Microscopy GmbH, Jena, Germany) inverted light microscope operating under JPK NanoWizard Control software (V 6.1.65). Images were captured using NuSense Scout 350R cantilevers (NuNano, Bristol, UK) with nominal spring constant, frequency and tip radius of 42 N/m, 350kHz and <10nm respectively. Height data was processed using JPK Data Processing software (V 6.1.65), and was 1^st^ order flattened prior to analysis.

### Nanoluciferase activity assay

Conditioned medium collected after washing edited cells twice with PBS, was placed in white walled 96 well plates (Nunc MicroWell 96-Well, Nunclon Delta-Treated, Flat-Bottom Microplate, Thermo Fisher Scientific, Paisley, UK# 136101) To assay Nluc activity 0.5 µL of coelenterazine (final concentration 3 µM) was added immediately prior to measurement. Light production was measured using filter cubes #114 and #3 on the Synergy Neo2 Multi-Mode Reader (Biotek), readings for each well were integrated over 200 ms with 4 replicate measurements per well (Gain 135 and read height 6 mm). For assessment of cellular and matrix derived Nluc activity cultures were decellularized by addition of extraction buffer (20 mM NH_4_OH, 0.5% Triton X-100 in PBS) for 2 mins at 37 °C until no intact cell is visible under light microscope, this cellular fraction was removed and the matrix was washed with PBS, and then scraped into 1 mL PBS and pelleted by centrifugation at 12,000 x *g* for 5 mins. Matrix was resuspended in 100 µL extraction buffer before assessing Nluc activity.

### Hydroxyproline Assay

Decellularized cultures were scraped and pelleted into 1.5 mL tubes before freezing at -20 °C for the hydroxyproline quantitation. Hydroxyproline was measured using methods previously described (Reddy and Enwemeka, 1996). Briefly, 100 µL 6M HCl was added to the pellet and incubated at 100 °C overnight. Samples were cooled to room temperature and spun at 12,000 *xg* for 3 mins to remove residual charcoal. For each sample (50 μL) was mixed with chloramine T (450 μL) and incubated at room temperature for 25 mins. Ehrlich’s reagent (500 μL) was added to each sample and incubated at 65 °C for 10 mins. All samples were compared to hydroxyproline standards treated identically. The absorbance of 100 μL was measured a 96-well plate and absorbance at 558 nm read on a H1 plate reader (Biotek).

### Lysosomal fractionation

Cells (30 million) were grown in 150 mm tissue culture dishes, trypsinized and pelleted at 1000 *xg* for 5 minutes. Lysosomal fractions were prepared using the Lysosome Isolation Kit (Sigma Aldrich, LYSISO1) according to manufacturer’s procedures. Nine fractions were collected after ultracentrifugation.

### Proteomic sample preparation

Cell pellets were resuspended in 30 µl SL-DOC (1.1% sodium laurate, 0.3% sodium deoxycholate in 25 mM ammonium bicarbonate supplemented with protease and phosphatase inhibitor cocktails). Six 1.6mm steel beads were added and the samples were homogenized in a Bullet Blender Tissue Homogeniser. A BCA was done to quantify the amount of protein in each sample. Each sample (50 µg) was made up to 5% SDS and then reduced and alkylated with DTT and Iodoacetamide (IAA) respectively. For lysosome fractions, samples were processed without further extraction procedures. Samples were acidified using H_3_PO_4_ and S-trap binding buffer (90% Methanol in 100 mM TEAB pH 7.1) was added. Samples were loaded, onto S-Trap columns (ProtiFi) and washed with S-trap binding buffer 4 times. Proteins were digested with 0.8 µg/µL trypsin solution (proteomics grade trypsin, Promega). Peptides were then eluted in 65 µl Digestion buffer (50 mM TEAB pH 8.5), 65 µL 0.1% formic acid (in water) and finally with 30 µL 0.1% formic acid, 30% Acetonitrile (ACN) (in water). Samples were then desalted using Oligo R3 resin beads in a 96-well, 0.2 µm PVDF filter plate (Corning). The beads were washed and then samples added and washed twice with 0.1% formic acid. Samples were then eluted in 0.1% formic acid in 30% ACN and lyophilized using a speed-vac (Heto Cooling System).

### Mass Spectrometry

Dried peptides were resuspended in 10 µl 0.1% formic acid in 5% ACN. Samples were analyzed using an ultiMate® 3000 Rapid Separation LC system (RSLC, Dionex Corporation) coupled to first a Orbitrap Elite, for quality control, and then a Q Exactive HF Mass Spectrometer (Thermo Fisher). For both, mobile phase A was 0.1% formic acid in water and B was 0.1% formic acid in ACN. The Orbitrap used a 75 mm x 250 um inner diameter 1.7 µM CSH C18 analytical column (Waters) with a gradient from 92% A and 8% B to 33% B in 10 minutes at a rate of 300 nL/min. The Q Exactive used a gradient of 95% A and 5% B to 18% B at 58 minutes, 27% at 72 min and 60% at 74 min with the same flow rate and a 75 mm x 250 µm inner diameter CSH C18 analytical column (Waters). Peptides were selected by DDA for fragmentation automatically and data was acquired for 90 min in positive mode.

### SILAC labeling

Mouse tendon fibroblasts were pulsed with 100 mg/L C^13^ (“Heavy”) lysine for 48 h and then washed, trypsinized, lysed and processed for mass spectrometry (see below). Mass spectrometry results files were exported into Proteome Discoverer for identification and quantification using a SILAC 1plex (Lys6) method. All searches included the fixed modification for carbamidomethylation on cysteine residues resulting from IAA treatment to prevent cysteine bonding. The variable modifications included in the search were oxidized methionine (monoisotopic mass change, +15.955 Da) and phosphorylation of threonine, serine and tyrosine (79.966 Da). A maximum of 2 missed cleavages per peptide was allowed. The minimum precursor mass was set to 350 Da with a maximum of 5000 Da. Precursor mass tolerance was set to 10 ppm, fragment mass tolerance was 0.02 Da and minimum peptide length was 6. Peptides were searched against the Swissprot database using Sequest HT with a maximum false discovery rate of 1%.

Half-lives were calculated from heavy to light ratios (HL) as shown by Schwanhäusser using the following equations (Schwanhausser et al., 2009):

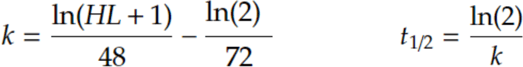

Where 48 represents the SILAC pulse time, 72 is the calculated cell doubling rate and k is the rate constant of protein decay, which is then used to calculate the half-life in the second equation.

This calculation assumes that no new light protein is produced and that the amount of light protein decays exponentially over time. The total amount of protein is assumed to double per complete cell cycle.

Proteins were only selected for half-life calculation if at least 3 peptides were detected, where at least one is heavy, across 2 repeats.

