## Supplementary figures and images for "Collagen fibril formation at the plasma membrane occurs independently from collagen secretion"

### Supplementary figures

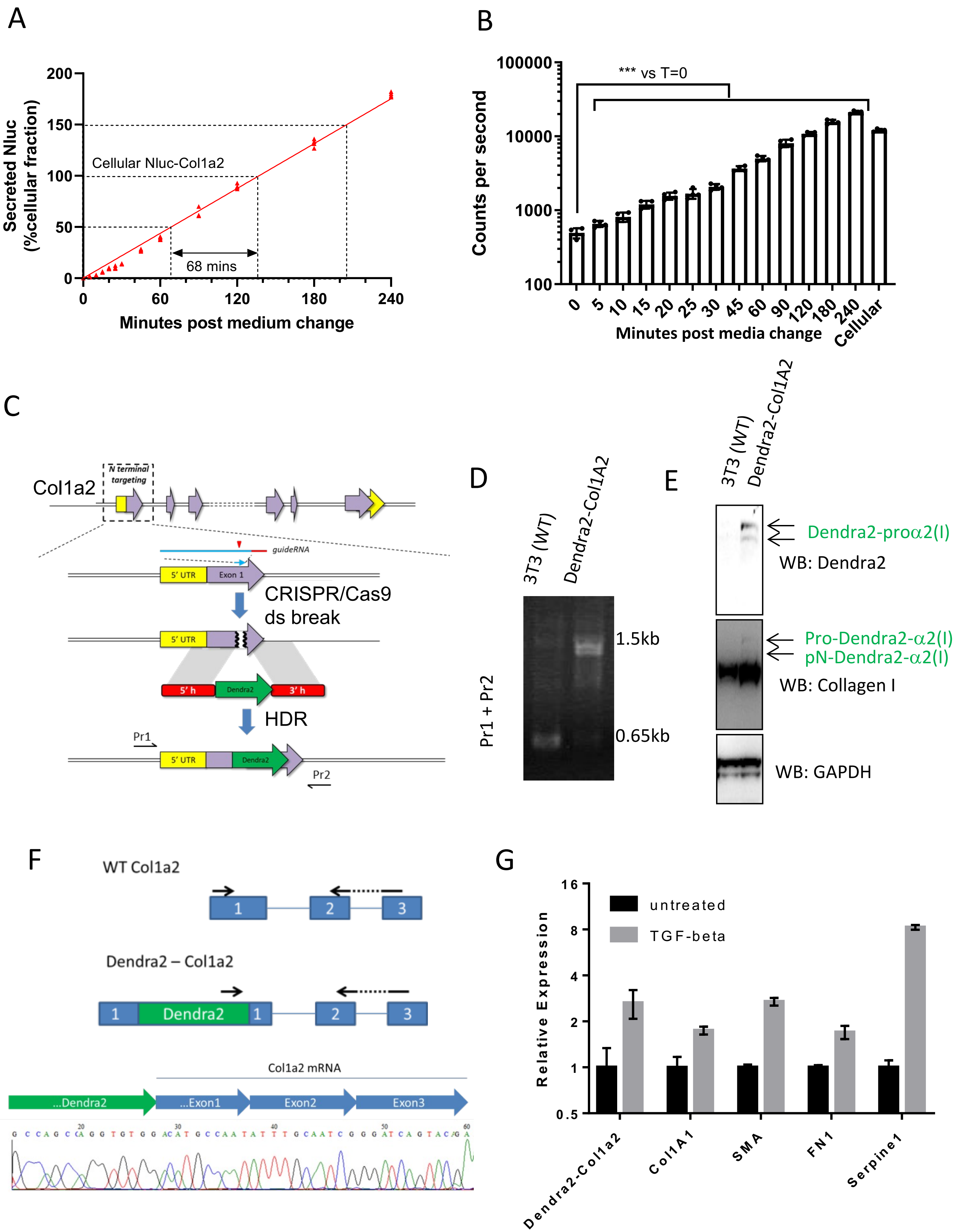

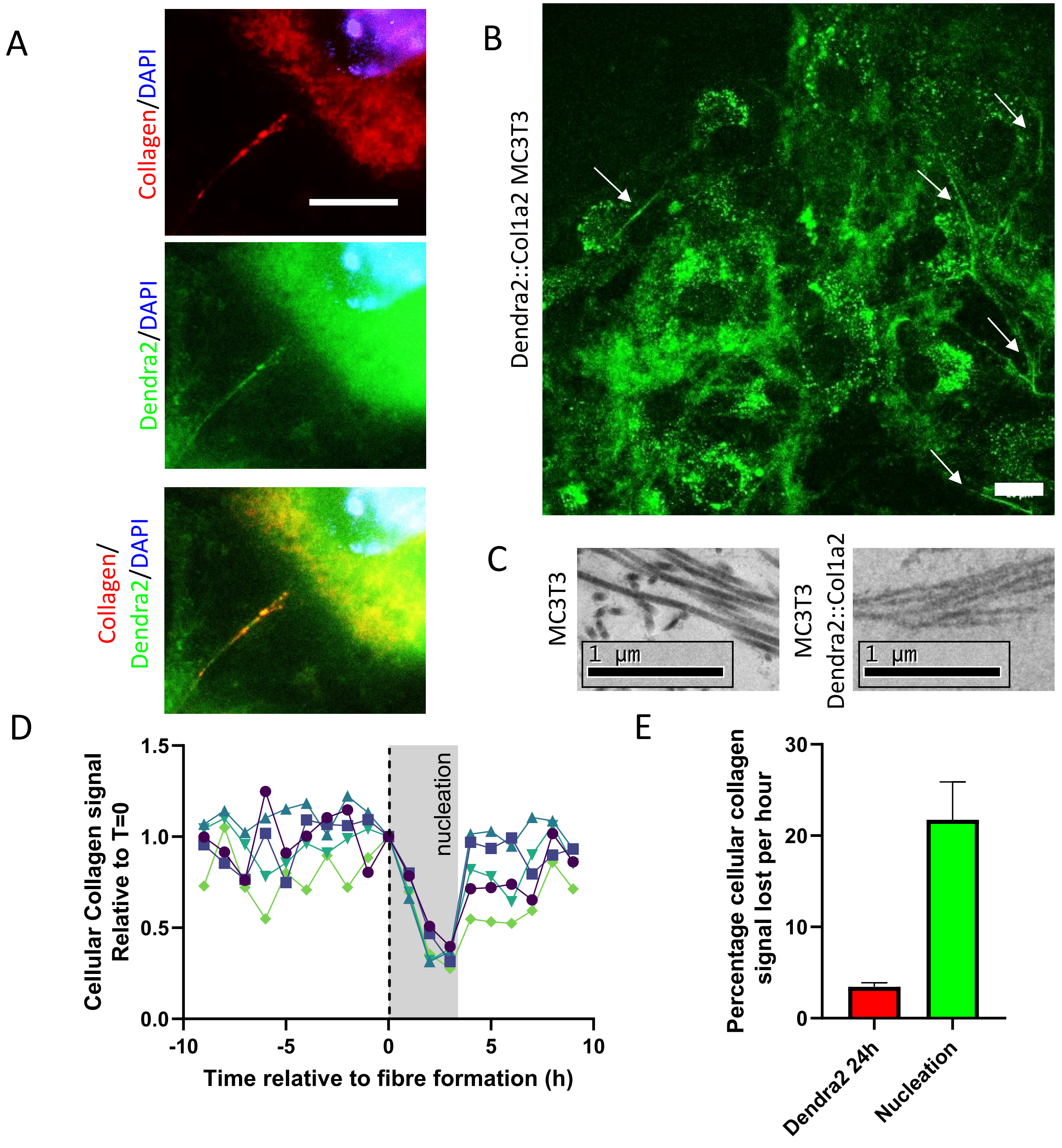

**Figure S3 Pickard *et al.***

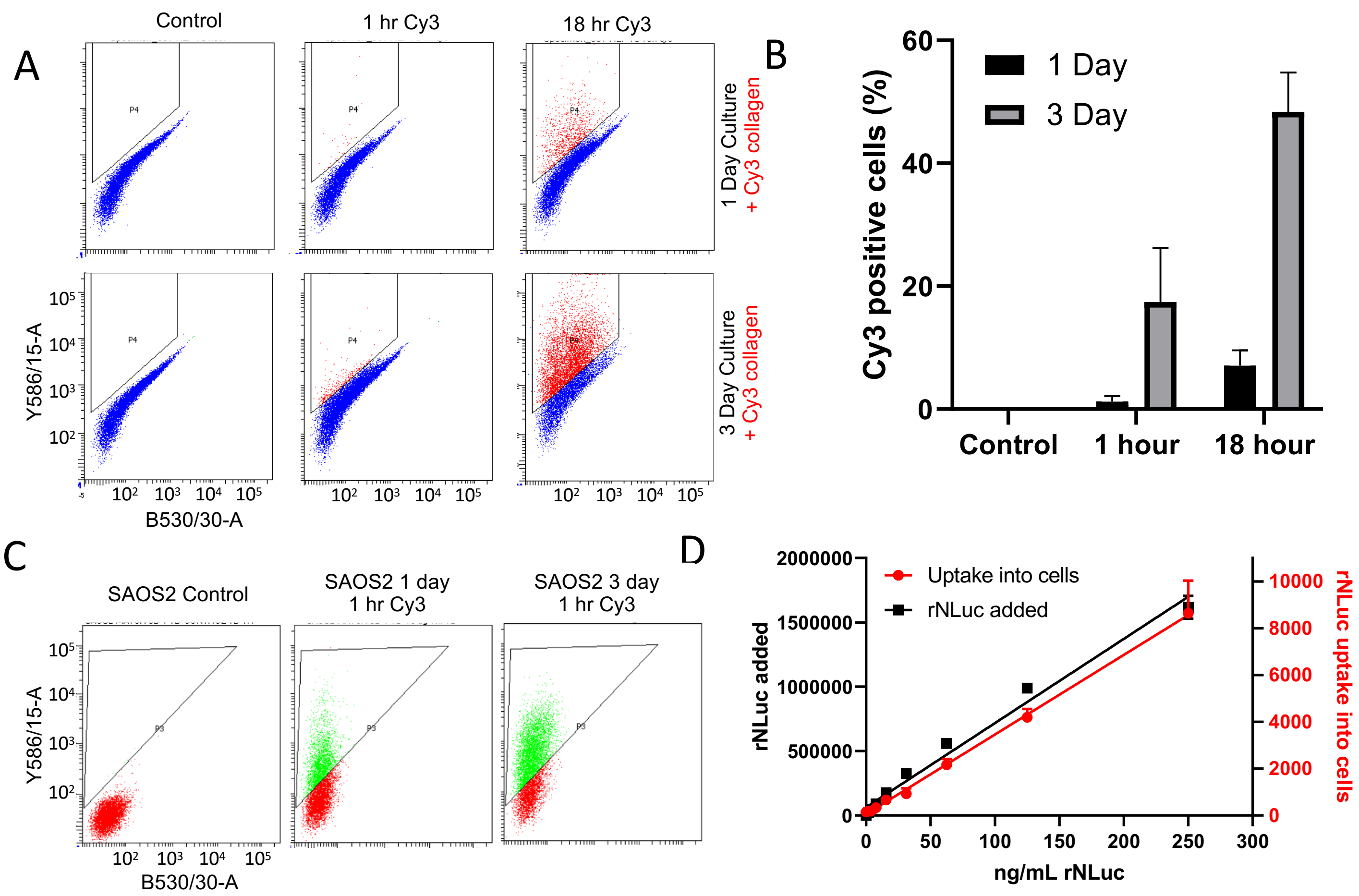

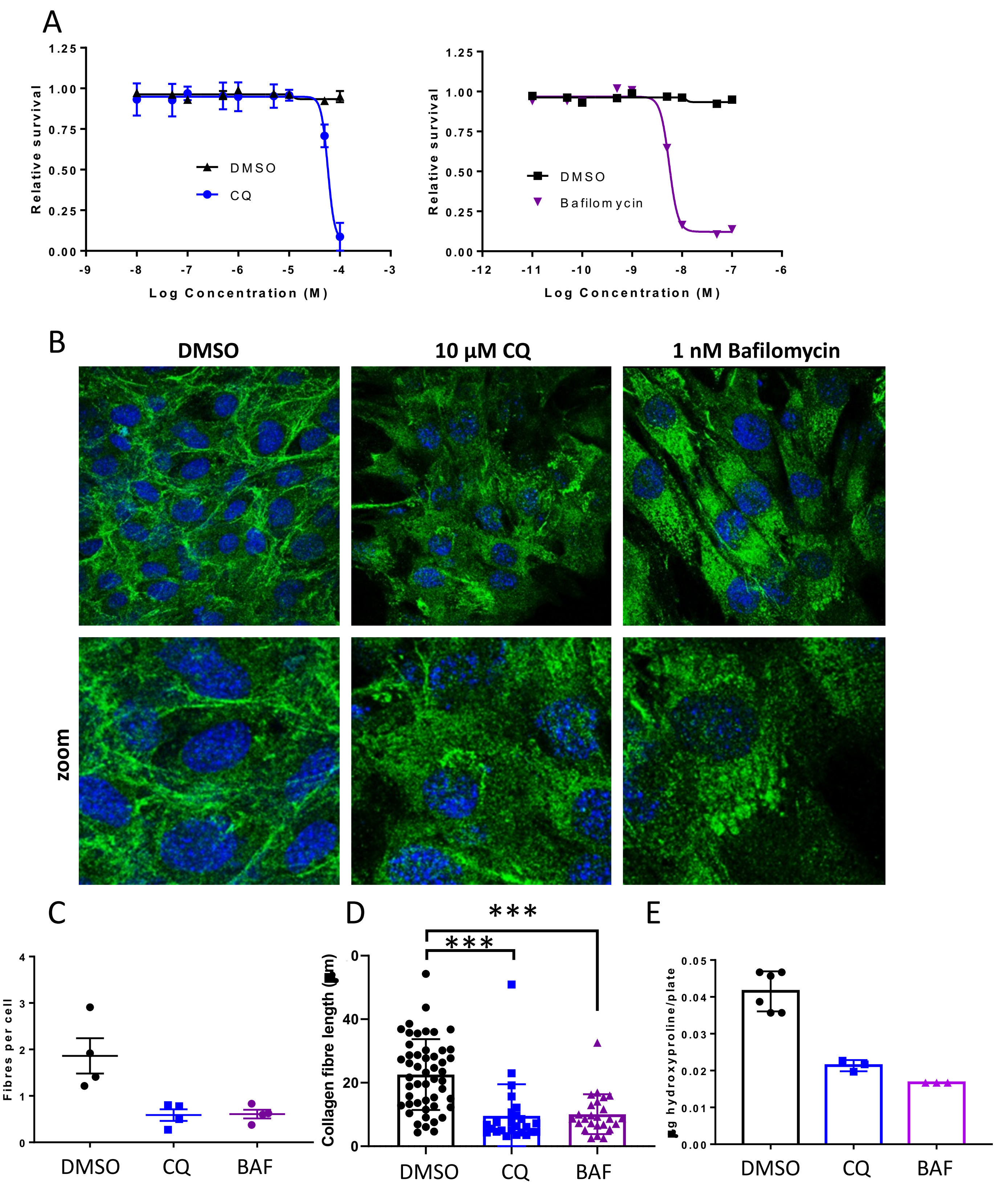

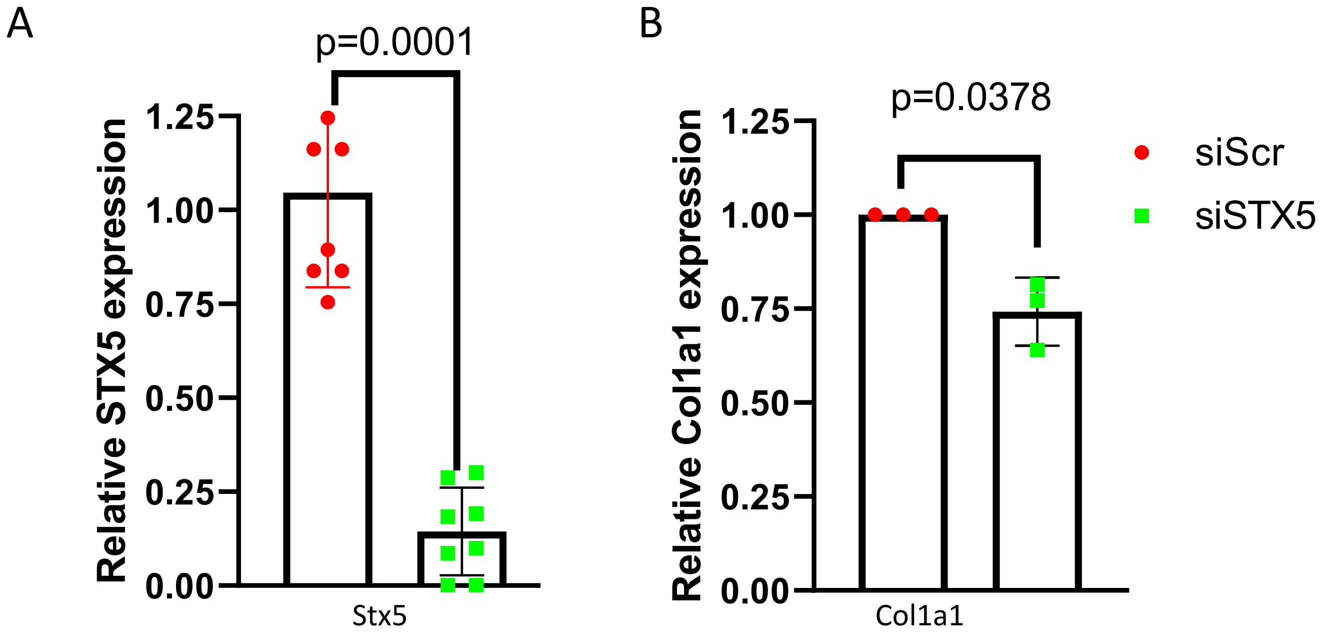

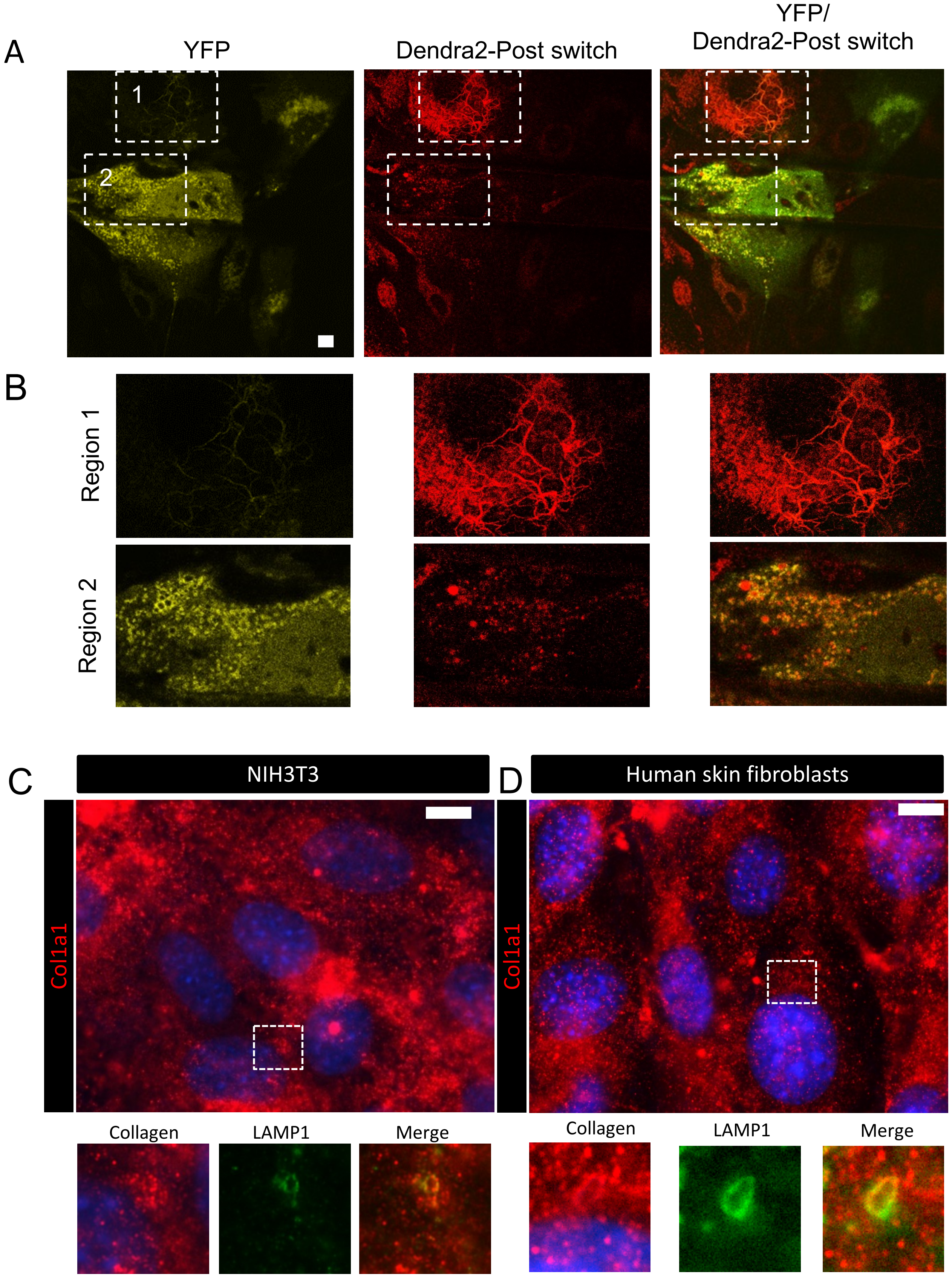

**Figure S7 Pickard *et al.***

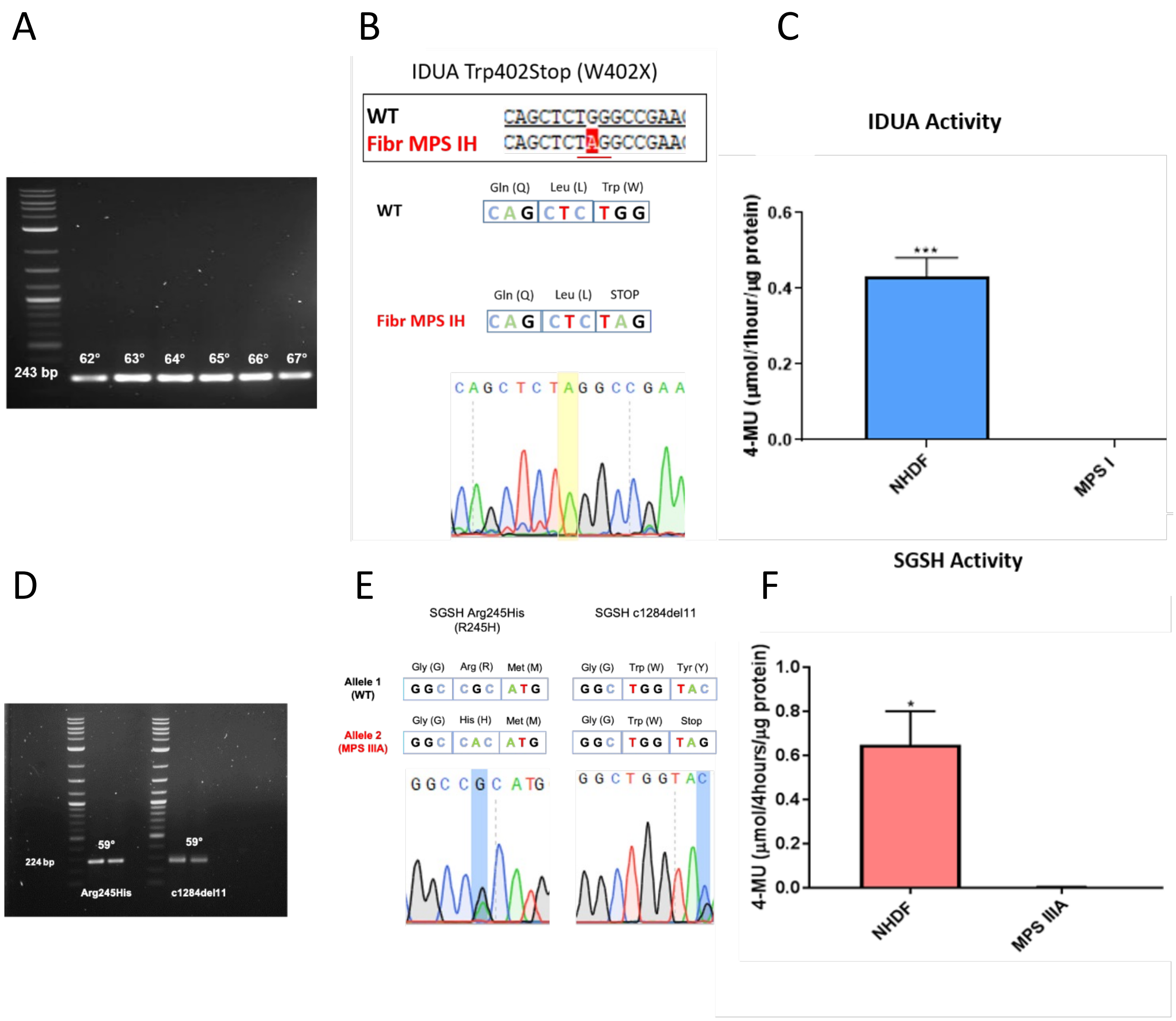
